# SEPALLATA MADS transcription factors act as key regulators in fertilization efficiency, ovule outer integument growth and mucilage secretory cell differentiation in Arabidopsis

**DOI:** 10.64898/2026.08.20.745741

**Authors:** Aline Janeau, Léa Rambaud-Lavigne, Nicola Babolin, Michel Paul, Anaïs Michaud, Lorène Masson, Jérémy Lucas, Charlie Scutt, François Parcy, Lucia Colombo, Chloé Zubieta, Véronique Hugouvieux

## Abstract

In angiosperms, ovule development requires the activity of the C, D and E classes of MADS genes, which encode key transcriptional regulators of reproductive development. The SEPALLATA (SEP) MADS transcription factors (MTFs), which belong to the E class, act as organizing hubs of MADS heterotetrameric complexes and play an essential role in the development of flower organs. However, the role of the *SEP* genes in ovule and seed development has been difficult to determine due to redundancy in the subclade, the lack of observable phenotypes in single and double *sep1 sep2* mutants and the homeotic conversion of the carpel into sepal or leaf in higher order *sep* mutants. Here, we engineered a version of *SEP3* (*SEP3^ΔM^*) that encodes a protein lacking the DNA-binding MADS-domain but retains the oligomerization domains needed for MADS protein heterotetramerization. *In vitro* experiments demonstrated the ability of SEP3^ΔM^ to interact with the C and D classes of MTF, reducing the capability of such MADS complex to efficiently bind DNA. *sep3^ΔM^* plants showed a delay in flower opening and organ maturation and a reduced fertility. The ovules exhibited reduced outer integument growth, and the few seeds that developed showed impaired mucilage secretion upon imbibition. RNA-seq analysis of *sep3^ΔM^* demonstrated misregulation of genes involved in outer integument and seed coat development. Taken together, these data indicate the key role of SEP3-containing MADS complexes in proper ovule outer integument growth and seed coat development.

## Introduction

The MADS-box genes are master regulators of plant development, orchestrating diverse biological processes that shape the architecture of reproductive structures and organs in all land plants (Shindo et al. 1999; Batman et al., 2022; Becker et al. 2000, 2003; Dreni and Ferrándiz 2022). The MADS-box genes expressed in the flower organs of angiosperms are the most studied and are divided into five main classes - A, B, C, D and E. The overlapping expression patterns of these genes determines the identity of each of the floral organs (Bowman et al. 1991; Coen and Meyerowitz 1991; Pelaz et al. 2000; Favaro et al. 2003; Pinyopich et al. 2003; Ditta et al. 2004). In Arabidopsis, the A class consists of *APETALA1* (*AP1*) and the non-MADS-box gene *APETALA2 (AP2)*, which in combination with E MADS genes confer sepal identity. The B class genes, *PISTILLATA* (*PI*) and *APETALA3* (*AP3*), in conjunction with A and E genes, confer petal identity. The C class gene, *AGAMOUS* (*AG*), along with B and E, confers stamen identity and the combination of C and E determines carpel identity. The D class genes, members of the *AG* clade, include *SEEDSTICK* (*STK*), *SHATTERPROOF1* and *2* (*SHP1/2*) and specify ovule identity together with the C and E classes. The E class contains four closely related and highly redundant genes, *SEPALLATA1-4 (SEP1-4)*, which are involved in the development of all flower organs and are required for the formation of transcriptionally active MADS protein complexes. With the exception of A class function, which is poorly conserved outside the Brassicaceae family, the B-, C-, D-, and E-class framework represents a widely applicable model for flower development, including ovule and seed development (Litt and Kramer 2010; Heijmans et al. 2012).

At the protein level, the MADS genes described above encode for MADS transcription factors (MTFs) from the type II MIKC^C^ subgroup (De Bodt et al. 2003; Parenicová et al. 2003; Lai et al. 2019). MIKC^C^ have a four domain structure consisting of a ∼50-60 amino acid (aa) <u>M</u>ADS DNA-binding domain (M-domain), a short ∼20-30 aa dimerization Intervening domain (I-domain), a dimerization/tetramerization <u>K</u>eratin-like domain (K-domain), as well as a largely unstructured <u>C</u>arboxyl-terminal domain (C-domain), implicated in transactivation and potentially in higher order complex formation (Kaufmann et al. 2005b). MTFs directly interact with members of the subgroup, forming homo- and heterodimers and tetramers. Based on genetic, biochemical and structural studies, tetramer formation is required for MTF physiological function, and the identity of the MADS proteins forming the tetramer determines the organ identity in the corresponding floral whorls (Pelaz et al. 2000; Honma and Goto 2001; Theissen and Saedler 2001; Favaro et al., 2003; Hugouvieux et al. 2024). In angiosperms, the SEPs act as central organizing hubs of MTF heterotetrameric complexes and are required for MTF tetramer formation, resulting in the *in vivo* transcriptional activity of MTFs during floral organ development (Theissen and Saedler 2001; Immink et al. 2009; Melzer et al. 2009; Mendes et al. 2013; Hugouvieux et al. 2024).

The roles of MTFs in ovules and seeds are less well studied than their function in floral organ determination. In flowering plants, the female floral whorl, or gynoecium, contains one or more organs termed carpels. Carpels may occur as free structures, or be fused longitudinally into a syncarpous pistil. The pistil, or each free carpel in apocarpous species, is typically differentiated into an apical stigma, a style, and a basal ovary that contains the ovules. In Arabidopsis and most other flowering plants, the ovule is composed of a nucellus, in which a Megaspore Mother Cell (MMC) undergoes meiosis to generate four megaspores, one of which is named the Functional Megaspore (FM) and will generate the female gametophyte by three mitotic division, a chalaza from which originates the integuments that enclose the nucellus, and a funiculus that attaches the ovule to the placenta to sustain ovule and seed development (Schneitz et al. 1995; Drews and Koltunow 2011). After double fertilization, the ovule in most angiosperm species develops into a seed containing a diploid embryo and a triploid endosperm, surrounded by a seed coat derived from the maternal integuments (Schneitz et al. 1995; Magnani 2018). The seed coat protects the developing embryo (Haughn and Chaudhury 2005). *AG* (C class), *STK* and *SHP1/2* (D class) genes form a monophyletic clade and are highly expressed in the placenta and ovule primordia (Pinyopich et al. 2003; Cucinotta et al. 2014). Triple *sep1^+/−^sep2 sep3* and *stk shp1 shp2* mutants develop carpeloid structures in place of ovules, while *ag* or *sep1 sep2 sep3* mutants fail to develop carpels. Furthermore, ectopic expression of *STK* or *SHP* genes in the first whorl, where *SEP* genes are expressed, converts sepals into carpeloid organs bearing ovules (Favaro et al. 2003). These studies suggest that a protein complex containing at least SEP, STK and SHP1/2 plays a central role in ovule development in *Arabidopsis* (Favaro et al. 2003; Pinyopich et al. 2003). Although SEP proteins have been proposed to form heterotetrameric MADS-domain complexes with C- and/or D-class factors to promote ovule development, direct evidence for their specific functions in ovules and seeds remains limited. In addition, while the role of *STK* has been well established through genetic studies, the identity of the MTFs in STK-containing MTF complexes has not been fully determined. *STK* has been shown to regulate several aspects of seed development, including seed coat mechanical properties through the control of mucilage composition, seed size, and seed abscission (Pinyopich et al. 2003; Mizzotti et al. 2014; Balanzà et al. 2016; Ezquer et al. 2016; Di Marzo et al. 2020; Paolo et al. 2021). More recently, STK was found to participate in MMC differentiation through its interaction with SPOROCYTELESS/NOZZLE (SPL/NZZ), leading to the proposition of a SEP–STK–SPL protein complex (Cavalleri et al. 2025). Furthermore, together with ABS, a B-sister MTF, STK has been shown to play critical roles in regulating maternal nutrient flow during ovule maturation and seed development (Banfi et al., 2026). The other D class MADS genes, *SHP1/2,* play roles in aspects of seed development as the *shp1 shp2* double mutant exhibits a slight defect in mucilage production (Ehlers et al. 2016). In addition, genetic interactions between *SHP1/2* and *ABS*, as well as between *STK* and *ABS,* have been shown to regulate differentiation of the endothelium, the innermost layer of the inner integument (Mizzotti et al., 2012, Ehlers et al., 2016).

To examine SEP functions beyond their role in perianth, stamen and carpel organogenesis, we developed a genetic tool to selectively impair the DNA-binding ability of SEPs while preserving their protein-protein interaction potential. Using genome editing, we targeted the M-domain of SEP3, generating the *sep3^ΔM^*allele that encodes a version of SEP3 containing the I, K and C domains, capable of oligomerizing with MADS partners but unable to bind DNA. This allele acts to titrate the *in vivo* activity of SEPs and reveals novel functions of SEP3 in ovule and seed development. *In vitro* DNA-binding assays confirm both the loss of DNA-binding capacity and the ability of *sep3^ΔM^*to interact with MADS partners, AG and STK, while RNA-seq analyses reveal transcriptional deregulation of *SEP* related genes. Phenotypic and microscopic analyses of *sep3^ΔM^* mutants highlight new phenotypes specifically affecting outer integument growth and outer layer seed coat differentiation. In addition, our experiments suggest that beyond their well-defined roles in floral organ development, SEPs also act as central regulators in ovule development, seed set and seed viability. These data further support their role as the key hub in reproductive efficiency in angiosperms.

## Results

### Generation of *sep3^ΔM^* homozygous lines

The *SEP* MADS genes are highly redundant in *Arabidopsis*. However previous studies have shown that a single allele of *SEP3* is able to restore organ formation in *sep1 sep2 sep3*^+/−^ plants, whereas single copies of *SEP1* and *SEP2* were unable to restore organ identity (Favaro et al. 2003; Hugouvieux et al. 2024). Moreover, SEP3 interacts promiscuously with other MADS-domain proteins to form active tetrameric complexes (Immink et al. 2009). For this reason, we decided to delete the SEP3 DNA-binding domain using CRISPR-Cas9 genome editing. We designed two highly specific guide RNAs that targeted and removed the first exon encoding the M-domain (Figure 1A). Based on structural studies, dimerization and tetramerization of MTFs are known to be primarily mediated by the I and K domains (Puranik et al. 2014; Hugouvieux et al. 2024). Accordingly, in the newly generated mutant, the SEP3^ΔM^ protein was expected to lack DNA-binding capacity but to retain its ability to interact with other MTF partners, including floral organ A, B, C and D proteins. Two stable and fertile homozygous *sep3^ΔM^* lines carrying a deletion between the two guides but no longer expressing Cas9, were isolated from the T3 generation. Sequencing of the *SEP3* locus confirmed the deletion of the majority of the M-domain and the in-frame retention of the I, K and C domains in both independent lines (Figure S1A). Quantitative PCR (qPCR) analysis of flower buds from the homozygous *sep3^ΔM^* plants confirmed that *SEP3^ΔM^* was expressed at levels comparable to full-length *SEP3* in wild-type plants (Figure S1B, C). When introduced into the *sep1 sep2* background, the *sep3^ΔM^*mutation triggered loss of floral organ identity and flower indeterminacy typical of the triple *sep1 sep2 sep3* mutant (Pelaz et al. 2000; Hugouvieux et al. 2018, 2024) indicating that the mutant had lost SEP3 activity due to deletion of the M-domain (Figure S1D).

**Figure 1:**
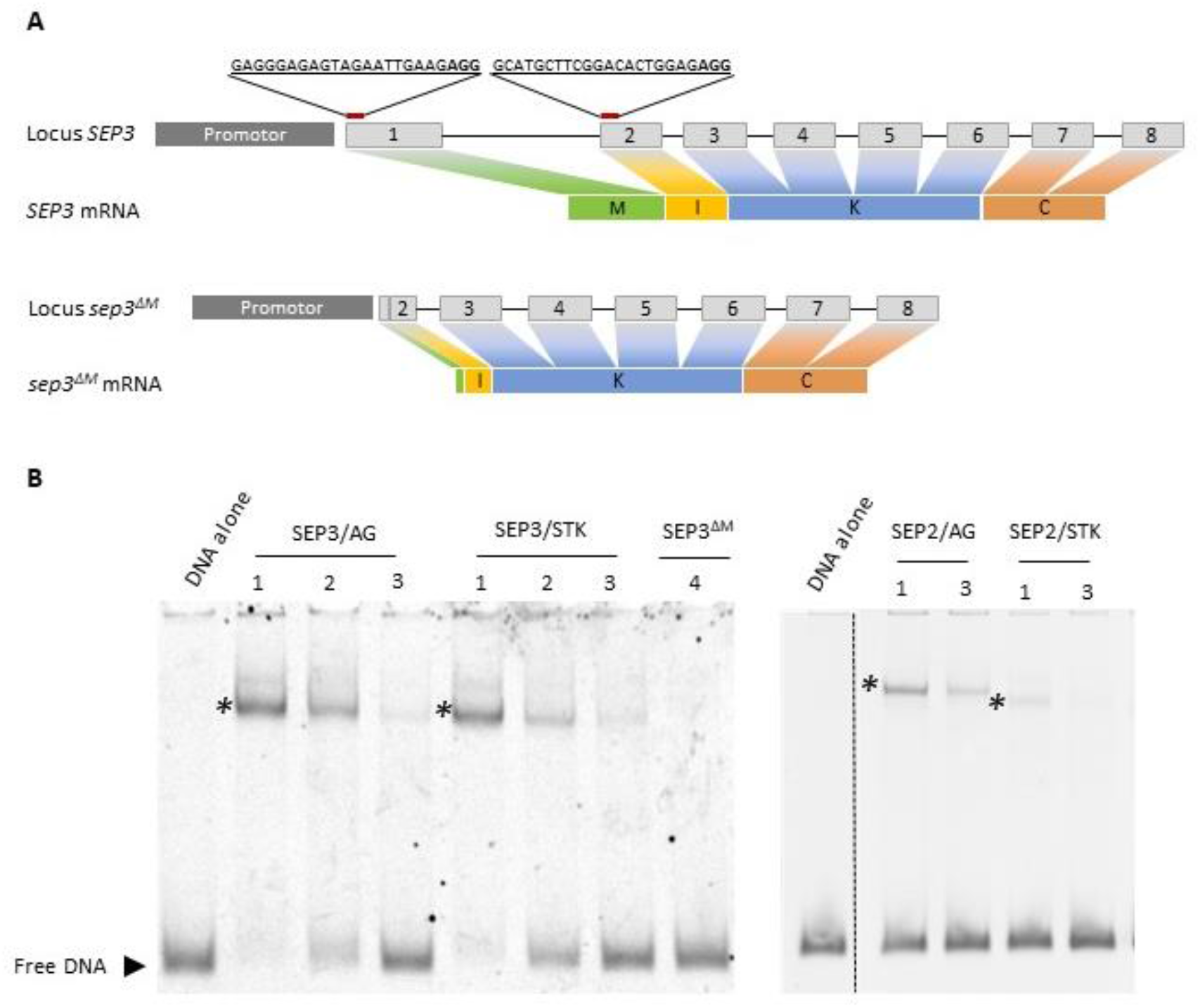
SEP3^ΔM^ protein decreases MADS complex DNA binding. **(A)** *SEP3* and *SEP3^ΔM^* loci, and position of guides RNA and sequences to generate *sep3^ΔM^* by CRISPER Cas9. Exons are represented by grey squares. M, I, K and C refer to the 4 domains of the SEP3 protein. **(B)** EMSA showing the ability of SEP3^ΔM^ to interact with MADS partners and to reduce DNA binding in the presence of *SEP* members. The DNA probe used is a 103-bp *SEP3* genomic fragment that contains two DNA binding sites (CArG-box). The control labelled “DNA alone” was run with the *in vitro* translation assay with pSP64 empty vector and incubated with the DNA probe. Protein production was performed with 0.25µg of each plasmid encoding SEP3, AG and STK, with 0, 0.5 and 1µg of plasmid encoding SEP3^ΔM^ in lanes 1, 2 and 3 respectively. Protein production was performed with 1µg of *SEP3^ΔM^* plasmid alone in lane 4. *\**: 4 MADS bound to DNA. The vertical black dashed line indicates a cut in the gel.

### SEP3^ΔM^ interacts with MTFs in vitro and reduces MADS DNA binding

We next investigated whether SEP3^ΔM^ retains the ability to interact with its MTFs partners and the impact of the loss of its M-domain on the DNA-binding capacity of any MADS complexes formed, using electrophoretic mobility shift assays (EMSA). A fluorescent labeled DNA probe containing two CArG-box MADS binding sites was used to examine SEP3^ΔM^ interactions and DNA binding with AG and STK, two key MTFs involved in carpel, ovule and seed development (STK) (Favaro et al. 2003; Pinyopich et al. 2003; Ezquer et al. 2016). EMSA demonstrated that SEP3^ΔM^ alone was unable to bind DNA (Figure 1B). However, it retained the ability to interact with AG and STK, competing with full-length SEP3 or SEP2, another member of the family, and resulting in reduced DNA-binding (Figure 1B).

### *sep3^ΔM^* mutant plants present delayed flower and silique development

Flower development in *sep3^ΔM^* plants was examined and compared with WT and the *sep3* loss-of-function mutant, referred to below as *sep3* (Hugouvieux et al. 2024). As shown in Figure 2A, *sep3^ΔM^*flowers exhibited mild developmental defects relative to both WT and *sep3*, including a delay in flower opening time and floral organ maturation (Figure 2A). In WT and *sep3* mutants, flowers opened at stage 13 based on (Smyth et al. 1990) (Figure 2A, column 1) and displayed fully expanded petals, as well as well-developed stamens and carpels, with **s**tigmatic cells abundantly covered with pollen grains (Figure 2A, columns 2-3). At this stage, WT stamens and carpels had similar lengths, as in *sep3* (Figure 2B), allowing an optimal deposition of pollen on stigma. At the corresponding developmental time determined by the position of the flower on the stem, *sep3^ΔM^* flowers remained at the pre-anthesis stage, unopened, with immature stamens and carpels (Figure 2A, column 1-2). When the flower buds opened, petals were incompletely expanded, with the apical margins frequently remaining partially folded (Figure 2A, column 3). Stamens were significantly shorter than carpels (Figure 2B) and pollen deposition on stigmatic cells was frequently reduced compared to WT (Figure 2A, columns 2, 3; Figure 2C). At stage 17 (Smyth et al. 1990) while silique elongation was significant in the WT and *sep3*, silique development was still not visible in *sep3^ΔM^* plants at the equivalent stage (data not shown). At maturity, silique length in *sep3^ΔM^* was reduced by approximately 40–50% (p<0.001) compared to WT (Figure 2D). In homozygous *sep3^ΔM^* expressing *pSEP3::SEP3* (Figure 2C, D), flower development and silique length were completely rescued and these plants looked phenotypically like the WT plants indicating that the observed phenotypes were not due to off targets. In addition, T1 *sep3* plants expressing *SEP3*^ΔM^ as a transgene under the *SEP3* promoter showed similar flower growth defects and silique reduced length to those of *sep3^ΔM^*(Figure 2C, D, S2A, B). None of these phenotypes were observed on T1 WT plants expressing *SEP3*^ΔM^ (Figure S2A, B) indicating a semi-dominant effect of *SEP3*^ΔM^. All together these data demonstrate that *sep3^ΔM^*show enhanced floral organ and silique development defects compared to *sep3*, linked to *SEP3*^ΔM^ expression.

**Figure 2:**
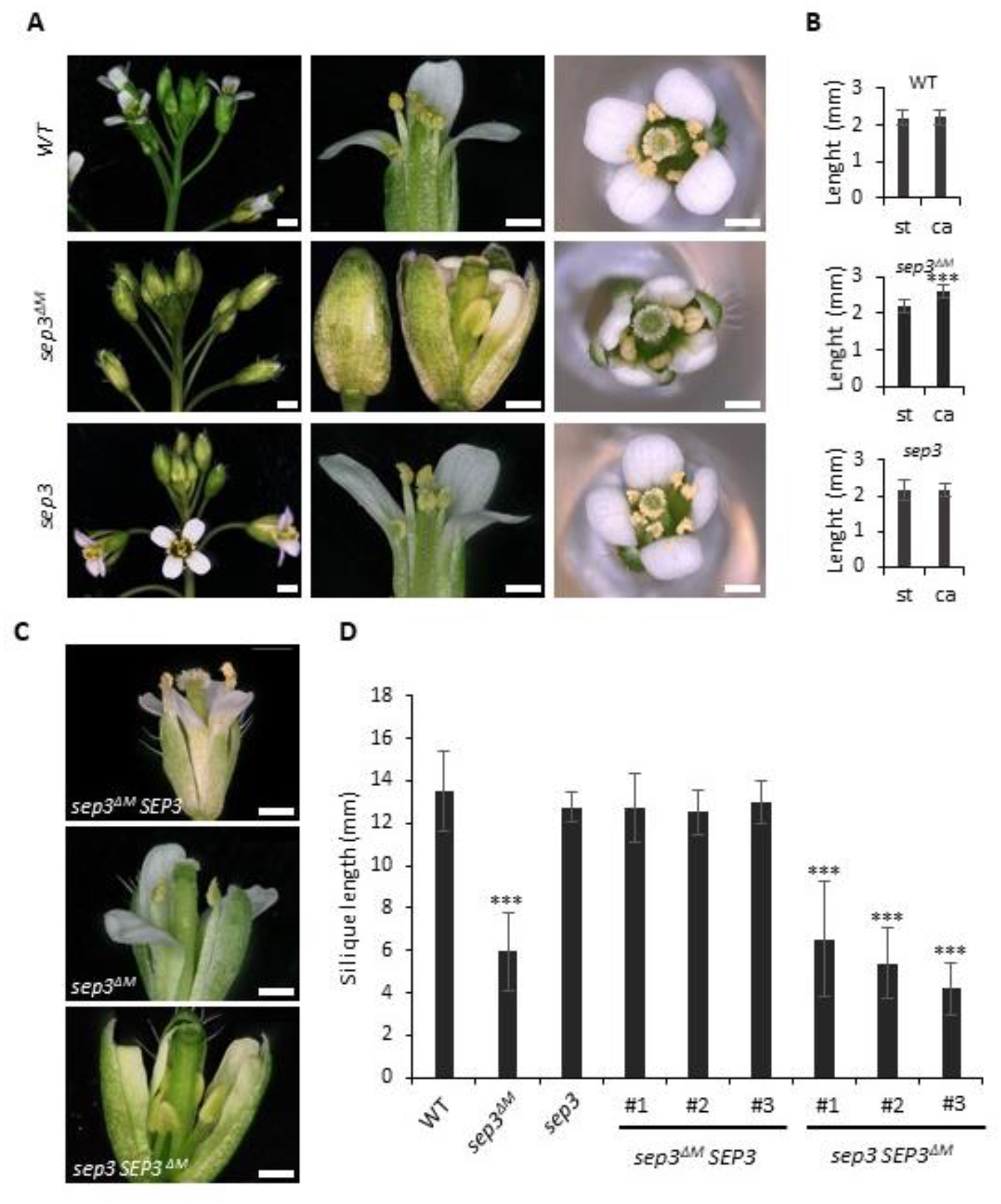
Floral organs and silique development is affected in *sep3*^ΔM^ mutant. (A) Comparative inflorescence (first column) and flower development (second and third columns) in WT, *sep3^ΔM^* and *sep3.* Second column indicates a flower at anthesis in WT and *sep3*, and at the corresponding time of development in s*ep3^ΔM^.* The unopened flower *sep3^ΔM^* is shown before and after manual opening. Third column shows a top view of a flower just opened in each background. Scales: 1 mm in column 1 and 500µm in columns 2 and 3. **(B)** Stamens and carpel length measured in WT, *sep3* and *sep3^ΔM^* first opened flowers. Data represent the mean ± SD. *st:* stamens; *ca*: carpel. Asterisks indicate significant differences between stamens and carpel length (\*\*\**P <* 0.001). No significant difference was measured between WT and *sep3*. **(C)** Representative image of the first opened flower in s*ep3^ΔM^*, *sep3^ΔM^* complemented lines expressing *SEP3* (*sep3^ΔM^* SEP3), and T1 *sep3* expressing *SEP3*^ΔM^ as transgene (*sep3 SEP3^ΔM^*). Scales: 500µm. **(D)** Silique length at maturity in WT, *sep3^ΔM^, sep3,* 3 homozygous *sep3^ΔM^*complemented lines expressing *SEP3* (*sep3^ΔM^* SEP3), and 3 T1 *sep3* expressing *SEP3^ΔM^* as transgene (*sep3 SEP3^ΔM^*). Data represent the mean ± SD. Asterisks indicate significant differences compared to WT (\*\*\**P <* 0.001). Data from 10 T1 *sep3 SEP3^ΔM^* in comparison with 10 T1 WT *SEP3^ΔM^* are shown in Figure S2.

### SEP3^ΔM^ does not properly regulate *SEP* target genes expression

RNA-seq analyses were performed on stage 10–11 flower buds from WT Col-0 and *sep3^ΔM^* plants and compared with previously published RNA-seq data obtained from *sep1 sep2 sep3* triple mutants at the same developmental stage (Lai et al. 2020). Three hundred and sixty genes were deregulated in *sep3^ΔM^* (−1 < log FC > 1, FDR < 0.05) compared to WT, versus 2358 in the *sep1 sep2 sep3* triple mutant, with the majority of them being downregulated (Figure S3; Table SI). Among the 360 genes deregulated in *sep3*^ΔM^, 56% (203) were common with deregulated genes in the *sep1 sep2 sep3* mutant (Table S1), however with lower fold changes correlating with a milder phenotype of *sep3^ΔM^* compared to the *sep1 sep2 sep3* mutant. Together, these results indicate that SEP3^ΔM^ selectively disrupts a subset of SEP-dependent regulatory networks *in vivo*.

### The *sep3^ΔM^* mutant exhibits a reduced fertilization rate and maternal inherited seed development defects

Experiments described above confirmed that *SEP3^ΔM^* impairs *SEP*-related pathways *in vivo*. We therefore pursued detailed examination of silique development. Seeds developed in *sep3^ΔM^* siliques but on average, 44% of ovules remained unfertilized and 11% of seeds were aborted (Figure 3A, B). WT fertilization rates and amount of seed development were nearly fully restored in *sep3^ΔM^* plants expressing *SEP3* under its native promoter (Figure 3B) while few seed developed per silique in *sep3* expressing *SEP3^ΔM^* (Figure S2C). In addition, seed morphology showed pronounced differences between WT, *sep3* and *sep3^ΔM^* plants (Figure 3C, D). While WT and *sep3* seeds displayed a uniform oval shape with a length to width ratio of 1.55 ± 0.17 and 1.31 ± 0.21, respectively, 95% of *sep3^ΔM^* seeds named class 1, exhibited a distinct heart-shaped phenotype (Figure 3C, D), characterized by a decreased ratio of 0.95 ± 0.25. Within this set of seeds, about 50% appeared to be not fully closed at the micropyle region. Stronger morphological defects were observed in roughly 5% of seeds, named class 2, that show embryo extrusion at the micropyle site (Figure 3C, D). The extent of this class 2 phenotype varies among seed sets, showing slight or extensive internal tissue exposure. Despite these abnormalities, all but the most malformed class 2 seeds were able to germinate. Seed morphology was well complemented in homozygous *sep3^ΔM^* expressing *pSEP3::SEP3* (Figure 3D). In addition, T1 *sep3* plants expressing *SEP3^ΔM^* as a transgene under the *SEP3* promoter reproduced the abnormal seed phenotype of *sep3^ΔM^* plants (Figure 3D; Figure S2C, D). Seed phenotypes were not observed in WT expressing *SEP3^ΔM^* (Figure S2D). The complementation experiment highlighted the sporophytic origin of the mutant seed phenotype. Indeed, T1 *sep3^ΔM^* plants containing *pSEP3::SEP3* generated 100% of seeds with a WT phenotype, including the 25% transformants that are homozygous for the *SEP3^ΔM^* locus (Figure S4A). Reciprocal crosses between WT and *sep3^ΔM^* plants (Figure S4B) also confirmed the maternal origin of the seed phenotype since WT ovules fertilized with *sep3^ΔM^* pollen produced normal, oval-shaped seeds, whereas *sep3^ΔM^* ovules fertilized with WT pollen yielded heart-shaped seeds (Figure S4B). Of note, silique elongation measurements and seed content of reciprocal crosses (Figure S5A, B) suggested a reduced fitness of the *sep3^ΔM^* pollen compared to WT. In addition, ovule fertilization efficiency in the mutant was still reduced even when fertilized with WT pollen (Figure S5A and B, right panels). Collectively, these findings demonstrate that the impaired ovule fertilization efficiency observed in *sep3^ΔM^* plants might result from both reduced pollen fitness and female reproductive structure defect, whereas seed morphogenesis defects were maternal inherited.

**Figure 3:**
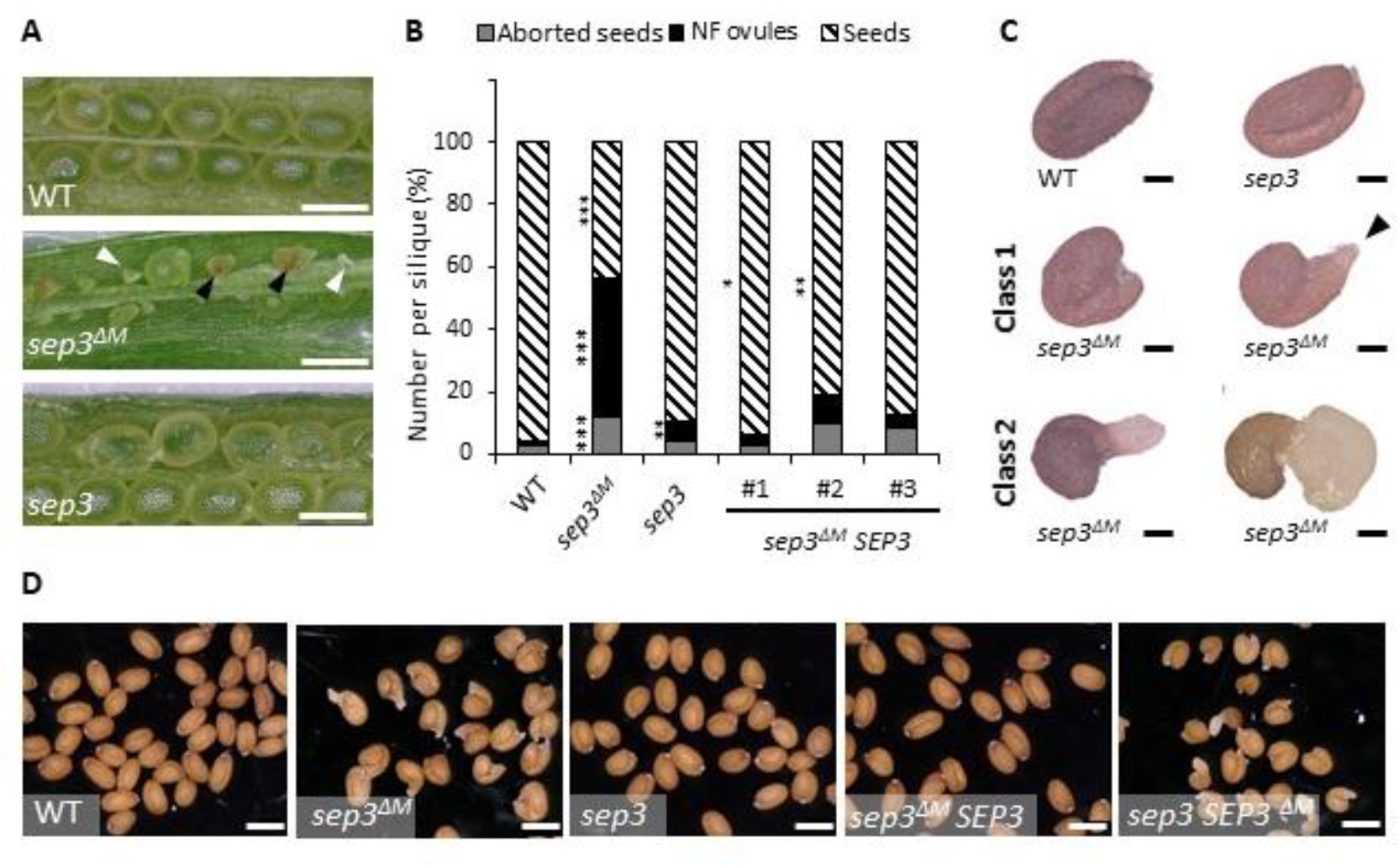
Fertilization rate and seed development is affected in *sep3*^ΔM^ mutant. **(A)** Opened mature silique in WT, s*ep3^ΔM^* and *sep3* highlighting unfertilized ovules (white arrow) and aborted seeds (black arrow) in s*ep3^ΔM^*. **(B)** Quantification of unfertilized ovules, aborted seeds and developed seeds in WT, s*ep3^ΔM^, sep3* and *sep3^ΔM^* expressing *SEP3* (*sep3^ΔM^ SEP3*). Asterisks indicate significant differences for each specie from WT (* *P <* 0.05, \*\**P <* 0.01, \*\*\**P <* 0.001). NF: non fertilized. **(C)** Zoom on seeds in WT, *sep3* and s*ep3^ΔM^* after harvest. 95% of the seeds present a heart shape (class 1), while 5% of the seeds show extrusion of the embryo (Class 2). Black arrowhead points to a slight extrusion at the micropyle region of 50% of class 1 seeds. **(D)** Representative seeds in WT, *sep3^ΔM^*, *sep3*, *sep3^ΔM^* complemented lines expressing *SEP3* (*sep3^ΔM^ SEP3*), and *sep3* expressing *SEP3^ΔM^* as transgene (*sep3 SEP3^ΔM^*). Scales: 500µm in A and D, 100µm in C.

### The *sep3^ΔM^* mutant exhibits defects in ovule integument development and seed coat differentiation

Since the abnormal seed development of the *sep3^ΔM^* mutant was maternally inherited, we examined integument development and seed coat differentiation after fertilization. Confocal analysis confirmed that in WT and *sep3* ovules, the outer integument (oi) overgrows the inner integument (ii), enclosing the embryo sac within the ovule at anthesis (Figure 4A) (Vijayan et al. 2021). The integuments usually consist of two layers of oi (oi1 and oi2) and two to three cell layers of ii (ii1, ii1’ and ii2) (Beeckman et al. 2000). In contrast, the majority of *sep3*^ΔM^ ovules exhibited obvious defective oi growth compared with WT and *sep3* (Figure 4A). Among 95 ovules analyzed in the mutant, 66% displayed reduced oi growth leading to a reduction in *sep3^ΔM^*enclosure of the embryo sac. The other 34% resembled WT, although the oi appeared shorter than in WT. Reduction in oi growth was also observed at anthesis in *sep1 sep3* or *sep2 sep3* double mutants but not in the *sep1 sep2* double mutant (Figure S6). In *sep3^ΔM^* complemented with *SEP3*, the growth of integument layers and enclosure of the embryo sac were comparable to WT (Figure S7). We next analyzed seed development at three days after pollination (DAP), when the embryo had reached the globular stage and differentiation of the seed coat was clearly visible. As expected, seed coat differentiation was evident in WT, *sep3* and *sep3^ΔM^ SEP3* lines with five distinct layers with the ii1′ layer, originating from the ii1 layer, being highly vacuolated (Figure 4B; Figure S7) (Schneitz et al. 1995; Coen et al. 2017). At this time, the endosperm nuclei were visible in the three genotypes, and developing seeds had already acquired an oval morphology in WT and *sep3* and complemented lines (Figure 4B; Figure S7). In contrast, in *sep3^ΔM^* (Figure 4B) the two outer layers oi1 and oi2 appeared thinner than in WT, particularly in the region opposite the micropyle, where larger and irregularly shaped cells were observed. The ii1′ layer was barely visible in *sep3^ΔM^*. Among ten developing seeds analyzed in *sep3^ΔM^*, four exhibited a round shape with a closed micropyle, while in the other six, the embryo was partially extruded, indicating defective confinement within the developing seed coat that resembled an unenclosed embryo sac in some ovules. Embryo development appeared slightly delayed in *sep3^ΔM^* (Figure 4B). Together, these results demonstrate that *sep3^ΔM^* is impaired in ovule oi growth and seed coat differentiation.

**Figure 4:**
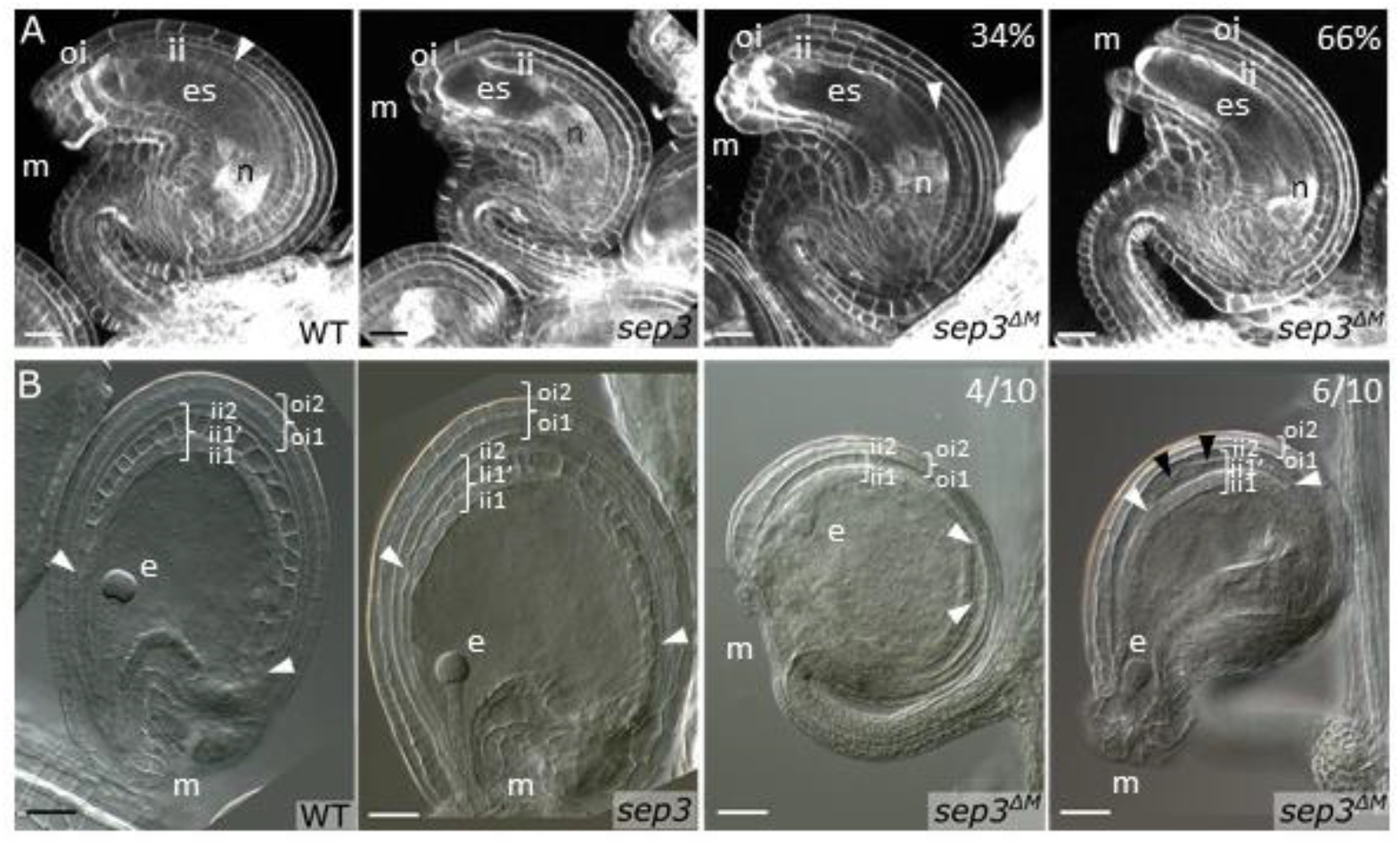
s*ep3^ΔM^* shows defects in ovule integument growth and seed coat differentiation. **(A)** Confocal analyses of WT (n=102), *sep3* (n=6) and s*ep3^ΔM^ (*n=95) ovules extracted from carpel on flower just opened. Pictures highlight outer integument growth defect in the s*ep3^ΔM^* mutant with 66% of the ovules showing shorter outer integument than inner integument leading to unclosed embryo-sac (s*ep3^ΔM^* right panel). The other 34% show however shorter outer integument growth than in WT (s*ep3^ΔM^* left panel). **(B)** Seed clearing 3 days after pollination of WT (n=4), *sep3* (n=8) and s*ep3^ΔM^* (n=10). The 5 layers of seed coat are differentiating in WT and *sep3*, with the ii1’ highly vacuolated. Reduced seed coat development is visible in *sep3^ΔM^*Layers appeared thinner and ii1’, when observed, was hardly visible. Seeds morphology in s*ep3^ΔM^* was already affected at 3 DAP. 6/10 seeds (*sep3^ΔM^* right panel) show growth of the developing embryo at the margin of the micropyle, while the other set have adopted a round shape. White arrows indicated ii1’ when visible, appearing from ii1 division. Black arrows indicate the appearance of an extra layer in *sep3^ΔM^. ii*: inner integument, *oi*: outer integument, *n*: nucellus, *m*: micropyle, *e*: embryo, es: embryo sac. Scales: 20µm in A and 40µm in B.

### *sep3^ΔM^* seeds exhibit undifferentiated mucilage secretory cells and defect in mucilage production

Both proanthocyanidins (PAs) and mucilage are major components of the *Arabidopsis* seed coat, that play distinct but complementary roles in seed protection (Haughn and Chaudhury 2005). While the inner most ii1 layer, the endothelium, accumulates PAs, the oi is specialized in mucilage production (Beeckman et al. 2000; Windsor et al. 2000; Debeaujon et al. 2003; Haughn and Chaudhury 2005). We used vanillin staining as a marker for the differentiation of the inner seed coat development, which stains the PAs produced in the endothelium after fertilization. As shown in Figure S8, three DAP PAs accumulation was similar in the WT, *sep3*, and *sep3^ΔM^* endothelium, though a slight reduction was detected in the chalazal and micropylar regions of *sep3* and *sep3^ΔM^* seeds that was restored back to WT in the *sep3^ΔM^ SEP3* complemented lines. We next examined the outermost seed coat layer (oi2), which consist of well-differentiated mucilage secretory cells (MSCs) in the WT seeds, by scanning electron microscopy (SEM) on dried seeds (Figure 5A, B). In WT seeds, MSCs display a polygonal shape with prominent radial cell walls, a central columella, and deposited mucilage (Haughn and Chaudhury 2005). In *sep3* and *sep3^ΔM^*-complemented lines expressing SEP3 (Figures 5 and S7C, D), MSCs appeared largely similar to those in WT, although slightly less uniform in shape. In all these genotypes, the columella structure, radial cell wall patterning, and mucilage deposition were clearly visible. In contrast, MSCs were poorly differentiated in *sep3^ΔM^* with the cells appearing larger and irregular, columella structures were mainly absent with depression in the center, and radial cell walls were poorly defined (Figure 5A, B). To assess mucilage production and release, seeds were subjected to water imbibition with or without EDTA treatment (Figure 5C, D). WT, *sep3* and *sep3^ΔM^*-complemented lines expressing SEP3 exhibited mucilage release after water imbibition, highlighted by the presence of a dark shadow in the MSC observed by SEM (Figure 5C), and confirmed by the presence of a red mucilage halo after staining with ruthenium red indicating mucilage extrusion (Figure 5D). Mucilage release was absent for *sep3^ΔM^*. EDTA treatment, a more stringent treatment for mucilage release, did not favor mucilage release in *sep3^ΔM^* (Figure 5D). The *stk* mutant, also characterized for mucilage release defect, showed classical architecture of MCS and mucilage released after EDTA treatment, but not after water imbibition (Figure S7) as previously described (Ezquer et al. 2016). Together, these observations demonstrate that SEP3^ΔM^ severely disrupts specifically the differentiation of MSCs leading to seed with an outermost layer of the seed coat lacking mucilage.

**Figure 5:**
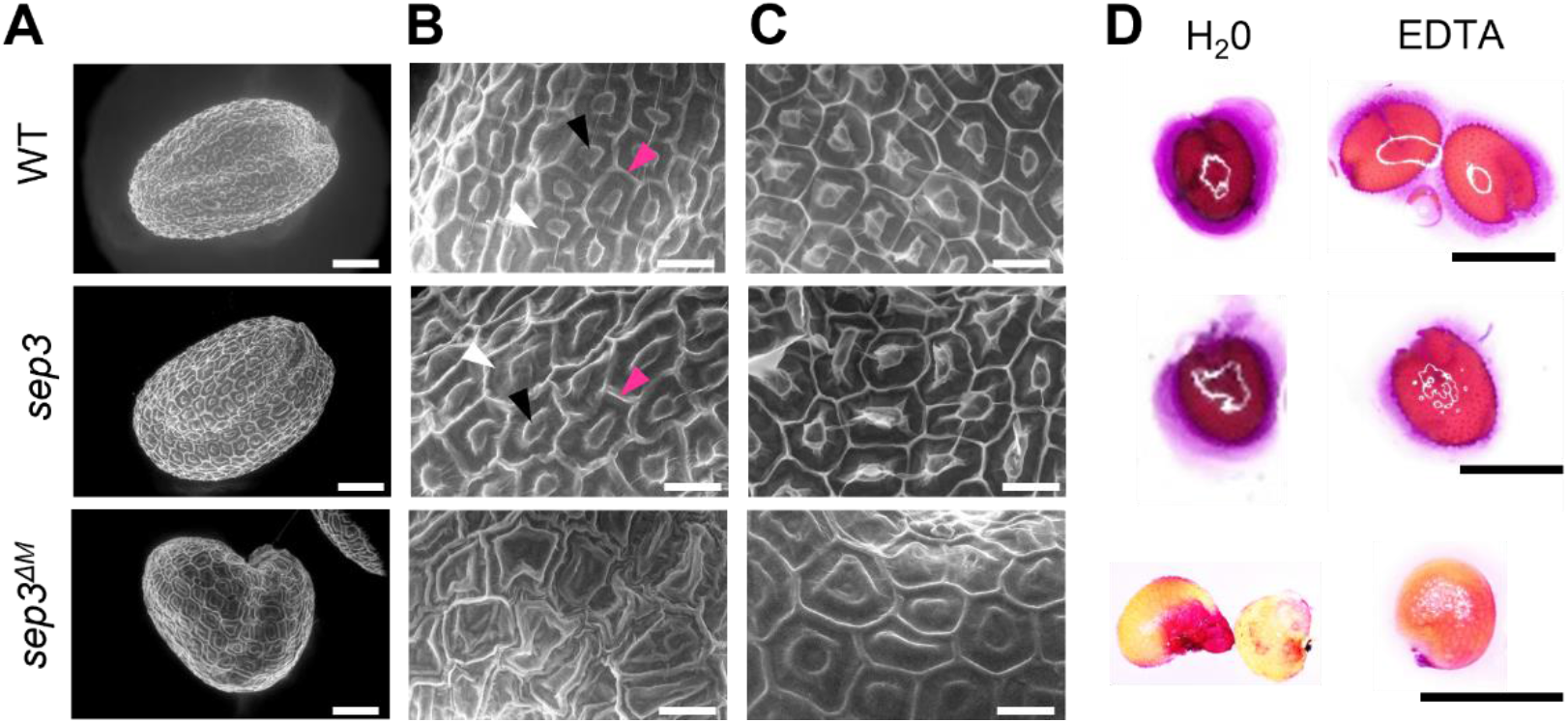
Impaired differentiation of MSC is observed in s*ep3^ΔM^* seeds. Representative scanning electron microscopy of WT, *sep3* and *sep3^ΔM^* performed on whole dried seeds **(A)**, dried cell surface **(B)** and after overnight water imbibition **(C)** highlight the abrogated MSC differentiation in s*ep3^ΔM^* and the lack of mucilage release in s*ep3^ΔM^* mutant. **(D)** Mucilage staining with ruthenium red after overnight imbibition and EDTA treatment in the various seeds genotypes. The absence of red staining after water and EDTA treatments confirm the lack of differentiation of MSC in s*ep3^ΔM^*. Black arrow indicate columella, pink arrow radial cell wall and white arrow mucilage. Scales: 100 µm in A, 300µm in B and C, and 500µm in D.

### Identification of potential molecular mechanisms

Based on the ovule phenotype described above, one likely downstream target affected in the s*ep3^ΔM^* mutant is *INO* (*INNER NO OUTER*), a member of the YABBY transcription factor family. *INO* is a key and specific regulator of oi development in *Arabidopsis* ovules (Villanueva et al. 1999; Skinner et al. 2023). Based on our RNA-seq analysis performed at a developmental stage at which ii and oi integuments initiate on the ovule primordia (Roeder and Yanofsky 2006), a reduction in *INO* expression (log FC = −1.2), was noted, although the associated FDR was close to the significance threshold. qPCR analysis, however, demonstrated that *INO* transcript levels were reduced by >80% in the mutant, linking ovule phenotype to *INO* expression (Figure S9). The three master regulators in MSCs differentiation include *APETALA2* (*AP2*), and two NAC TFs, *NAC-REGULATED SEED MORPHOLOGY (NARS)* 1 and 2 (Leon-Kloosterziel et al. 1994; Kunieda et al. 2008; Golz et al. 2018; Xu et al. 2023). Interestingly *NARS1* transcript levels were significantly reduced in the RNA-seq (log FC −2.3 FDR <0.03). Reduction in *NARS1* expression was further validated by qPCR (Figure S9). Together, these results suggest a link between ovule and seed phenotype in *sep3^ΔM^* to altered master regulators such as *INO* and *NARS1*.

## Discussion

In this work, we developed a genetic tool to probe the role of the *SEP* gene family beyond perianth, stamen and carpel organ identity. Because *SEP* genes act redundantly, functional analysis in different developmental contexts is challenging. Given the central role of SEP3 within the MADS family in flower and ovule development (Favaro et al. 2003; Hugouvieux et al. 2024) and the high level of *SEP3* expression in the ovule primordia and early stages of seed development (Figure S10; Mandel and Yanofsky 1998; Urbanus *et al*. 2009; Cavalleri *et al*. 2025; Martin et al., 2025) we generated a novel CRISPR-Cas9 *sep3* mutant expressing a SEP3 protein lacking the MADS DNA-binding domain while retaining its protein–protein interaction I and K domains. This strategy was designed to broadly reduce SEP complex DNA-binding activity across floral, ovule, and seed tissues. Characterization of *sep3^ΔM^* provides genetic evidence that *SEP3* functions not only in floral organ specification but also contributes to the successful fertilization process and outer layer development in angiosperm ovules and seeds.

The SEP3^ΔM^ protein was able to reproduce the interaction specificity of the full length SEP3 and SEP2 proteins based on *in vitro* titration experiments with AG and STK (Figure 1B). Indeed, structural data for SEP3 homotetramers and SEP3/AG heterotetramers have demonstrated that the K-domain alone is able to form tetrameric MADS complexes independently of other domains (Puranik et al. 2014; Hugouvieux et al. 2024). This mechanism of specific interaction with different MADS partners is likely recapitulated *in vivo* as more than 50% of the deregulated genes in *sep3^ΔM^* were also deregulated in the triple *sep* mutant, suggesting general perturbation of *sep*-related targets. The wide range of phenotypes observed in the *sep3^ΔM^* mutant, spanning defects in floral organ growth to seed development likely reflects the activity of SEP3^ΔM^ on multiple pathways. While reduced fertilization rate and increased seed abortion was never reported for single MADS mutant, these phenotypes are similar to those observed in the *SEEDSTICK ARABIDOPSIS BSISTER* (*stk abs*) double mutant (Mizzotti et al. 2012; Banfi 2025). Of note*, STK* and *ABS* have been shown to directly interact with SEP3 in DNA-binding assays and in yeast-2-hybrid experiments (Kaufmann et al. 2005a). Interestingly, the lack of PAs biosynthesis described in the a*bs* (Debeaujon et al. 2003) or *stk abs* (Mizzotti et al. 2012) mutants due to the lack of endothelium differentiation, was not observed in *sep3^ΔM^* highlighting that possibly a subset of ABS containing complexes are not significantly impaired in the s*ep3^ΔM^* mutant. One explanation could be that SEP3 is not the main SEP actor in PAs synthesis, but rather SEP1 or SEP2, as their expression was detected in the endothelium <u>(Martin et al. 2026;</u> Figure S10B). Both the *abs shp1 shp2* triple mutant and the *sep3^ΔM^* mutant display a similar reduction in endothelium divisions that give rise to the ii1′ layer (Ehlers et al. 2016). These findings suggest that distinct ABS-containing complexes may independently regulate PAs synthesis and endothelium division.

SEP3^ΔM^ may also affect the interaction of AG, SHP1, SHP2 and/or STK with the Zn-finger TF, NO TRANSMITTING TRACT (NTT) (Herrera-Ubaldo et al. 2018). The double mutant *stk ntt* showed a reduced fertilization rate due to transmitting track defects and abrogated pollen guidance (Herrera-Ubaldo et al. 2018). Interestingly, the co-repressors, SEUSS, and LEUNIG_HOMOLOG (LUH) are described to physically interact with SEP3 (Sridhar et al. 2006; Di Marzo et al. 2022a) and MSC lack of differentiation was observed in the double mutant *stk luh* (Di Marzo et al. 2022a). This raises the possibility that SEP3^ΔM^ might also impede recruitment of corepressors to their proper targets via the decrease in DNA-binding competent MADS complexes.

In WT plants, MSCs differentiate from the outer integument shortly after fertilization (Beeckman et al. 2000; Western et al. 2000). Confocal analysis of developing seeds at 3 DAP (Figure 4B) showed that s*ep3^ΔM^*embryos can initiate development from embryo sac that are not properly enclosed by the integuments, suggesting that the outer integument might not be fully differentiated in *sep3^ΔM^* for proper MSC initiation. In this context, reduced *INO* expression and the degree of outer integument growth at the time of fertilization is likely a key determinant of subsequent seed developmental outcome. Structural defects in *sep3^ΔM^* ovules at the micropyle region may directly impair fertilization as proper development of the outer integument is required for correct micropyle formation and accurate pollen tube guidance toward the embryo sac (Lora et al. 2019). Abnormal seed phenotypes observed in the *sep3^ΔM^* mutant, including seeds displaying tissue extrusion and seed abortion, is likely due to fertilization of ovules that exhibit varying degrees of outer integument growth defects. In addition to *INO*, *NARS1* transcript level was also reduced in the s*ep3^ΔM^*mutant (Figure S9) indicating that early stages of MSC differentiation might be strongly compromised in the *sep3^ΔM^* mutant. Lack of MSC differentiation was not observed in previously described *stk* (Ezquer et al. 2016) and *shp1 shp2* mutants (Ehlers et al. 2016), in which only slight mucilage composition defects were described and MSC architecture was preserved.

Globally, the *sep3^ΔM^* mutant recapitulates several phenotypes previously reported in two *Arabidopsis* mutants: the *ats* (*ABERANT TESTA)* mutant and the constitutive GA signaling *global* mutant (mutated in the 5 *DELLA* genes) (Gomez et al. 2016). These shared defects include not only reduced silique length, reduced fertility with nonfertilized ovules, but also round seeds shape, integument growth leading to incompletely enclosed ovules, and finally defects in MSC differentiation. In the Gomez et al. study, elevated GA signaling was proposed to negatively affect ovule development by restricting integument growth through the ATS–DELLA regulatory module. Interestingly, neither *ATS* nor *DELLAs* were deregulated in *sep3*^ΔM^ RNA-seq data. However, GIBBERELLIN 2-OXIDASE3, one of the key enzyme involved in GA catabolism (Rieu et al. 2008), was downregulated in *sep3^ΔM^* (Table S1; Figure S9), suggesting that GA levels may be elevated in the s*ep3^ΔM^*mutant and contribute to its ovule integument development defects RNA-seq analyses performed on ovules or developing seeds from the mutant would provide valuable insights into the signaling pathways affected in the mutant and provide further insights into the downstream targets and regulatory functions of SEP3 in these tissues.

Finally, in addition to phenotypes in the outer most cell layers of ovules and seeds, our analysis also revealed a significant reduced fitness in the pollen of *sep3^ΔM^* plants (Figure S5). Detailed examination of the RNA-seq data revealed a widespread deregulation of genes involved in pollen fitness and pollen coat formation such as the downregulation of spermidine hydroxycinnamate conjugate biosynthesis genes *SHT, CYP98A8* and *TSM1* (Grienenberger et al. 2009; Elejalde-Palmett et al. 2015; Vogt 2018), and several additional genes required for pollen development and coat formation, including *KCS15, KCS21, KCS7* (β-ketoacyl-CoA synthases), *LTP12* (lipid transfer protein), *CHS* (chalcone synthase), *EXL4–6* (extracellular lipases) *CYP98A9*, *CYP98A8*, *CYP86C3* (cytochrome P450s), *GRP18*, *GRP19* (glycine-rich proteins) and *ABCG9* (ABC transporter) (Mayfield et al. 2001; Choi et al. 2014; Xu et al. 2014; Lu et al. 2020) (Table SI). In addition, *MYB26*, a key regulator of endothecium cell wall thickening required for anther dehiscence and fertility (Yang et al. 2017) was downregulated (log FC −0.9, FDR < 0.05) in *sep3^ΔM^* plants as compared to WT. These observations point to a possibly broader function for SEP proteins in specifying and maintaining epidermal cell identity across reproductive tissues. Indeed, recent work by Chen et al., 2025 proposes that distinct combinations of floral homeotic transcription factors, including SEP3, directly regulate functionally specialized epidermal “hub genes” in an organ-specific manner. Among these target genes are *AtMYB26* and *INO*, both of which are downregulated in the *sep3^ΔM^* mutant. Together, these results strongly support the role of SEP proteins as central regulators of epidermal cell fate within reproductive structures.

By selectively excising the DNA-binding domain of SEP3, additional roles of SEP-containing MADS complexes in reproductive developmental programs were demonstrated. Our phenotypic and gene expression analysis point to SEP3 as a hub connecting MADS-complex mediated regulation of ovule, pollen, and fruit development, ensuring the coordination of male and female reproductive structures required for successful fertilization and seed production. Furthermore, *SEP^ΔM^* may be a valuable tool to further dissect MADS tetramer function if expressed under cell-specific promoters, allowing us to uncouple the various phenotypes observed for *sep3^ΔM^* plants and thereby providing deeper insight into the distinct biological activities of different SEP-containing MTF complexes.

## Materials and Methods

### Plant material and growth conditions

All experiments were performed using *Arabidopsis thaliana* WT and MADS mutants in the Col-0 background. The *sep3* and *sep3^ΔM^*mutants were generated by CRISPR-Cas9 and corresponded to the *sep3-3* and *sep3-4* mutants respectively (Hugouvieux et al. 2024). The *sep3* mutant carries a 1,081-bp deletion removing the last 38 bp of intron 1 and up to the 83^rd^ bp of exon 8. The *sep3^ΔM^* mutant carries a deletion of 963 bp starting from the 25^th^ bp of exon 1. The *stk* mutant carries a 74 nucleotides insertion near the splice site of the 3^rd^ intron, and corresponds to the *stk-*2 allele (Pinyopich et al. 2003). Both, the *sep3* and the *stk* mutants were used as controls and compared to *sep3^ΔM^* for ovule and seed phenotype analysis. Seedlings were grown in parallel in controlled growth chambers in long day conditions (16h light**/**8h dark) at 22^◦^C for plant transformation, and flower and seed phenotype analysis.

### Generation and genotyping of the *sep3^ΔM^* mutant

The *sep3^ΔM^* mutation was previously generated using CRISPR-Cas9 genome editing (Hugouvieux et al. 2024). On 23 primary transformants carrying the CRISPR-Cas9 construct, PCR was performed on cauline leaves with the two following primers outside the expected deletion (Fwd 5’ gcttgcaaggaaagagagagag3’; Rev 5’ caagagaggccttagcagttgt 3’) to identify potential mutants deleted in the SEP3 MADS DNA-binding domain. Seven plants were identified that carried the deletion. T2 seeds from four positive T1 plants were sown on soil to further identify plants that had retained the deletion but lost the plasmid, to prevent potential additional unwanted Cas9 activity. To do so, a PCR was performed on plant genomic DNA (Hugouvieux et al. 2024). Stable *sep3^ΔM^*lacking the CRISPR-Cas9 plasmid were identified from two independent lines. The *SEP3^ΔM^* locus was sequenced in these two lines to ensure that deletion did not trigger frame-shift (Hugouvieux et al. 2024) and one of them was selected for further characterization.

### Vector construction for complementation analyses and expression of *SEP3^ΔM^*

For complementation analysis of the *sep3^ΔM^* mutant phenotypes, *pSEP3:SEP3* (ABRC stock number CD3-2708) (Hugouvieux et al. 2018) was used. This plasmid allows the expression of *SEP3* under the control of the *SEP3* promoter as described previously (Hugouvieux et al. 2018). The vector backbone, *pFP100*, allows GFP expression in seeds for selection of transformants (Bensmihen et al. 2004). The vector *pSEP3:SEP3^ΔM^* was constructed as described for the above plasmid using the PCR amplified *SEP3* promoter, the *SEP3^ΔM^* coding sequence, and the SEP3 terminator using the three pairs of primers (Fwd 5’ cctgcaggtcgactctagaggtaattcaacatgtagcatag 3’ and Rev 5’ctcttcccattttctttttctttctcctctc 3’; Fwd 5’ gaaaaagaaaatgggaagagggagagtag 3’ and Rev 5’ tgagaaagattcaaatagagttggtgtcataag 3’; Fwd ctctatttgaatctttctcacttaatcaatcc and Rev aaaatccagtggtacccgggccattactatacatcaagagg) respectively. The three rsulting PCR fragments were cloned by Gibson assembly into the BamHI linearized *pFP100* vector. Transgenes were inserted into Arabidopsis plants using the floral dip method (Clough and Bent 1998). Transformants were selected based on the fluorescence of GFP-positive seeds.

### Morphological analysis of flowers, silique, stamen, carpel, seeds, and fertilization efficiency measurements

For flower phenotype analysis at different stage of development, a minimum of ten flowers per genotype were analyzed and photographed using a digital microscope (KEYENCE S5550E). For silique, pistil, stamen and seeds measurements, samples were stick onto a glass slide covered with double-sided tape. For stamen and pistil measurements, flowers at anthesis were stick as described above, and sepals and petals were gently removed from each flower with a syringe needle. Pistil and stamen were then detached for each flower, and stick onto the glass slide. Length of about 15 to 20 representative siliques was measured under a binocular microscope (Nikon SMZ800N) with a graduated ruler. Length of pistil and stamen of about 30 flowers, and length and width of about 180 seeds were carried out using the “measurement between two points” option of the KEYENCE VHX5000 digital microscope. For fertilization efficiency analysis, the valves of 10 representative silique within a batch of 3 plants were opened using syringe needles. Structures emerging from the septum were counted for each silique and categorized as non-fertilized ovule, aborted seeds or viable seeds.

### Environmental scanning electron microscopy of seeds

Scanning electron microscopy (SEM) experiments were performed at the Electron Microscopy Facility of the Institut de Chimie Moleculaire of Grenoble Nanobio-Chemistry Platform, as previously described (Hugouvieux et al. 2024). Dried and 16h water imbibed seed were directly observed. Care was taken to maintain humidity during the pressure decrease in the chamber in order to prevent tissue drying. Secondary electron images were recorded with a Quanta FEG 250 (FEI) microscope while maintaining the tissue at 2 °C, under a pressure of 500 Pa and 70% relative humidity. The accelerating voltage was 14 kV and the image magnification ranged from 100 to 800Å. At least three seeds from each genotype were observed.

### Electrophoretic mobility shift assays (EMSA)

Vectors containing *AG* (At4g18960.1), *SEP3* (At1g24260.2) and *STK* (AT4G09960.1) cDNAs were previously described (Kaufmann et al. 2005a; Hugouvieux et al. 2018). *SEP2* (At3g02310) was cloned with a C terminus FLAG tag as described for SEP3 (Lai et al. 2020). The vector containing *SEP3^ΔM^* was PCR amplified using the *SEP3* containing vector as template, with the two specific primers (Fwd 5’ <u>gaattggagaggtaccaaaa</u>gtgtaac 3’; Rev 5’ gtacctctccaattctactctccctcttccc 3’) allowing the removal of the DNA binding M-domain, after ligation by Gibson Assembly^®^. These vectors were used for *in vitro* protein production using SP6 High-Yield Wheat Germ Protein Expression System (Promega L3260) according to the manufacturer’s instructions. EMSAs were performed as described previously (Hugouvieux et al. 2018). The *SEP3* promoter probe contains two CArG-box-binding sites of predicted high affinity. Probes were labeled with Cy5 (Eurofins) for visualization. For each EMSA, a negative control was run corresponding to the labeled DNA incubated with *in vitro* transcription and translation mix and empty *pSP64* vector.

### RNA-seq experiments and data analysis

WT and *sep3^ΔM^* plants were grown in parallel and total RNA were extracted in duplicate, from both genotypes, from inflorescence meristems with small closed buds up to stage 10– 11(Smyth et al. 1990) as described by (Lai et al. 2020). Quality of the total RNA was validated by their 260*/*280 absorbance ratio and the integrity of the ribosomal RNA by gel electrophoresis. RNA library construction and sequencing was performed by GENEWIZ (USA) using Illumina HiSeq and 2x 150bp configuration as described (Lai et al. 2020). Between 25 and 35 million reads were obtained for each library. Mapping onto the *Arabidopsis* genome (TAIR10), read count per gene and statistical analysis were done using STAR (no multimapping, mismatch number < 10), FeatureCount (default parameters) and EdgeR (default parameters), respectively, available in the Galaxy platform (Afgan et al. 2018). Unless stated, genes were considered differentially expressed (DE) between WT and *sep3^ΔM^* plants when the log FC was > 1 or <**−** 1 and the FDR < 0,05.

### Gene expression quantification by real time qRT-PCR

Complementary DNA (cDNA) was primed with random hexamers on 1µg DNase-treated RNA using Superscript III Reverse Transcriptase (Thermo Fisher Scientific). Real time qRT-PCR were performed and analysed as described (Hugouvieux et al. 2018) from the two biological replicates used for RNA seq analysis. For *SEP3^ΔM^*quantification, quantitative PCR was performed using one set of primers specific to *SEP3^ΔM^* encoding cDNA (Fwd P2 5’ gagagtagaattggagaggtac 3’; Rev P3 5’cagtcagcatgcgttcctta 3’) and the other set amplifying both mutant and WT cDNA (Fwd P1 5’gcagttgaacttagtagccag 3’; and Rev P3 5’ cagtcagcatgcgttcctta 3’), to evaluate the level of expression of *SEP3^ΔM^* (Figure SI). For *INO*, *NARS1*, *NARS2* and *GA2ox3* quantification, the following primers were used; INO Fwd 5’aatggtggtgactgtgagatg 3’; *INO* Rv 5’gcaaggagatggagaggaatg 3’; *NASR1* Fwd 5’ agtccacgagatcggaagtatc 3’; *NARS1* Rv: 5’cggaagcaagtaccggtttatc 3’; *NARS2* Fwd 5’tcgccgacgttgatctttac 3’; *NARS2* Rv 5’ccgttgggatatttccgatctc3’; *Ga2OX3* Fwd 5’ctcagatcaaacgacacagagg3’; *GA2Ox3* Rv 5’ ctccgacaagaacgaagaaaga3’. *ACTIN2* (At3g18780) and/or *EF-1α* (At1g09740) were used to normalize the data (Hugouvieux et al. 2018).

### Confocal laser scanning microscopy of ovules

A protocol adapted from (Vijayan et al. 2021) was used to perform confocal analysis on ovules from flowers at stage 13 (Smyth et al. 1990). Flowers were dissected on double-sided tape under a stereomicroscope to isolate the carpel from the other floral pieces and were opened longitudinally to expose the ovules. Dissected carpels were transferred to 1X Dulbecco’s Phosphate Buffered Saline (D-PBS, Gibco 14200067) and fixed in 4% PFA (EMS EM-15710) at 4°C for at least one night and up to one month. Samples were then rinsed twice with 1X D-PBS, cleared in a ClearSee solution (10% xylitol [w/v], 15% sodium deoxycholate [w/v], 25% urea [w/v], H_2_O to final volume) at room temperature overnight with gentle agitation. The cleared tissues were stained at room temperature for 20 min in a 1X D-PBS solution containing 0.1% SR2200 stain (Renaissance Chemicals, Selby, UK) to visualize cell walls and, finally, transferred back into ClearSee. Carpels were open on a microscopy slide in a drop of ClearSee to expose ovules and covered with a coverslip. Confocal imaging was performed on a commercial upright Leica TCS SP8 (DM6000 CS) confocal laser scanning microscope equipped with a 40x water immersion objective (Leica HCX APO L U-V-I 40x/0.8) and a Leica HyD hybrid detector. SR2200 was excited with a 405 nm laser diode (power set to 0.1-1%), and detected at 420-500 nm. 16-bit unidirectional scans of a 1024 × 1024 pixels region of interest were acquired at 8000 Hz (resonant mode: on) with a pixel size of 227.27 nm, and a z-step size of 240 nm. The pinhole was set either at 1 or 0.6, and line averaging was set at least to 8.

### Differential Interference Contrast (DIC) analysis of seeds

Flowers were harvested three days after pollination (DAP) for WT, *sep3*, *stk* and *sep3^ΔM^* lines complemented with SEP3. For *sep3^ΔM^* manual fertilization was performed to synchronize and enhance seed production. Siliques were excised from the flowers with tweezers, attached to a slide covered with double-sided tape, and open longitudinally with a needle. Open siliques were then placed in fixative (EtOH:acetic acid 9:1) and incubated overnight at 4°C. Samples were washed in 90% EtOH and transferred to 70% EtOH at 4°C overnight (or up to several months). Seed clearing was performed in a chloral hydrate (VWR 87804.180) solution (chloral hydrate:glycerol 7:1) at 4°C overnight. Seeds were removed from the siliques on a slide in a droplet of clearing solution, covered with a coverslip, and imaged with an Axio Imager. M2 microscope (Zeiss) in DIC mode, equipped with an Axio Cam 705 Color camera (Zeiss), and EC Plan-NEOFLUAR 10x/0.3, 20x/0.5 and 40x/0.75 objectives.

### Image analysis

Confocal and DIC microscopy images were analyzed with ImageJ version 1.54g and Fiji (Schindelin et al. 2012).

### Mucilage release experiment

Experiments were performed as described by (Ezquer et al. 2016). Briefly, whole seeds were incubated in a 0.01% [w/v] ruthenium red (Sigma-Aldrich) solution for 90min. Seeds treated with EDTA (0.5 M) were imbibed for 2 h before staining with ruthenium red. Samples were rinsed in deionized water prior to visualization. Experiments were performed on at least 50 seeds from at least 2 different batches of seeds.

### Proanthocyanidin accumulation (PA)

PA accumulation was analyzed as reported (Di Marzo et al. 2022b). Fresh seeds were removed from the siliques three DAP with a needle and placed on a microscopy slide. A drop of solution containing 1% (w/v) vanillin (Sigma-Aldrich) and 5 M HCl was placed on the sample which was then incubated at room temperature for 5 minutes. Samples were observed by using a Zeiss Axiophot D1 microscope and pictures acquired with an Axiocam MRc5 camera (Zeiss) using the Axiovision program (version 4.1).

### Statistical analysis

The robustness of sample comparisons two by two was tested using a Wilcoxon & Mann-Witney non-parametric test (Kruskal and Wallis, 1952) with Excel software (version 2607). The p-value significance threshold was set at 0,05.

## Acknowledgments

The authors would like to thank K. Kaufmann and C. Smaczniak for *STK* plasmids, C. Mizzotti for the *stk* mutant and G. Tichtinsky for helpful discussions. The authors thank the NanoBio-ICMG Platform (UAR 2607, Grenoble) for granting access to the Electron Microscopy facility and Christine Lancelon-Pin for assistance.

## Data availability

RNA- seq data sets have been deposited in the GEO database under accession number GSE337717.

## Author contributions

V.H. and C.Z. conceived the study and designed the experimental strategy. A.J., L.R-L., N.B., V.H., M.P., A.M. and L.M. generated the data. All authors contributed to data interpretation. V.H. and C.Z. wrote the manuscript with input from all authors.

## Funding

This project received support from the Agence Nationale de la Recherche (ANR-23-CE20-0019 and ANR-21-CE11-0037) and GRAL, a program from the Chemistry and Biology Health Graduate School of the Université Grenoble Alpes (ANR-17-EURE-0003), with a thesis fellowship to A.J. .

## Conflict of interest statement

None declared.

## Notes

### Competing Interest Statement

The authors have declared no competing interest.

