## supplemental Table 1 for "SEPALLATA MADS transcription factors act as key regulators in fertilization efficiency, ovule outer integument growth and mucilage secretory cell differentiation in Arabidopsis"

Deregulated genes in sep3delM mutant

| GeneID | Symbol | GeneName | FDR | logFC |
|---|---|---|---|---|
| AT1G51210 | NA | UDP-Glycosylt | 0,00036179 | -8,32437684 |
| AT3G44300 | AtNIT2 | nitrilase 2 | 2,07E-05 | -5,76960006 |
| AT1G18520 | TET11 | tetraspanin11 | 8,09E-05 | -5,325412 |
| AT1G43160 | RAP2.6 | related to AP2 | 0,0024853 | -5,32348765 |
| AT3G22740 | HMT3 | homocysteine | 0,02191701 | -4,09326664 |
| AT1G25400 | NA | NA | 0,00091554 | -3,57383194 |
| AT3G49300 | NA | proline-rich fa | 0,00189543 | -3,47967798 |
| AT2G23800 | GGPS2 | geranylgerany | 0,00017019 | -3,2137888 |
| AT3G48520 | CYP94B3 | cytochrome P | 0,00815194 | -3,16891551 |
| AT2G43820 | ATSAGT1 | UDP-glucosylt | 0,00021455 | -3,15632843 |
| AT3G11340 | NA | UDP-Glycosylt | 0,00180201 | -3,087659 |
| AT3G15500 | ANAC055 | NAC domain c | 0,00062473 | -3,0116895 |
| AT2G17940 | NA | Plant protein ( | 0,00107086 | -2,98930753 |
| AT1G06990 | NA | GDSL-like Lipa | 1,50E-05 | -2,97982196 |
| AT1G33700 | NA | Beta-glucosida | 3,00E-05 | -2,94601323 |
| AT1G26710 | NA | NA | 1,09E-05 | -2,83927289 |
| AT2G29880 | NA | NA | 0,00152978 | -2,8124662 |
| AT1G74550 | CYP98A9 | cytochrome P | 0,02793212 | -2,79057346 |
| AT4G37990 | ATCAD8 | elicitor-activat | 0,00307797 | -2,78691529 |
| AT4G19690 | ATIRT1 | iron-regulatec | 1,26E-05 | -2,6980481 |
| AT3G26125 | CYP86C2 | cytochrome P | 1,52E-05 | -2,69533954 |
| AT3G57157 | NA | NA | 0,00399675 | -2,67316601 |
| AT2G19070 | SHT | spermidine hy | 9,46E-09 | -2,65490754 |
| AT3G22235 | NA | NA | 0,00680227 | -2,63183267 |
| AT1G75920 | NA | GDSL-like Lipa | 1,50E-05 | -2,61532278 |
| AT1G75940 | ATA27 | Glycosyl hydro | 0,00019535 | -2,59897782 |
| AT2G28990 | NA | Leucine-rich r | 0,00742554 | -2,52454017 |
| AT3G21720 | ICL | isocitrate lyas | 4,59E-05 | -2,51333054 |
| AT1G23580 | NA | Domain of unl | 0,00356171 | -2,51199579 |
| AT5G41750 | NA | Disease resista | 0,00024021 | -2,50607553 |
| AT2G18420 | NA | Gibberellin-re | 0,00303265 | -2,49327176 |

| | | | | |
|---|---|---|---|---|
| AT2G27080 | NA | Late embryog | 0,00957126 | -2,4686213 |
| AT1G20120 | NA | GDSL-like Lipa | 1,74E-06 | -2,42813385 |
| AT1G44224 | NA | ECA1 gametog | 0,00011938 | -2,38866317 |
| AT1G28480 | GRX480 | Thioredoxin su | 0,00773055 | -2,37499287 |
| AT5G07600 | NA | Oleosin family | 1,77E-05 | -2,35525431 |
| AT1G48940 | AtENODL6 | early nodulin- | 0,00025449 | -2,35476431 |
| AT3G48360 | ATBT2 | BTB and TAZ d | 0,00533747 | -2,34357316 |
| AT2G26530 | AR781 | Protein of unk | 0,00089452 | -2,32510248 |
| AT1G30020 | NA | Protein of unk | 0,00052665 | -2,32435651 |
| AT1G80840 | ATWRKY40 | WRKY DNA-bi | 0,00071051 | -2,32220806 |
| AT5G07520 | ATGRP-8 | glycine-rich pr | 1,09E-05 | -2,31336389 |
| AT1G52690 | NA | Late embryog | 0,00019535 | -2,30364537 |
| AT3G44860 | FAMT | farnesoic acid | 0,02892268 | -2,30267774 |
| AT3G15510 | ANAC056 | NAC domain c | 0,0303236 | -2,29869943 |
| AT3G28740 | CYP81D1 | Cytochrome P | 0,00106508 | -2,28837302 |
| AT5G28470 | NA | Major facilitat | 7,99E-06 | -2,28726171 |
| AT5G28237 | NA | Pyridoxal-5pri | 0,01483822 | -2,28451214 |
| AT1G78750 | NA | F-box/RNI-like | 0,00535875 | -2,26295744 |
| AT1G67110 | CYP735A2 | cytochrome P | 0,00962307 | -2,25866817 |
| AT3G58390 | NA | Eukaryotic rel | 0,00707989 | -2,24571077 |
| AT5G60140 | NA | AP2/B3-like tr | 3,83E-05 | -2,24522724 |
| AT5G48210 | NA | Protein of unk | 0,00298469 | -2,22767852 |
| AT2G46400 | ATWRKY46 | WRKY DNA-bi | 0,00265058 | -2,21949301 |
| AT5G52760 | NA | Copper transp | 0,00398139 | -2,21748596 |
| AT1G10747 | NA | Maternally ex | 0,00369182 | -2,21516617 |
| AT4G37900 | NA | Protein of unk | 5,33E-09 | -2,2095976 |
| AT2G03740 | NA | late embryoge | 5,10E-08 | -2,20484177 |
| AT1G61110 | anac025 | NAC domain c | 3,36E-05 | -2,18311342 |
| AT4G17250 | NA | NA | 0,03627371 | -2,17112741 |
| AT2G36110 | NA | Polynucleotid | 0,00966915 | -2,15675794 |
| AT4G29250 | NA | HXXXD-type a | 8,63E-08 | -2,14729823 |
| AT5G44400 | NA | FAD-binding B | 0,01240015 | -2,12242722 |
| AT3G45060 | ATNRT2.6 | high affinity n | 1,09E-05 | -2,11751977 |

| | | | | |
|---|---|---|---|---|
| AT2G47180 | AtGolS1 | galactinol synt | 2,72E-08 | -2,10126207 |
| AT2G21890 | ATCAD3 | cinnamyl alco | 0,0042695 | -2,09350185 |
| AT1G49120 | NA | Integrase-type | 0,00351916 | -2,08639138 |
| AT3G47340 | ASN1 | glutamine-dep | 0,00406542 | -2,0823172 |
| AT4G28090 | sks10 | SKU5 similar | 0,00312165 | -2,07573351 |
| AT5G17500 | NA | Glycosyl hydro | 0,02146494 | -2,07018713 |
| AT1G68875 | NA | NA | 1,72E-07 | -2,06468964 |
| AT2G32510 | MAPKKK17 | mitogen-activ | 0,0133226 | -2,06322581 |
| AT3G17320 | NA | F-box and ass | 0,00826225 | -2,03698874 |
| AT4G36740 | ATHB40 | homeobox pro | 0,00107838 | -2,03584995 |
| AT4G01430 | NA | nodulin MtN2 | 6,42E-07 | -2,0342003 |
| AT1G15460 | ATBOR4 | HCO3- transpo | 0,00102756 | -2,02643295 |
| AT1G44191 | NA | ECA1 gametog | 0,00032919 | -2,02441443 |
| AT5G62240 | NA | Cell cycle regu | 0,02161879 | -2,01764281 |
| AT5G44610 | MAP18 | microtubule-a | 0,01974026 | -2,0120982 |
| AT1G74540 | CYP98A8 | cytochrome P | 1,58E-08 | -2,00407148 |
| AT1G26720 | NA | NA | 1,09E-06 | -1,99869755 |
| AT1G67980 | CCOAMT | caffeoyl-CoA 3 | 0,00647304 | -1,99633734 |
| AT5G16960 | NA | Zinc-binding d | 8,23E-07 | -1,99154312 |
| AT3G26940 | CDG1 | Protein kinase | 0,00798739 | -1,98159781 |
| AT4G19460 | NA | UDP-Glycosylt | 0,00028378 | -1,97687571 |
| AT5G60500 | NA | Undecaprenyl | 1,96E-05 | -1,97641264 |
| AT4G01450 | NA | nodulin MtN2 | 0,00077926 | -1,97605923 |
| AT4G24040 | ATTRE1 | trehalase 1 | 0,00011366 | -1,96807543 |
| AT5G59845 | NA | Gibberellin-re | 5,52E-06 | -1,96705045 |
| AT4G26390 | NA | Pyruvate kina | 0,00826948 | -1,96480924 |
| AT3G42850 | NA | Mevalonate/g | 1,17E-05 | -1,9540771 |
| AT4G15530 | PPDK | pyruvate orth | 0,00599578 | -1,95399678 |
| AT1G67990 | ATTSM1 | S-adenosyl-L-r | 2,54E-08 | -1,95306161 |
| AT4G27420 | NA | ABC-2 type tra | 1,09E-05 | -1,95187338 |
| AT1G73220 | 01-oct | organic cation | 0,00481846 | -1,94206844 |
| AT4G37520 | NA | Peroxidase su | 0,00534915 | -1,93904265 |
| AT4G10260 | NA | pfkB-like carb | 0,00756358 | -1,93583629 |

| | | | | |
|---|---|---|---|---|
| AT4G08670 | NA | Bifunctional ir | 0,00318147 | -1,93425944 |
| AT5G19640 | NA | Major facilitat | 0,00410908 | -1,93265521 |
| AT2G35090 | NA | Protein of unk | 0,02831811 | -1,91915262 |
| AT5G37870 | NA | Protein with R | 0,02831811 | -1,91009946 |
| AT3G19030 | NA | NA | 0,02691091 | -1,90896596 |
| AT3G58210 | NA | TRAF-like fam | 0,00782364 | -1,90461208 |
| AT3G60100 | CSY5 | citrate syntha | 0,00844937 | -1,90452274 |
| AT2G31760 | ARI10 | RING/U-box su | 0,01111179 | -1,89390621 |
| AT5G45640 | NA | Subtilisin-like | 0,0308909 | -1,88841946 |
| AT1G20132 | NA | GDSL-like Lipa | 0,00020161 | -1,88275133 |
| AT3G03910 | GDH3 | glutamate deh | 0,02666545 | -1,88113606 |
| AT4G15100 | scpl30 | serine carboxy | 0,00905887 | -1,87991232 |
| AT3G43910 | NA | NA | 0,00925841 | -1,8798577 |
| AT2G24210 | TPS10 | terpene synth | 2,35E-05 | -1,8787865 |
| AT2G18130 | ATPAP11 | purple acid ph | 0,01505899 | -1,87429023 |
| AT5G49070 | KCS21 | 3-ketoacyl-Co | 2,12E-05 | -1,86978761 |
| AT5G59810 | ATSBT5.4 | Subtilase fami | 0,0393588 | -1,86666303 |
| AT2G39510 | NA | nodulin MtN2 | 1,09E-05 | -1,86484547 |
| AT3G46520 | ACT12 | actin-12 | 0,00015525 | -1,85054082 |
| AT2G46880 | ATPAP14 | purple acid ph | 8,50E-05 | -1,84419167 |
| AT1G23060 | NA | NA | 0,003387 | -1,84022581 |
| AT2G28120 | NA | Major facilitat | 0,00240712 | -1,83533018 |
| AT3G58240 | NA | TRAF-like supe | 0,01275069 | -1,83391811 |
| AT1G31740 | BGAL15 | beta-galactosi | 0,01549151 | -1,81546637 |
| AT3G04660 | NA | F-box and ass | 0,03434465 | -1,81462272 |
| AT5G57240 | ORP4C | OSBP(oxyster | 0,0052304 | -1,81437956 |
| AT5G52570 | B2 | beta-carotene | 0,01137379 | -1,81298813 |
| AT5G60510 | NA | Undecaprenyl | 1,54E-06 | -1,81177738 |
| AT5G64120 | NA | Peroxidase su | 0,00632181 | -1,81115657 |
| AT3G15830 | NA | phosphatidic a | 0,00018074 | -1,80667446 |
| AT1G65342 | NA | NA | 0,01355741 | -1,80406744 |
| AT4G18550 | NA | alpha/beta-Hy | 4,78E-06 | -1,80003621 |
| AT2G37290 | NA | Ypt/Rab-GAP | 0,01231717 | -1,79056477 |

| | | | | |
|---|---|---|---|---|
| AT5G35110 | NA | NA | 0,01865457 | -1,78878449 |
| AT4G13700 | ATPAP23 | purple acid ph | 0,018124 | -1,78740647 |
| AT1G49500 | NA | NA | 0,0139001 | -1,7860518 |
| AT3G49020 | NA | FBD F-box anc | 0,00945837 | -1,78592832 |
| AT1G23560 | NA | Domain of unl | 1,03E-05 | -1,76615324 |
| AT3G23430 | ATPHO1 | phosphate 1 | 0,01032082 | -1,7648411 |
| AT5G13600 | NA | Phototropic-re | 0,02387817 | -1,76019458 |
| AT4G14815 | NA | Bifunctional ir | 5,80E-08 | -1,75936617 |
| AT2G36190 | AtcwINV4 | cell wall inver | 0,00431229 | -1,75796006 |
| AT3G03760 | LBD20 | LOB domain-c | 2,09E-06 | -1,7567998 |
| AT5G44050 | NA | MATE efflux fa | 0,03627371 | -1,75633538 |
| AT2G34555 | ATGA2OX3 | gibberellin 2-c | 0,00487738 | -1,75472487 |
| AT1G15010 | NA | NA | 0,03026121 | -1,74222929 |
| AT5G13370 | NA | Auxin-respons | 0,01192026 | -1,74056184 |
| AT1G18120 | NA | NA | 1,56E-05 | -1,7404055 |
| AT1G80120 | NA | Protein of unk | 0,00976658 | -1,73232307 |
| AT5G45116.1 | NA | NA | 0,00524888 | -1,72710531 |
| AT3G51590 | LTP12 | lipid transfer p | 0,00527453 | -1,71432652 |
| AT5G53510 | ATOPT9 | oligopeptide t | 0,04600234 | -1,70807146 |
| AT3G45670 | NA | Protein kinase | 0,04169355 | -1,70560121 |
| AT1G56360 | ATPAP6 | purple acid ph | 0,00531423 | -1,69615117 |
| AT3G59845 | NA | Zinc-binding d | 0,04174356 | -1,68554421 |
| AT3G28315.1 | NA | NA | 0,00795791 | -1,68080597 |
| AT1G10070 | ATBCAT-2 | branched-cha | 0,00042003 | -1,6693924 |
| AT3G18360 | NA | VQ motif-cont | 0,01372626 | -1,6623016 |
| AT1G32960 | ATSBT3.3 | Subtilase fami | 0,03564823 | -1,66203635 |
| AT4G04460 | NA | Saposin-like a: | 1,28E-07 | -1,66014345 |
| AT5G46590 | anac096 | NAC domain c | 0,0003142 | -1,65743793 |
| AT1G27860 | NA | Protein of unk | 0,00826225 | -1,65728434 |
| AT1G18280 | NA | Bifunctional ir | 0,04362713 | -1,65697635 |
| AT4G02380 | AtLEA5 | senescence-as | 0,00153709 | -1,64818306 |
| AT5G59040 | COPT3 | copper transp | 0,03453338 | -1,64362766 |
| AT4G04760 | NA | Major facilitat | 0,00092904 | -1,64230299 |

| | | | | |
|---|---|---|---|---|
| AT5G17340 | NA | Putative mem | 0,03190587 | -1,6367263 |
| AT5G39610 | ANAC092 | NAC domain c | 0,04813156 | -1,63153516 |
| AT2G02020 | NA | Major facilitat | 0,03778658 | -1,63150646 |
| AT5G13380 | NA | Auxin-respons | 1,17E-05 | -1,62083831 |
| AT1G72980 | LBD7 | LOB domain-c | 0,01081414 | -1,61639783 |
| AT3G26820 | NA | Esterase/lipas | 0,00034435 | -1,61457047 |
| AT4G39760 | NA | Galactose oxi | 0,03101266 | -1,59814559 |
| AT1G24400 | AATL2 | lysine histidin | 4,59E-05 | -1,59394849 |
| AT5G39670 | NA | Calcium-bindi | 0,0454883 | -1,59027236 |
| AT4G35733 | NA | F-box family p | 0,03318374 | -1,58994186 |
| AT4G14365 | XBAT34 | XB3 ortholog | 0,00011771 | -1,58885531 |
| AT3G52160 | KCS15 | 3-ketoacyl-Co | 1,31E-07 | -1,58872863 |
| AT1G02520 | PGP11 | P-glycoprotei | 2,42E-05 | -1,58587083 |
| AT1G56540 | NA | Disease resist | 0,01951124 | -1,57228579 |
| AT2G32460 | ATM1 | myb domain p | 0,00033718 | -1,57027552 |
| AT4G28397 | NA | NA | 9,34E-06 | -1,56565376 |
| AT1G28700 | NA | Nucleotide-di | 1,56E-05 | -1,56465495 |
| AT5G54330 | NA | Protein of unk | 0,01599402 | -1,56183929 |
| AT3G48650 | NA | NA | 0,0391161 | -1,56145161 |
| AT2G23970 | NA | Class I glutam | 0,03394031 | -1,56014719 |
| AT1G73805 | NA | Calmodulin bi | 0,0013834 | -1,55564138 |
| AT1G72280 | AERO1 | endoplasmic r | 1,57E-05 | -1,55440438 |
| AT2G41040 | NA | S-adenosyl-L-r | 1,28E-07 | -1,55383429 |
| AT3G45480 | NA | RING/U-box p | 0,0044048 | -1,54403111 |
| AT2G41290 | SSL2 | strictosidine s | 5,20E-06 | -1,54028904 |
| AT1G78160 | APUM7 | pumilio 7 | 0,001327 | -1,54027572 |
| AT5G17590 | NA | Putative mem | 0,02046492 | -1,53978441 |
| AT3G12143 | NA | NA | 0,00477989 | -1,53090477 |
| AT3G07070 | NA | Protein kinase | 0,025435 | -1,5268683 |
| AT2G20970 | NA | NA | 0,00268593 | -1,5257274 |
| AT2G21237 | NA | NA | 0,02954402 | -1,5256179 |
| AT4G01533 | NA | NA | 0,02739605 | -1,52227259 |
| AT3G28540 | NA | P-loop contair | 0,00535875 | -1,51307567 |

| | | | | |
|---|---|---|---|---|
| AT5G08030 | NA | PLC-like phosp | 0,00079608 | -1,51268442 |
| AT1G51820 | NA | Leucine-rich r | 0,03564327 | -1,51236081 |
| AT1G32780 | NA | GroES-like zin | 5,33E-06 | -1,51145339 |
| AT1G75490 | NA | Integrase-type | 0,03048836 | -1,5072831 |
| AT3G11680 | NA | Aluminium ac | 0,02601519 | -1,50662235 |
| AT2G21730 | ATCAD2 | cinnamyl alcol | 0,00423525 | -1,50486572 |
| AT4G00160 | NA | F-box/RNI-like | 0,02007448 | -1,50439941 |
| AT5G66110 | NA | Heavy metal t | 0,02865194 | -1,50358162 |
| AT4G36350 | ATPAP25 | purple acid ph | 0,00658482 | -1,50240029 |
| AT2G39590 | NA | Ribosomal pro | 0,02159733 | -1,49988447 |
| AT1G13140 | CYP86C3 | cytochrome P | 2,93E-07 | -1,49863671 |
| AT1G07180 | ATNDI1 | alternative N/ | 0,00273959 | -1,494468 |
| AT3G52400 | ATSYP122 | syntaxin of pla | 0,02228886 | -1,48765412 |
| AT5G43340 | PHT1;6 | phosphate tra | 0,00021455 | -1,46791526 |
| AT2G28140 | NA | Protein of unk | 0,00022001 | -1,462322 |
| AT5G09500 | NA | Ribosomal pro | 0,0005417 | -1,45932998 |
| AT5G04370 | NAMT1 | S-adenosyl-L-r | 0,04236663 | -1,45752845 |
| AT5G14180 | MPL1 | Myzus persica | 0,00487738 | -1,43696504 |
| AT4G18900 | NA | Transducin/W | 0,01966359 | -1,43216775 |
| AT4G28395 | A7 | Bifunctional ir | 9,60E-07 | -1,43082475 |
| AT5G40040 | NA | 60S acidic ribo | 0,00791398 | -1,42200425 |
| AT3G25180 | CYP82G1 | cytochrome P | 0,00111242 | -1,41740354 |
| AT3G47870 | LBD27 | LOB domain-c | 0,01800001 | -1,41728209 |
| AT3G23840 | NA | HXXXD-type a | 1,30E-07 | -1,41359206 |
| AT2G32470 | NA | F-box associat | 0,02243229 | -1,406264 |
| AT5G54010 | NA | UDP-Glycosylt | 1,18E-05 | -1,40485814 |
| AT5G66690 | UGT72E2 | UDP-Glycosylt | 0,00010233 | -1,39736855 |
| AT5G50950 | FUM2 | FUMARASE 2 | 1,09E-05 | -1,38019301 |
| AT5G38130 | NA | HXXXD-type a | 0,01340746 | -1,37188374 |
| AT1G22760 | PAB3 | poly(A) bindin | 0,00789896 | -1,37002472 |
| AT2G05380 | GRP3S | glycine-rich pr | 2,93E-07 | -1,36346429 |
| AT1G59740 | NA | Major facilitat | 0,01414067 | -1,3544885 |
| AT3G21670 | NA | Major facilitat | 1,63E-06 | -1,35162358 |

| | | | | |
|---|---|---|---|---|
| AT1G11580 | ATPMEPCRA | methylesteras | 0,00149569 | -1,35102322 |
| AT4G24890 | ATPAP24 | purple acid ph | 0,00095627 | -1,34928678 |
| AT4G32650 | ATKC1 | potassium cha | 0,04311411 | -1,34704468 |
| AT3G58340 | NA | TRAF-like fam | 0,00351916 | -1,34602555 |
| AT5G46240 | KAT1 | potassium cha | 0,00180201 | -1,34535181 |
| AT3G59010 | PME61 | pectin methyl | 0,01022664 | -1,33706834 |
| AT5G38120 | 4CL8 | AMP-depende | 2,39E-05 | -1,33566967 |
| AT2G27120 | POL2B | DNA polymera | 0,00373805 | -1,33267222 |
| AT1G05550 | NA | Protein of unk | 0,00318147 | -1,33097072 |
| AT3G15440 | NA | NA | 0,03033954 | -1,32095435 |
| AT4G19645 | NA | TRAM LAG1 ar | 1,96E-05 | -1,31916768 |
| AT5G59820 | RHL41 | C2H2-type zin | 0,00813363 | -1,3183574 |
| AT3G10116 | NA | COBRA-like ex | 0,00826948 | -1,31630603 |
| AT1G45145 | ATH5 | thioredoxin H- | 3,41E-05 | -1,31534362 |
| AT2G03200 | NA | Eukaryotic asp | 5,46E-06 | -1,315012 |
| AT2G34010 | NA | NA | 0,01757992 | -1,30651732 |
| AT3G52820 | ATPAP22 | purple acid ph | 0,01518374 | -1,30149456 |
| AT5G56970 | ATCKX3 | cytokinin oxid | 0,00090793 | -1,29017158 |
| AT1G70700 | JAZ9 | TIFY domain/I | 0,01333528 | -1,28541834 |
| AT3G26840 | NA | Esterase/lipas | 0,00351916 | -1,28503224 |
| AT2G26940 | NA | C2H2-type zin | 0,01505899 | -1,28371875 |
| AT2G16260 | NA | NA | 0,03778658 | -1,27777209 |
| AT2G20470 | NA | AGC (cAMP-de | 9,27E-05 | -1,26921672 |
| AT2G37900 | NA | Major facilitat | 0,00826225 | -1,26727815 |
| AT5G60330 | NA | NA | 0,0288758 | -1,26282922 |
| AT1G52890 | ANAC019 | NAC domain c | 0,00051176 | -1,26208218 |
| AT5G18090 | NA | AP2/B3-like tr | 0,02407326 | -1,26195213 |
| AT3G04210 | NA | Disease resist | 0,00741416 | -1,26071963 |
| AT5G58390 | NA | Peroxidase su | 0,00398139 | -1,25839532 |
| AT2G12480 | SCPL43 | serine carboxy | 0,00550944 | -1,2580726 |
| AT2G31770 | ARI9 | RING/U-box su | 0,00826225 | -1,25775939 |
| AT5G65300 | NA | NA | 0,04622243 | -1,25470594 |
| AT1G78230 | NA | Outer arm dyr | 0,00051676 | -1,24760983 |

| | | | | |
|---|---|---|---|---|
| AT2G42480 | NA | TRAF-like fam | 0,00546855 | -1,24755058 |
| AT5G38396 | NA | F-box/RNI-like | 0,03758836 | -1,24674195 |
| AT5G15960 | KIN1 | stress-respons | 0,00011366 | -1,2460604 |
| AT1G77210 | AtSTP14 | sugar transpo | 4,59E-05 | -1,23996833 |
| AT5G13330 | Rap2.6L | related to AP2 | 0,01757247 | -1,23922202 |
| AT3G15740 | NA | RING/U-box su | 0,04993678 | -1,2391799 |
| AT2G14710 | NA | F-box family p | 0,02975248 | -1,23482593 |
| AT1G20150 | NA | Subtilisin-like | 0,00491742 | -1,23109374 |
| AT2G16750 | NA | Protein kinase | 0,00015171 | -1,23038129 |
| AT5G24790 | NA | Protein of unk | 0,00254217 | -1,22991427 |
| AT1G34680 | NA | NA | 0,00179912 | -1,22441071 |
| AT2G22030 | NA | Galactose oxid | 0,00887561 | -1,2003448 |
| AT2G19110 | ATHMA4 | heavy metal a | 1,56E-05 | -1,19899694 |
| AT5G65205 | NA | NAD(P)-bindir | 1,18E-05 | -1,19094558 |
| AT4G33050 | EDA39 | calmodulin-bi | 0,00071986 | -1,19005206 |
| AT5G10250 | DOT3 | Phototropic-re | 0,04017832 | -1,18939843 |
| AT1G61630 | ATENT7 | equilibrative r | 0,00205952 | -1,18455732 |
| AT1G56030 | NA | RING/U-box su | 0,01620508 | -1,18414974 |
| AT3G45010 | scpl48 | serine carboxy | 2,02E-06 | -1,1825794 |
| AT3G58260 | NA | TRAF-like fam | 0,02061347 | -1,17744075 |
| AT4G28530 | anac074 | NAC domain c | 0,03846371 | -1,17703627 |
| AT5G19240 | NA | Glycoprotein r | 0,02929797 | -1,17652679 |
| AT1G24470 | ATKCR2 | beta-ketoacyl | 0,00115884 | -1,17565597 |
| AT1G19640 | JMT | jasmonic acid | 0,00198877 | -1,17483015 |
| AT2G05160 | NA | CCCH-type zin | 0,01118712 | -1,1689636 |
| AT5G06520 | NA | SWAP (Suppre | 0,02979525 | -1,16253581 |
| AT4G15490 | UGT84A3 | UDP-Glycosylt | 5,42E-05 | -1,16149005 |
| AT2G40670 | ARR16 | response regu | 0,00255484 | -1,16063448 |
| AT5G41130 | NA | Esterase/lipas | 1,48E-06 | -1,15738799 |
| AT2G31980 | AtCYS2 | PHYTOCYSTAT | 2,70E-05 | -1,14587798 |
| AT1G13150 | CYP86C4 | cytochrome P | 0,00162585 | -1,14299327 |
| AT4G17483 | NA | alpha/beta-Hy | 0,0060787 | -1,13742169 |
| AT1G62305 | NA | Core-2/I-branc | 2,88E-05 | -1,13666901 |

| | | | | |
|---|---|---|---|---|
| AT3G20450 | NA | B-cell recepto | 0,00165356 | -1,13300855 |
| AT5G61620 | NA | myb-like trans | 0,03891078 | -1,13098133 |
| AT5G52330 | NA | TRAF-like supe | 0,01750241 | -1,12610389 |
| AT2G37760 | NA | NAD(P)-linked | 0,00013534 | -1,12599424 |
| AT5G23350 | NA | GRAM domair | 0,00152175 | -1,12573471 |
| AT5G06980 | NA | NA | 0,00312165 | -1,12562282 |
| AT1G17710 | NA | Pyridoxal pho: | 0,03368606 | -1,11401255 |
| AT4G24480 | NA | Protein kinase | 0,0052304 | -1,11272535 |
| AT5G44630 | NA | Terpenoid cyc | 0,01052587 | -1,11247943 |
| AT4G39780 | NA | Integrase-type | 0,02345869 | -1,11123207 |
| AT1G27045 | NA | Homeobox-leu | 0,01887613 | -1,11088184 |
| AT5G43110 | APUM14 | pumilio 14 | 0,00333952 | -1,10845739 |
| AT3G29776.1 | NA | NA | 0,00755422 | -1,1029688 |
| AT1G66760 | NA | MATE efflux fa | 2,93E-05 | -1,10212885 |
| AT1G30220 | ATINT2 | inositol transp | 0,02010216 | -1,1011641 |
| AT1G64030 | ATSRP3 | serpin 3 | 0,01966359 | -1,10021647 |
| AT4G34060 | DML3 | demeter-like p | 0,00516674 | -1,09101867 |
| AT4G18190 | ATPUP6 | purine permea | 0,03490416 | -1,08558921 |
| AT1G14870 | PCR2 | PLANT CADMI | 0,00637981 | -1,08179427 |
| AT4G31800 | ATWRKY18 | WRKY DNA-bi | 0,04796555 | -1,0791162 |
| AT5G47635 | NA | Pollen Ole e 1 | 0,00329603 | -1,06958448 |
| AT3G12320 | NA | NA | 0,03279706 | -1,06943889 |
| AT4G26830 | NA | O-Glycosyl hy | 0,02954402 | -1,06875317 |
| AT3G15400 | ATA20 | anther 20 | 3,23E-06 | -1,06584795 |
| AT2G41415 | NA | Maternally ex | 0,00137786 | -1,06153328 |
| AT3G62860 | NA | alpha/beta-Hy | 0,00826225 | -1,05799246 |
| AT1G71160 | KCS7 | 3-ketoacyl-Co | 0,00013259 | -1,04657395 |
| AT5G04500 | NA | glycosyltransf | 0,0197791 | -1,04505056 |
| AT4G00040 | NA | Chalcone and | 1,59E-05 | -1,03040697 |
| AT1G02813 | NA | Protein of unk | 2,51E-05 | -1,028238 |
| AT3G10340 | PAL4 | phenylalanine | 0,00809482 | -1,02275254 |
| AT5G26340 | ATSTP13 | Major facilitat | 0,00531423 | -1,0211831 |
| AT2G38010 | NA | Neutral/alkali | 1,96E-05 | -1,01810892 |

| | | | | |
|---|---|---|---|---|
| AT4G01883 | NA | Polyketide cyc | 6,02E-05 | -1,01541992 |
| AT2G18193 | NA | P-loop contair | 0,03209303 | -1,00699606 |
| AT1G05680 | UGT74E2 | Uridine diphos | 0,00216133 | -1,00645827 |
| AT2G37180 | PIP2;3 | Aquaporin-like | 0,02415912 | -1,00568332 |
| AT5G14980 | NA | alpha/beta-Hy | 0,04107765 | -1,00229678 |
| AT1G08940 | NA | Phosphoglyce | 0,01317243 | -1,00094643 |
| AT1G06080 | ADS1 | delta 9 desatu | 1,96E-05 | 1,00209171 |
| AT5G36960 | NA | NA | 0,04622243 | 1,00212005 |
| AT2G32530 | ATCSLB03 | cellulose syntl | 0,00673568 | 1,01253599 |
| AT2G21880 | ATRAB7A | RAB GTPase h | 0,04761683 | 1,0129732 |
| AT3G52525 | ATOFP6 | ovate family p | 0,00887225 | 1,04749903 |
| AT5G03790 | ATHB51 | homeobox 51 | 0,00153709 | 1,05905707 |
| AT5G41900 | NA | alpha/beta-Hy | 0,03019353 | 1,0616069 |
| AT2G30550 | NA | alpha/beta-Hy | 0,00052665 | 1,06466764 |
| AT4G09455.1 | NA | NA | 0,01973835 | 1,08469381 |
| AT1G48470 | GLN1;5 | glutamine syn | 0,01121611 | 1,12885852 |
| AT5G42900 | COR27 | cold regulatec | 0,04608454 | 1,16960325 |
| AT3G29410 | NA | Terpenoid cyc | 0,00510624 | 1,20176507 |
| AT4G23240 | CRK16 | cysteine-rich I | 0,02249368 | 1,31600105 |
| AT1G74650 | ATMYB31 | myb domain p | 0,01505899 | 1,33936892 |
| AT3G20810 | NA | 2-oxoglutarate | 0,04040671 | 1,35485618 |
| AT4G29270 | NA | HAD superfam | 2,51E-05 | 1,41585981 |
| AT4G28780 | NA | GDSL-like Lipa | 0,00020161 | 1,47756785 |
| AT4G28160 | NA | hydroxyprolin | 0,00993768 | 1,49154944 |
| AT5G17700 | NA | MATE efflux fa | 0,01982472 | 1,52805032 |
| AT3G21080 | NA | ABC transport | 0,0107602 | 1,57700876 |
| AT4G16560 | NA | HSP20-like cha | 0,01381826 | 1,6460687 |
| AT1G04660 | NA | glycine-rich pr | 0,00844937 | 1,88893924 |
| AT3G58990 | IPMI1 | isopropylmala | 0,01722243 | 2,2854742 |
| AT1G56680 | NA | Chitinase fami | 1,28E-07 | 2,54290298 |
| AT1G62975 | NA | basic helix-loc | 0,00050877 | 2,60270689 |
| AT1G70890 | MLP43 | MLP-like prote | 2,32E-05 | 2,9855585 |

Deregulated genes in both sep3delM and sep1sep2sep3 mutants

| GeneID | Symbol_DEG s | GeneName_D | FDR_DEG sep: | logFC_DEG sep123 | Symbol_DEG s | GeneName_D | FDR_DEG sep: | logFC_DEG sep3 |
|---|---|---|---|---|---|---|---|---|
| AT5G60510 | NA | Undecaprenyl | 0,03823755 | -12,2010826 | NA | Undecaprenyl | 1,54E-06 | -1,81177738 |
| AT5G53510 | ATOPT9 | oligopeptide t | 0,00138765 | -11,4481734 | ATOPT9 | oligopeptide t | 0,04600234 | -1,70807146 |
| AT2G32460 | ATM1 | myb domain p | 1,63E-05 | -10,7362691 | ATM1 | myb domain p | 0,00033718 | -1,57027552 |
| AT5G60500 | NA | Undecaprenyl | 0,03162907 | -10,6325839 | NA | Undecaprenyl | 1,96E-05 | -1,97641264 |
| AT2G19070 | SHT | spermidine hy | 0,02916882 | -10,5856346 | SHT | spermidine hy | 9,46E-09 | -2,65490754 |
| AT3G03760 | LBD20 | LOB domain-c | 0,01697307 | -10,3601205 | LBD20 | LOB domain-c | 2,09E-06 | -1,7567998 |
| AT2G22030 | NA | Galactose oxid | 0,00140151 | -10,3350794 | NA | Galactose oxid | 0,00887561 | -1,2003448 |
| AT2G18130 | ATPAP11 | purple acid ph | 0,03947104 | -10,2460901 | ATPAP11 | purple acid ph | 0,01505899 | -1,87429023 |
| AT3G10116 | NA | COBRA-like ex | 5,11E-05 | -10,2121616 | NA | COBRA-like ex | 0,00826948 | -1,31630603 |
| AT3G51590 | LTP12 | lipid transfer p | 0,03498369 | -10,1607923 | LTP12 | lipid transfer p | 0,00527453 | -1,71432652 |
| AT3G46520 | ACT12 | actin-12 | 0,00684045 | -10,0371259 | ACT12 | actin-12 | 0,00015525 | -1,85054082 |
| AT5G16960 | NA | Zinc-binding d | 0,03453696 | -9,77795749 | NA | Zinc-binding d | 8,23E-07 | -1,99154312 |
| AT4G00160 | NA | F-box/RNI-like | 0,00024729 | -9,25564043 | NA | F-box/RNI-like | 0,02007448 | -1,50439941 |
| AT1G65342 | na | na | 0,00024126 | -9,22772638 | NA | NA | 0,01355741 | -1,80406744 |
| AT1G67990 | ATTSM1 | S-adenosyl-L-r | 0,0319056 | -9,14773944 | ATTSM1 | S-adenosyl-L-r | 2,54E-08 | -1,95306161 |
| AT1G31740 | BGAL15 | beta-galactosi | 0,00312955 | -8,89398822 | BGAL15 | beta-galactosi | 0,01549151 | -1,81546637 |
| AT4G19690 | ATIRT1 | iron-regulatec | 0,01015908 | -8,87169978 | ATIRT1 | iron-regulatec | 1,26E-05 | -2,6980481 |
| AT4G29250 | NA | HXXXD-type a | 0,0162974 | -8,87008509 | NA | HXXXD-type a | 8,63E-08 | -2,14729823 |
| AT2G35090 | NA | Protein of unk | 0,00048644 | -8,86505132 | NA | Protein of unk | 0,02831811 | -1,91915262 |
| AT5G19640 | NA | Major facilitat | 0,00044572 | -8,85853789 | NA | Major facilitat | 0,00410908 | -1,93265521 |
| AT2G36110 | NA | Polynucleotid | 0,00147568 | -8,78616694 | NA | Polynucleotid | 0,00966915 | -2,15675794 |
| AT5G17340 | NA | Putative mem | 0,02142932 | -8,50941236 | NA | Putative mem | 0,03190587 | -1,6367263 |
| AT4G39760 | NA | Galactose oxid | 6,49E-05 | -8,44092577 | NA | Galactose oxid | 0,03101266 | -1,59814559 |
| AT5G37870 | NA | Protein with R | 0,00199599 | -8,43752237 | NA | Protein with R | 0,02831811 | -1,91009946 |
| AT1G18120 | na | na | 0,01760972 | -8,36568121 | NA | NA | 1,56E-05 | -1,7404055 |
| AT1G20150 | NA | Subtilisin-like | 0,00774732 | -8,36365804 | NA | Subtilisin-like | 0,00491742 | -1,23109374 |
| AT4G28397 | na | na | 0,0096737 | -8,28461962 | NA | NA | 9,34E-06 | -1,56565376 |
| AT3G58390 | NA | Eukaryotic rel | 0,00098527 | -8,24845756 | NA | Eukaryotic rel | 0,00707989 | -2,24571077 |
| AT1G18520 | TET11 | tetraspanin11 | 0,02253741 | -8,23532767 | TET11 | tetraspanin11 | 8,09E-05 | -5,325412 |
| AT5G45116.1 | na | na | 0,00462224 | -8,18095055 | NA | NA | 0,00524888 | -1,72710531 |
| AT3G60100 | CSY5 | citrate syntha | 0,00576334 | -8,1701663 | CSY5 | citrate syntha | 0,00844937 | -1,90452274 |

| | | | | | | | | |
|---|---|---|---|---|---|---|---|---|
| AT2G17940 | NA | Plant protein ( | 0,00046357 | -7,99865142 | NA | Plant protein ( | 0,00107086 | -2,98930753 |
| AT3G21720 | ICL | isocitrate lyas | 1,60E-05 | -7,88981077 | ICL | isocitrate lyas | 4,59E-05 | -2,51333054 |
| AT5G13380 | NA | Auxin-respons | 0,02827219 | -7,86637003 | NA | Auxin-respons | 1,17E-05 | -1,62083831 |
| AT4G18190 | ATPUP6 | purine permea | 0,0029566 | -7,46793561 | ATPUP6 | purine permea | 0,03490416 | -1,08558921 |
| AT5G14980 | NA | alpha/beta-Hy | 0,0073971 | -7,42253295 | NA | alpha/beta-Hy | 0,04107765 | -1,00229678 |
| AT4G14815 | NA | Bifunctional ir | 0,00791392 | -7,40441073 | NA | Bifunctional ir | 5,80E-08 | -1,75936617 |
| AT1G23560 | NA | Domain of unl | 0,01131703 | -7,27915416 | NA | Domain of unl | 1,03E-05 | -1,76615324 |
| AT2G46880 | ATPAP14 | purple acid ph | 0,00705779 | -7,25651479 | ATPAP14 | purple acid ph | 8,50E-05 | -1,84419167 |
| AT1G15460 | ATBOR4 | HCO3- transpo | 0,02474239 | -7,18974691 | ATBOR4 | HCO3- transpo | 0,00102756 | -2,02643295 |
| AT4G27420 | NA | ABC-2 type tra | 0,00751788 | -7,04228172 | NA | ABC-2 type tra | 1,09E-05 | -1,95187338 |
| AT5G07600 | NA | Oleosin family | 0,03347974 | -6,94813967 | NA | Oleosin family | 1,77E-05 | -2,35525431 |
| AT1G61110 | anac025 | NAC domain c | 0,02203604 | -6,90387533 | anac025 | NAC domain c | 3,36E-05 | -2,18311342 |
| AT1G28700 | NA | Nucleotide-dip | 0,00796612 | -6,88745435 | NA | Nucleotide-dip | 1,56E-05 | -1,56465495 |
| AT2G03740 | NA | late embryoge | 0,01092132 | -6,84973306 | NA | late embryoge | 5,10E-08 | -2,20484177 |
| AT1G44191 | NA | ECA1 gametog | 0,00272342 | -6,68547391 | NA | ECA1 gametog | 0,00032919 | -2,02441443 |
| AT4G28395 | A7 | Bifunctional ir | 0,01041277 | -6,64335648 | A7 | Bifunctional ir | 9,60E-07 | -1,43082475 |
| AT1G56360 | ATPAP6 | purple acid ph | 0,00225602 | -6,57134315 | ATPAP6 | purple acid ph | 0,00531423 | -1,69615117 |
| AT2G20970 | na | na | 2,46E-05 | -6,55836265 | NA | NA | 0,00268593 | -1,5257274 |
| AT1G02813 | NA | Protein of unk | 0,0086609 | -6,55038273 | NA | Protein of unk | 2,51E-05 | -1,028238 |
| AT5G54010 | NA | UDP-Glycosylt | 0,01211838 | -6,53933038 | NA | UDP-Glycosylt | 1,18E-05 | -1,40485814 |
| AT2G31770 | ARI9 | RING/U-box su | 0,00274855 | -6,47091509 | ARI9 | RING/U-box su | 0,00826225 | -1,25775939 |
| AT3G15830 | NA | phosphatidic a | 0,00414741 | -6,46394048 | NA | phosphatidic a | 0,00018074 | -1,80667446 |
| AT4G36350 | ATPAP25 | purple acid ph | 0,01701183 | -6,40825022 | ATPAP25 | purple acid ph | 0,00658482 | -1,50240029 |
| AT2G12480 | SCPL43 | serine carboxy | 0,00026977 | -6,36875985 | SCPL43 | serine carboxy | 0,00550944 | -1,2580726 |
| AT2G36190 | AtcwINV4 | cell wall inver | 0,00048644 | -6,3456106 | AtcwINV4 | cell wall inver | 0,00431229 | -1,75796006 |
| AT3G52160 | KCS15 | 3-ketoacyl-Co. | 0,01303597 | -6,28946588 | KCS15 | 3-ketoacyl-Co. | 1,31E-07 | -1,58872863 |
| AT3G58260 | NA | TRAF-like fam | 5,90E-05 | -6,26163647 | NA | TRAF-like fam | 0,02061347 | -1,17744075 |
| AT2G32470 | NA | F-box associat | 8,48E-05 | -6,21131687 | NA | F-box associat | 0,02243229 | -1,406264 |
| AT5G59810 | ATSBT5.4 | Subtilase fami | 0,00302317 | -6,12208839 | ATSBT5.4 | Subtilase fami | 0,0393588 | -1,86666303 |
| AT3G26820 | NA | Esterase/lipas | 0,01747504 | -6,06802456 | NA | Esterase/lipas | 0,00034435 | -1,61457047 |
| AT2G41415 | NA | Maternally ex | 0,00407209 | -6,0561655 | NA | Maternally ex | 0,00137786 | -1,06153328 |
| AT5G54330 | NA | Protein of unk | 7,60E-05 | -5,90175003 | NA | Protein of unk | 0,01599402 | -1,56183929 |
| AT1G61630 | ATENT7 | equilibrative r | 3,26E-05 | -5,83884446 | ATENT7 | equilibrative r | 0,00205952 | -1,18455732 |

| | | | | | | | | |
|---|---|---|---|---|---|---|---|---|
| AT1G56030 | NA | RING/U-box su | 1,08E-05 | -5,80518106 | NA | RING/U-box su | 0,01620508 | -1,18414974 |
| AT4G18900 | NA | Transducin/W | 0,00020589 | -5,78786369 | NA | Transducin/W | 0,01966359 | -1,43216775 |
| AT5G38396 | NA | F-box/RNI-like | 0,00082733 | -5,78740518 | NA | F-box/RNI-like | 0,03758836 | -1,24674195 |
| AT5G09500 | NA | Ribosomal pro | 0,00708241 | -5,59835733 | NA | Ribosomal pro | 0,0005417 | -1,45932998 |
| AT1G72980 | LBD7 | LOB domain-c | 0,00020775 | -5,57548345 | LBD7 | LOB domain-c | 0,01081414 | -1,61639783 |
| AT2G23800 | GGPS2 | geranylgerany | 0,03822703 | -5,5720859 | GGPS2 | geranylgerany | 0,00017019 | -3,2137888 |
| AT2G34555 | ATGA2OX3 | gibberellin 2-c | 0,00048167 | -5,53277092 | ATGA2OX3 | gibberellin 2-c | 0,00487738 | -1,75472487 |
| AT3G43910 | na | na | 0,00107931 | -5,47755388 | NA | NA | 0,00925841 | -1,8798577 |
| AT5G43340 | PHT1;6 | phosphate tra | 0,00164682 | -5,44055491 | PHT1;6 | phosphate tra | 0,00021455 | -1,46791526 |
| AT2G42480 | NA | TRAF-like fam | 0,00239899 | -5,42468298 | NA | TRAF-like fam | 0,00546855 | -1,24755058 |
| AT3G04660 | NA | F-box and ass | 0,00015104 | -5,4062682 | NA | F-box and ass | 0,03434465 | -1,81462272 |
| AT5G52330 | NA | TRAF-like supe | 0,00017281 | -5,3712346 | NA | TRAF-like supe | 0,01750241 | -1,12610389 |
| AT2G14710 | NA | F-box family p | 0,00027057 | -5,36618459 | NA | F-box family p | 0,02975248 | -1,23482593 |
| AT3G58340 | NA | TRAF-like fam | 6,34E-06 | -5,34619854 | NA | TRAF-like fam | 0,00351916 | -1,34602555 |
| AT1G20120 | NA | GDSL-like Lipa | 0,03919301 | -5,33533377 | NA | GDSL-like Lipa | 1,74E-06 | -2,42813385 |
| AT5G43110 | APUM14 | pumilio 14 | 2,95E-06 | -5,2637121 | APUM14 | pumilio 14 | 0,00333952 | -1,10845739 |
| AT3G45480 | NA | RING/U-box p | 0,00066489 | -5,25149515 | NA | RING/U-box p | 0,0044048 | -1,54403111 |
| AT4G08670 | NA | Bifunctional ir | 0,01429626 | -5,25013489 | NA | Bifunctional ir | 0,00318147 | -1,93425944 |
| AT5G28470 | NA | Major facilitat | 0,00392743 | -5,24676193 | NA | Major facilitat | 7,99E-06 | -2,28726171 |
| AT5G06520 | NA | SWAP (Suppre | 5,66E-06 | -5,14325546 | NA | SWAP (Suppre | 0,02979525 | -1,16253581 |
| AT3G58210 | NA | TRAF-like fam | 0,00057624 | -5,14104138 | NA | TRAF-like fam | 0,00782364 | -1,90461208 |
| AT3G12143 | na | na | 3,30E-06 | -5,13912168 | NA | NA | 0,00477989 | -1,53090477 |
| AT2G31760 | ARI10 | RING/U-box su | 0,00011438 | -5,03941774 | ARI10 | RING/U-box su | 0,01111179 | -1,89390621 |
| AT4G28090 | sks10 | SKU5 similar 1 | 0,00083037 | -5,03811614 | sks10 | SKU5 similar 1 | 0,00312165 | -2,07573351 |
| AT1G20132 | NA | GDSL-like Lipa | 0,00449238 | -5,03484145 | NA | GDSL-like Lipa | 0,00020161 | -1,88275133 |
| AT4G24890 | ATPAP24 | purple acid ph | 0,01259584 | -4,94479731 | ATPAP24 | purple acid ph | 0,00095627 | -1,34928678 |
| AT4G15100 | scpl30 | serine carboxy | 0,02913456 | -4,92633773 | scpl30 | serine carboxy | 0,00905887 | -1,87991232 |
| AT1G27860 | NA | Protein of unk | 0,00030054 | -4,89881603 | NA | Protein of unk | 0,00826225 | -1,65728434 |
| AT5G57240 | ORP4C | OSBP(oxyster | 0,00721728 | -4,88227727 | ORP4C | OSBP(oxyster | 0,0052304 | -1,81437956 |
| AT1G27045 | NA | Homeobox-leu | 8,01E-06 | -4,86414703 | NA | Homeobox-leu | 0,01887613 | -1,11088184 |
| AT3G26940 | CDG1 | Protein kinase | 0,00103771 | -4,85211251 | CDG1 | Protein kinase | 0,00798739 | -1,98159781 |
| AT5G61620 | NA | myb-like trans | 3,74E-06 | -4,79387909 | NA | myb-like trans | 0,03891078 | -1,13098133 |
| AT4G09455.1 | na | na | 8,01E-06 | -4,78030812 | NA | NA | 0,01973835 | 1,08469381 |

| | | | | | | | | |
|---|---|---|---|---|---|---|---|---|
| AT1G78160 | APUM7 | pumilio 7 | 0,00013114 | -4,76048375 | APUM7 | pumilio 7 | 0,001327 | -1,54027572 |
| AT5G24790 | NA | Protein of unk | 0,00219837 | -4,76024554 | NA | Protein of unk | 0,00254217 | -1,22991427 |
| AT2G26940 | NA | C2H2-type zin | 9,78E-05 | -4,72842245 | NA | C2H2-type zin | 0,01505899 | -1,28371875 |
| AT4G35733 | NA | F-box family p | 0,00079487 | -4,64449998 | NA | F-box family p | 0,03318374 | -1,58994186 |
| AT2G39590 | NA | Ribosomal prc | 0,00039488 | -4,58457564 | NA | Ribosomal prc | 0,02159733 | -1,49988447 |
| AT2G03200 | NA | Eukaryotic asp | 0,0037273 | -4,54821075 | NA | Eukaryotic asp | 5,46E-06 | -1,315012 |
| AT3G20450 | NA | B-cell recepto | 0,01043253 | -4,51405643 | NA | B-cell recepto | 0,00165356 | -1,13300855 |
| AT3G49020 | NA | FBD F-box anc | 0,00259641 | -4,48411426 | NA | FBD F-box anc | 0,00945837 | -1,78592832 |
| AT1G17710 | NA | Pyridoxal pho: | 0,00562966 | -4,48402523 | NA | Pyridoxal pho: | 0,03368606 | -1,11401255 |
| AT3G11340 | NA | UDP-Glycosylt | 0,00048644 | -4,45228295 | NA | UDP-Glycosylt | 0,00180201 | -3,087659 |
| AT1G73220 | 01-oct | organic cation | 0,00025508 | -4,44143559 | 01-oct | organic cation | 0,00481846 | -1,94206844 |
| AT3G17320 | NA | F-box and ass | 0,00142522 | -4,42767193 | NA | F-box and ass | 0,00826225 | -2,03698874 |
| AT1G75920 | NA | GDSL-like Lipa | 0,0030117 | -4,40703202 | NA | GDSL-like Lipa | 1,50E-05 | -2,61532278 |
| AT5G35110 | na | na | 0,00683483 | -4,3717975 | NA | NA | 0,01865457 | -1,78878449 |
| AT3G15400 | ATA20 | anther 20 | 0,00112396 | -4,34037396 | ATA20 | anther 20 | 3,23E-06 | -1,06584795 |
| AT5G47635 | NA | Pollen Ole e 1 | 0,0002565 | -4,32827985 | NA | Pollen Ole e 1 | 0,00329603 | -1,06958448 |
| AT1G64030 | ATSRP3 | serpin 3 | 0,00047377 | -4,29293118 | ATSRP3 | serpin 3 | 0,01966359 | -1,10021647 |
| AT5G60140 | NA | AP2/B3-like tr | 0,00826872 | -4,18019658 | NA | AP2/B3-like tr | 3,83E-05 | -2,24522724 |
| AT1G77210 | AtSTP14 | sugar transpo | 0,01381203 | -4,16802847 | AtSTP14 | sugar transpo | 4,59E-05 | -1,23996833 |
| AT3G07070 | NA | Protein kinase | 0,0060625 | -4,15813987 | NA | Protein kinase | 0,025435 | -1,5268683 |
| AT2G21730 | ATCAD2 | cinnamyl alco | 0,0002177 | -4,13445913 | ATCAD2 | cinnamyl alco | 0,00423525 | -1,50486572 |
| AT1G74540 | CYP98A8 | cytochrome P | 0,02114149 | -4,12566116 | CYP98A8 | cytochrome P | 1,58E-08 | -2,00407148 |
| AT4G37520 | NA | Peroxidase su | 4,25E-05 | -4,12163536 | NA | Peroxidase su | 0,00534915 | -1,93904265 |
| AT1G22760 | PAB3 | poly(A) bindin | 0,0009365 | -4,0538499 | PAB3 | poly(A) bindin | 0,00789896 | -1,37002472 |
| AT1G02520 | PGP11 | P-glycoprotei | 0,00659757 | -4,02762436 | PGP11 | P-glycoprotei | 2,42E-05 | -1,58587083 |
| AT4G04760 | NA | Major facilitat | 0,02518596 | -4,00967967 | NA | Major facilitat | 0,00092904 | -1,64230299 |
| AT1G71160 | KCS7 | 3-ketoacyl-Co | 3,58E-05 | -3,92911799 | KCS7 | 3-ketoacyl-Co | 0,00013259 | -1,04657395 |
| AT5G59845 | NA | Gibberellin-re | 0,01334239 | -3,9179379 | NA | Gibberellin-re | 5,52E-06 | -1,96705045 |
| AT4G19460 | NA | UDP-Glycosylt | 9,96E-05 | -3,87599695 | NA | UDP-Glycosylt | 0,00028378 | -1,97687571 |
| AT4G26390 | NA | Pyruvate kina: | 0,00053143 | -3,84489071 | NA | Pyruvate kina: | 0,00826948 | -1,96480924 |
| AT1G13140 | CYP86C3 | cytochrome P | 0,00588785 | -3,76043301 | CYP86C3 | cytochrome P | 2,93E-07 | -1,49863671 |
| AT2G34010 | na | na | 0,00416028 | -3,70722796 | NA | NA | 0,01757992 | -1,30651732 |
| AT4G18550 | NA | alpha/beta-Hy | 0,02778627 | -3,67777307 | NA | alpha/beta-Hy | 4,78E-06 | -1,80003621 |

| | | | | | | | | |
|---|---|---|---|---|---|---|---|---|
| AT1G23060 | na | na | 0,03504672 | -3,63676925 | NA | NA | 0,003387 | -1,84022581 |
| AT5G49070 | KCS21 | 3-ketoacyl-Co. | 0,00911659 | -3,54374098 | KCS21 | 3-ketoacyl-Co. | 2,12E-05 | -1,86978761 |
| AT5G66690 | UGT72E2 | UDP-Glycosylt | 0,02300443 | -3,5332683 | UGT72E2 | UDP-Glycosylt | 0,00010233 | -1,39736855 |
| AT2G20470 | NA | AGC (cAMP-de | 5,63E-06 | -3,52781104 | NA | AGC (cAMP-de | 9,27E-05 | -1,26921672 |
| AT1G49120 | NA | Integrase-type | 0,00266899 | -3,5143749 | NA | Integrase-type | 0,00351916 | -2,08639138 |
| AT2G16750 | NA | Protein kinase | 0,00259706 | -3,50945682 | NA | Protein kinase | 0,00015171 | -1,23038129 |
| AT4G04460 | NA | Saposin-like a: | 0,0015021 | -3,47658986 | NA | Saposin-like a: | 1,28E-07 | -1,66014345 |
| AT5G40040 | NA | 60S acidic ribo | 0,00043747 | -3,43721859 | NA | 60S acidic ribo | 0,00791398 | -1,42200425 |
| AT5G17590 | NA | Putative mem | 0,00149772 | -3,39839178 | NA | Putative mem | 0,02046492 | -1,53978441 |
| AT4G19645 | NA | TRAM LAG1 ar | 0,00798881 | -3,3291667 | NA | TRAM LAG1 ar | 1,96E-05 | -1,31916768 |
| AT3G42850 | NA | Mevalonate/g | 0,00106814 | -3,26857067 | NA | Mevalonate/g | 1,17E-05 | -1,9540771 |
| AT2G27120 | POL2B | DNA polymera | 0,00013605 | -3,25671961 | POL2B | DNA polymera | 0,00373805 | -1,33267222 |
| AT5G66110 | NA | Heavy metal t | 0,00218105 | -3,21195585 | NA | Heavy metal t | 0,02865194 | -1,50358162 |
| AT2G21237 | na | na | 0,02311175 | -3,11961987 | NA | NA | 0,02954402 | -1,5256179 |
| AT2G21890 | ATCAD3 | cinnamyl alcol | 0,00185985 | -3,02957767 | ATCAD3 | cinnamyl alco | 0,0042695 | -2,09350185 |
| AT2G41040 | NA | S-adenosyl-L-r | 0,01978069 | -2,98872381 | NA | S-adenosyl-L-r | 1,28E-07 | -1,55383429 |
| AT3G15510 | ANAC056 | NAC domain c | 0,00381754 | -2,97681768 | ANAC056 | NAC domain c | 0,0303236 | -2,29869943 |
| AT3G45060 | ATNRT2.6 | high affinity n | 0,01132122 | -2,94487695 | ATNRT2.6 | high affinity n | 1,09E-05 | -2,11751977 |
| AT1G62305 | NA | Core-2/I-bran | 5,37E-06 | -2,93472549 | NA | Core-2/I-bran | 2,88E-05 | -1,13666901 |
| AT4G26830 | NA | O-Glycosyl hy | 0,01070608 | -2,92799529 | NA | O-Glycosyl hy | 0,02954402 | -1,06875317 |
| AT5G59040 | COPT3 | copper transp | 0,00407199 | -2,90124874 | COPT3 | copper transp | 0,03453338 | -1,64362766 |
| AT5G04500 | NA | glycosyltransf | 2,18E-05 | -2,83213465 | NA | glycosyltransf | 0,0197791 | -1,04505056 |
| AT5G39610 | ANAC092 | NAC domain c | 0,00010267 | -2,80518382 | ANAC092 | NAC domain c | 0,04813156 | -1,63153516 |
| AT5G18090 | NA | AP2/B3-like tr | 0,00294066 | -2,78814268 | NA | AP2/B3-like tr | 0,02407326 | -1,26195213 |
| AT3G23840 | NA | HXXXD-type a | 1,28E-06 | -2,72812572 | NA | HXXXD-type a | 1,30E-07 | -1,41359206 |
| AT2G28120 | NA | Major facilitat | 2,76E-05 | -2,6122564 | NA | Major facilitat | 0,00240712 | -1,83533018 |
| AT2G23970 | NA | Class I glutam | 0,03385468 | -2,60846314 | NA | Class I glutam | 0,03394031 | -1,56014719 |
| AT4G01450 | NA | nodulin MtN2 | 0,00068263 | -2,55804789 | NA | nodulin MtN2 | 0,00077926 | -1,97605923 |
| AT4G16560 | NA | HSP20-like cha | 0,00552859 | -2,50563257 | NA | HSP20-like cha | 0,01381826 | 1,6460687 |
| AT2G19110 | ATHMA4 | heavy metal a | 1,80E-05 | -2,4856838 | ATHMA4 | heavy metal a | 1,56E-05 | -1,19899694 |
| AT3G15740 | NA | RING/U-box su | 0,01389775 | -2,433278 | NA | RING/U-box su | 0,04993678 | -1,2391799 |
| AT2G28140 | NA | Protein of unk | 0,00010465 | -2,4255916 | NA | Protein of unk | 0,00022001 | -1,462322 |
| AT5G14180 | MPL1 | Myzus persica | 7,63E-05 | -2,362592 | MPL1 | Myzus persica | 0,00487738 | -1,43696504 |

| | | | | | | | | |
|---|---|---|---|---|---|---|---|---|
| AT4G01430 | NA | nodulin MtN2 | 0,00586826 | -2,33191216 | NA | nodulin MtN2 | 6,42E-07 | -2,0342003 |
| AT4G02380 | AtLEA5 | senescence-as | 0,00056295 | -2,21681208 | AtLEA5 | senescence-as | 0,00153709 | -1,64818306 |
| AT1G78230 | NA | Outer arm dyr | 0,01865004 | -2,1455363 | NA | Outer arm dyr | 0,00051676 | -1,24760983 |
| AT4G17483 | NA | alpha/beta-Hy | 0,02186487 | -2,1399069 | NA | alpha/beta-Hy | 0,0060787 | -1,13742169 |
| AT1G72280 | AERO1 | endoplasmic r | 0,02014346 | -2,13898223 | AERO1 | endoplasmic r | 1,57E-05 | -1,55440438 |
| AT3G28540 | NA | P-loop contair | 9,81E-05 | -2,08957723 | NA | P-loop contair | 0,00535875 | -1,51307567 |
| AT5G58390 | NA | Peroxidase su | 0,04085389 | -2,08067626 | NA | Peroxidase su | 0,00398139 | -1,25839532 |
| AT5G59820 | RHL41 | C2H2-type zin | 0,0278334 | -1,99826096 | RHL41 | C2H2-type zin | 0,00813363 | -1,3183574 |
| AT1G19640 | JMT | jasmonic acid | 0,0086009 | -1,99187281 | JMT | jasmonic acid | 0,00198877 | -1,17483015 |
| AT3G10340 | PAL4 | phenylalanine | 0,01244758 | -1,90511481 | PAL4 | phenylalanine | 0,00809482 | -1,02275254 |
| AT3G45010 | scpl48 | serine carboxy | 0,00131753 | -1,89485481 | scpl48 | serine carboxy | 2,02E-06 | -1,1825794 |
| AT2G16260 | na | na | 0,03097065 | -1,8529974 | NA | NA | 0,03778658 | -1,27777209 |
| AT5G65205 | NA | NAD(P)-bindir | 0,003869 | -1,81749913 | NA | NAD(P)-bindir | 1,18E-05 | -1,19094558 |
| AT4G28530 | anac074 | NAC domain c | 0,00183161 | -1,77140272 | anac074 | NAC domain c | 0,03846371 | -1,17703627 |
| AT1G75490 | NA | Integrase-type | 0,04470253 | -1,71540232 | NA | Integrase-type | 0,03048836 | -1,5072831 |
| AT5G60330 | na | na | 0,04525158 | -1,7030443 | NA | NA | 0,0288758 | -1,26282922 |
| AT5G41130 | NA | Esterase/lipas | 0,00033531 | -1,62385586 | NA | Esterase/lipas | 1,48E-06 | -1,15738799 |
| AT1G18280 | NA | Bifunctional ir | 0,01372898 | -1,55536587 | NA | Bifunctional ir | 0,04362713 | -1,65697635 |
| AT5G15960 | KIN1 | stress-respons | 0,0145838 | -1,5494545 | KIN1 | stress-respons | 0,00011366 | -1,2460604 |
| AT2G31980 | AtCYS2 | PHYTOCYSTAT | 0,02187029 | -1,50619549 | AtCYS2 | PHYTOCYSTAT | 2,70E-05 | -1,14587798 |
| AT4G24040 | ATTRE1 | trehalase 1 | 0,00315617 | -1,4452894 | ATTRE1 | trehalase 1 | 0,00011366 | -1,96807543 |
| AT4G01883 | NA | Polyketide cyc | 0,00520693 | -1,43724573 | NA | Polyketide cyc | 6,02E-05 | -1,01541992 |
| AT1G33700 | NA | Beta-glucosida | 0,00987424 | -1,4324506 | NA | Beta-glucosida | 3,00E-05 | -2,94601323 |
| AT3G47340 | ASN1 | glutamine-dep | 0,01954695 | -1,42010821 | ASN1 | glutamine-dep | 0,00406542 | -2,0823172 |
| AT3G11680 | NA | Aluminium act | 0,04134917 | -1,32307763 | NA | Aluminium act | 0,02601519 | -1,50662235 |
| AT2G38010 | NA | Neutral/alkali | 0,00341376 | -1,25323603 | NA | Neutral/alkali | 1,96E-05 | -1,01810892 |
| AT4G00040 | NA | Chalcone and | 0,0139725 | -1,18513236 | NA | Chalcone and | 1,59E-05 | -1,03040697 |
| AT1G45145 | ATH5 | thioredoxin H- | 0,00066489 | -1,15808035 | ATH5 | thioredoxin H- | 3,41E-05 | -1,31534362 |
| AT2G05160 | NA | CCCH-type zin | 0,02155031 | -1,15517775 | NA | CCCH-type zin | 0,01118712 | -1,1689636 |
| AT1G10070 | ATBCAT-2 | branched-cha | 0,00602487 | -1,10341844 | ATBCAT-2 | branched-cha | 0,00042003 | -1,6693924 |
| AT3G23430 | ATPHO1 | phosphate 1 | 0,00344879 | -1,08882341 | ATPHO1 | phosphate 1 | 0,01032082 | -1,7648411 |
| AT3G62860 | NA | alpha/beta-Hy | 0,00397392 | -1,03945925 | NA | alpha/beta-Hy | 0,00826225 | -1,05799246 |
| AT2G05380 | GRP3S | glycine-rich pr | 0,04166133 | -1,0271543 | GRP3S | glycine-rich pr | 2,93E-07 | -1,36346429 |

| | | | | | | | | |
|---|---|---|---|---|---|---|---|---|
| AT1G52890 | ANAC019 | NAC domain c | 0,01336893 | -1,02424389 | ANAC019 | NAC domain c | 0,00051176 | -1,26208218 |
| AT5G41750 | NA | Disease resist: | 0,02953665 | -1,02238824 | NA | Disease resist: | 0,00024021 | -2,50607553 |
| AT4G10260 | NA | pfkB-like carb | 0,03241584 | 1,3304106 | NA | pfkB-like carb | 0,00756358 | -1,93583629 |
| AT5G03790 | ATHB51 | homeobox 51 | 0,00022518 | 1,42927598 | ATHB51 | homeobox 51 | 0,00153709 | 1,05905707 |
| AT1G62975 | NA | basic helix-loc | 0,02356472 | 1,48843974 | NA | basic helix-loc | 0,00050877 | 2,60270689 |
| AT2G32530 | ATCSLB03 | cellulose syntl | 2,43E-06 | 1,7912819 | ATCSLB03 | cellulose syntl | 0,00673568 | 1,01253599 |
| AT1G70890 | MLP43 | MLP-like prot | 0,00067961 | 2,41664737 | MLP43 | MLP-like prot | 2,32E-05 | 2,9855585 |

Deregulated gene in the sep1sep2sep3 mutant versus WT

| GeneID | Symbol | GeneName | FDR | logFC |
|---|---|---|---|---|
| AT4G19240 | na | na | 2,40E-10 | 11,4107182 |
| AT5G15360 | na | na | 2,40E-10 | 10,4981351 |
| AT1G54040 | ESP | epithiospecifi | 2,40E-10 | 6,43054158 |
| AT5G25980 | BGLU37 | glucoside gluc | 1,40E-09 | -4,88837969 |
| AT4G15242 | na | na | 3,07E-09 | 13,2175448 |
| AT3G42658.1 | na | na | 6,28E-09 | 10,519266 |
| AT4G08093 | na | na | 1,37E-08 | 8,32804439 |
| AT5G17100 | NA | Cystatin/mon | 2,01E-08 | 5,73802105 |
| AT1G60020.1 | na | na | 2,69E-08 | 13,492982 |
| AT5G34871 | na | na | 2,69E-08 | 6,46324234 |
| AT4G20480 | NA | Putative endo | 3,30E-08 | -5,80547958 |
| AT5G24820 | NA | Eukaryotic asp | 5,01E-08 | -2,92751986 |
| AT4G19239 | na | na | 5,11E-08 | 12,330033 |
| AT1G65150 | NA | TRAF-like fam | 6,04E-08 | -6,78096879 |
| AT1G02900 | ATRALF1 | rapid alkaliniz | 6,04E-08 | -3,59503763 |
| AT1G71920 | HISN6B | HISTIDINE BIO | 6,81E-08 | 11,224429 |
| AT1G66850 | NA | Bifunctional ir | 6,81E-08 | -5,70653458 |
| AT4G19510 | NA | Disease resista | 7,20E-08 | -5,46714577 |
| AT1G57860 | NA | Translation pr | 7,87E-08 | -2,86797463 |
| AT1G08320 | bZIP21 | bZIP transcrip | 9,67E-08 | -2,69151725 |
| AT1G60710 | ATB2 | NAD(P)-linked | 9,67E-08 | -2,18740204 |
| AT4G13495 | na | na | 1,04E-07 | 2,68669862 |
| AT1G73490 | NA | RNA-binding ( | 1,71E-07 | -9,44372561 |
| AT5G17090 | NA | Cystatin/mon | 1,71E-07 | -5,68889907 |
| AT1G68945 | na | na | 1,71E-07 | -5,18137928 |
| AT4G10990.1 | na | na | 1,86E-07 | 10,3524475 |
| AT5G41090 | anac095 | NAC domain c | 1,91E-07 | -4,34718915 |
| AT4G35420 | DRL1 | dihydroflavon | 1,91E-07 | -3,63511143 |
| AT2G16910 | AMS | basic helix-loc | 1,93E-07 | -3,45218033 |
| AT5G32460 | NA | Transcriptiona | 2,03E-07 | 2,94167777 |
| AT4G27330 | NZZ | sporocyteless | 2,03E-07 | -2,81653778 |

| | | | | |
|---|---|---|---|---|
| AT1G06640 | NA | 2-oxoglutarat | 2,04E-07 | -2,65085689 |
| AT5G14760 | AO | L-aspartate ox | 2,04E-07 | -2,32864373 |
| AT2G35810 | na | na | 2,16E-07 | 1,9184943 |
| AT5G49360 | ATBXL1 | beta-xylosidas | 2,16E-07 | -2,71394576 |
| AT4G10850 | NA | Nodulin MtN3 | 2,60E-07 | -2,48811779 |
| AT1G44970 | NA | Peroxidase su | 2,81E-07 | -2,3979188 |
| AT1G54260 | NA | winged-helix I | 2,93E-07 | -5,82185925 |
| AT5G53190 | NA | Nodulin MtN3 | 2,99E-07 | -3,30442371 |
| AT3G25260 | NA | Major facilitat | 2,99E-07 | -2,98781089 |
| AT4G16215 | na | na | 3,06E-07 | 8,61523329 |
| AT1G48180 | na | na | 3,06E-07 | 2,74910975 |
| AT1G63880 | NA | Disease resist | 3,06E-07 | -11,8665984 |
| AT2G21430 | NA | Papain family | 3,06E-07 | -3,28267269 |
| AT4G29980 | na | na | 3,06E-07 | -2,85769024 |
| AT1G65040 | NA | RING/U-box su | 3,06E-07 | -1,90110731 |
| AT4G32208 | NA | heat shock pr | 3,08E-07 | 3,89546986 |
| AT2G30140 | NA | UDP-Glycosylt | 3,08E-07 | 2,67912582 |
| AT1G61070 | LCR66 | low-molecula | 3,08E-07 | -6,17444146 |
| AT5G44540 | NA | Tapetum spec | 3,08E-07 | -4,60465156 |
| AT5G52160 | NA | Bifunctional in | 3,08E-07 | -3,27511327 |
| AT5G50400 | ATPAP27 | purple acid ph | 3,08E-07 | -1,71215201 |
| AT1G63320 | NA | Pentatricopep | 3,22E-07 | 10,3542427 |
| AT5G47590 | NA | Heat shock pr | 3,22E-07 | -2,06008888 |
| AT1G27710 | NA | Glycine-rich p | 3,32E-07 | -3,02988504 |
| AT5G11230 | NA | Nucleotide-su | 3,32E-07 | -2,05935602 |
| AT4G02540 | NA | Cysteine/Histi | 3,47E-07 | -8,54519962 |
| AT5G15140 | NA | Galactose mu | 3,47E-07 | -3,53059883 |
| AT3G17675 | NA | Cupredoxin su | 3,56E-07 | -3,26408188 |
| AT5G27890 | na | na | 3,66E-07 | -2,47506757 |
| AT5G35935.1 | na | na | 3,73E-07 | -13,0090939 |
| AT2G35690 | ACX5 | acyl-CoA oxida | 4,33E-07 | 1,59035843 |
| AT4G19500 | NA | nucleoside-tri | 4,33E-07 | -13,776917 |
| AT3G57370 | NA | Cyclin family p | 4,33E-07 | -3,80667029 |

| | | | | |
|---|---|---|---|---|
| AT1G27270 | NA | Paired amphip | 4,40E-07 | -3,90600404 |
| AT1G62940 | ACOS5 | acyl-CoA syntl | 4,50E-07 | -4,34099662 |
| AT4G11310 | NA | Papain family | 4,97E-07 | 3,06428619 |
| AT1G07780 | PAI1 | phosphoribosy | 5,01E-07 | 1,63999106 |
| AT3G48690 | ATCXE12 | alpha/beta-Hy | 5,06E-07 | -2,16863797 |
| AT4G14080 | MEE48 | O-Glycosyl hy | 5,29E-07 | -3,6567112 |
| AT5G60080 | NA | Protein kinase | 5,53E-07 | -3,44970898 |
| AT5G59390 | NA | XH/XS domair | 5,54E-07 | -1,97784003 |
| AT5G40260 | NA | Nodulin MtN3 | 5,99E-07 | -3,48716741 |
| AT5G56110 | AtMYB103 | myb domain p | 6,01E-07 | -3,91675673 |
| AT1G58380 | XW6 | Ribosomal prc | 6,21E-07 | 1,5175962 |
| AT1G73050 | NA | Glucose-meth | 6,63E-07 | -2,32255534 |
| AT4G01590 | na | na | 7,40E-07 | -1,63727321 |
| AT5G07830 | AtGUS2 | glucuronidase | 7,61E-07 | -2,39811129 |
| AT5G60090 | NA | Protein kinase | 7,82E-07 | -3,37441859 |
| AT4G13630 | NA | Protein of unk | 8,08E-07 | -3,47956844 |
| AT5G62065 | NA | Bifunctional ir | 8,08E-07 | -3,07153306 |
| AT4G23010 | ATUTR2 | UDP-galactose | 8,08E-07 | -2,18364537 |
| AT5G27010 | NA | ARM repeat su | 9,03E-07 | 3,75457288 |
| AT5G61110 | NA | zinc ion bindir | 9,43E-07 | -4,15416577 |
| AT1G04880 | NA | HMG (high mc | 9,43E-07 | -2,17665664 |
| AT5G17900 | NA | microfibrillar-a | 9,43E-07 | -1,65866972 |
| AT5G02660 | NA | Protein with d | 9,59E-07 | -4,12793461 |
| AT1G19230 | NA | Riboflavin syn | 1,05E-06 | -2,62024707 |
| AT1G75790 | sks18 | SKU5 similar 1 | 1,08E-06 | -2,74543421 |
| AT5G56030 | AtHsp90.2 | heat shock prc | 1,08E-06 | -1,72690998 |
| AT1G64710 | NA | GroES-like zin | 1,14E-06 | 2,00634893 |
| AT2G38690 | na | na | 1,16E-06 | -4,16236003 |
| AT5G37410 | NA | Family of unkı | 1,28E-06 | -4,24951214 |
| AT1G65370 | NA | TRAF-like fam | 1,28E-06 | -3,83612823 |
| AT3G52130 | NA | Bifunctional ir | 1,28E-06 | -3,35813705 |
| AT3G23840 | NA | HXXXD-type a | 1,28E-06 | -2,72812572 |
| AT2G29780 | NA | Galactose oxid | 1,29E-06 | -4,96108797 |

| | | | | |
|---|---|---|---|---|
| AT3G59530 | LAP3 | Calcium-depe | 1,29E-06 | -3,77095092 |
| AT5G40390 | SIP1 | Raffinose synt | 1,31E-06 | 2,284415 |
| AT5G65530 | NA | Protein kinase | 1,31E-06 | 1,86019771 |
| AT2G31210 | NA | basic helix-loc | 1,31E-06 | -3,06198336 |
| AT5G14840 | na | na | 1,31E-06 | -2,70534988 |
| AT1G19460 | NA | Galactose oxi | 1,37E-06 | -3,50757547 |
| AT1G06520 | ATGPAT1 | glycerol-3-pho | 1,37E-06 | -2,41843937 |
| AT1G06170 | NA | basic helix-loc | 1,37E-06 | -2,23830844 |
| AT3G18610 | ATNUC-L2 | nucleolin like | 1,40E-06 | -4,10282629 |
| AT5G53750 | NA | CBS domain-c | 1,41E-06 | -2,25544707 |
| AT5G17890 | CHS3 | DA1-related p | 1,42E-06 | -12,9991541 |
| AT4G08035 | na | na | 1,43E-06 | -1,67944056 |
| AT1G54240 | NA | winged-helix I | 1,50E-06 | -3,11401673 |
| AT1G71990 | ATFT4 | fucosyltransfe | 1,53E-06 | 1,4990054 |
| AT1G52905 | na | na | 1,54E-06 | -11,9738914 |
| AT2G41440 | na | na | 1,54E-06 | -9,57683739 |
| AT1G33430 | NA | Galactosyltrar | 1,54E-06 | -2,96835611 |
| AT5G26280 | NA | TRAF-like fam | 1,60E-06 | 6,91216155 |
| AT4G16950 | RPP5 | Disease resist | 1,60E-06 | -3,63351793 |
| AT5G61260 | NA | Plant calmodu | 1,63E-06 | -3,03880761 |
| AT2G31220 | NA | basic helix-loc | 1,63E-06 | -2,2630688 |
| AT1G05690 | BT3 | BTB and TAZ c | 1,65E-06 | -2,44173496 |
| AT1G06280 | LBD2 | LOB domain-c | 1,66E-06 | -4,27441832 |
| AT5G26642.1 | na | na | 1,67E-06 | 9,77640385 |
| AT3G60020 | ASK5 | SKP1-like 5 | 1,67E-06 | -3,78180172 |
| AT5G62080 | NA | Bifunctional ir | 1,67E-06 | -3,37086944 |
| AT1G74870 | NA | RING/U-box su | 1,67E-06 | -3,25793494 |
| AT1G69500 | CYP704B1 | cytochrome P | 1,71E-06 | -3,5894768 |
| AT5G21900 | NA | RNI-like super | 1,78E-06 | -1,59188238 |
| AT5G63900 | NA | Acyl-CoA N-ac | 1,81E-06 | -5,29166706 |
| AT1G04645 | NA | Plant self-inco | 1,81E-06 | -3,27204566 |
| AT1G28710 | NA | Nucleotide-di | 1,81E-06 | -1,96469124 |
| AT1G68540 | NA | NAD(P)-bindir | 1,87E-06 | -3,12759074 |

| | | | | |
|---|---|---|---|---|
| AT2G36890 | ATMYB38 | Duplicated ho | 1,87E-06 | -2,17486176 |
| AT1G74130 | NA | Rhomboid-rel | 1,93E-06 | -2,9611208 |
| AT1G27260 | NA | Paired amphip | 1,98E-06 | -4,86113442 |
| AT5G24670 | NA | Cytidine/deox | 1,98E-06 | -2,03332877 |
| AT1G02980 | ATCUL2 | cullin 2 | 1,99E-06 | -3,50900082 |
| AT3G11000 | NA | DCD (Develop | 1,99E-06 | -1,23565869 |
| AT1G72130 | NA | Major facilitat | 2,03E-06 | 1,99188356 |
| AT4G36140 | NA | disease resista | 2,03E-06 | -9,25886951 |
| AT3G04960 | NA | Domain of unl | 2,03E-06 | -1,47287224 |
| AT2G42510 | na | na | 2,06E-06 | -1,87457024 |
| AT3G26140 | NA | Cellulase (glyc | 2,08E-06 | -3,48613646 |
| AT4G34850 | LAP5 | Chalcone and | 2,09E-06 | -3,97807223 |
| AT1G60240 | NA | NAC (No Apica | 2,09E-06 | -3,96422545 |
| AT1G22015 | DD46 | Galactosyltrar | 2,09E-06 | -2,44836515 |
| AT4G14965 | AtMAPR4 | membrane-as | 2,22E-06 | -1,42539611 |
| AT1G72230 | NA | Cupredoxin su | 2,24E-06 | -1,9351417 |
| AT2G33010 | NA | Ubiquitin-assc | 2,26E-06 | -2,96229363 |
| AT5G10400 | NA | Histone super | 2,28E-06 | -3,5043319 |
| AT1G03390 | NA | HXXXD-type a | 2,30E-06 | -4,60760357 |
| AT5G18320 | NA | ARM repeat su | 2,32E-06 | -3,50938127 |
| AT5G51950 | NA | Glucose-meth | 2,39E-06 | -2,81047223 |
| AT5G11130 | NA | Exostosin fam | 2,40E-06 | -5,48029075 |
| AT2G32530 | ATCSLB03 | cellulose syntl | 2,43E-06 | 1,7912819 |
| AT4G10120 | ATSPS4F | Sucrose-phosp | 2,49E-06 | -2,12091186 |
| AT5G39770 | NA | Restriction en | 2,55E-06 | 9,15609254 |
| AT1G60960 | ATIRT3 | iron regulated | 2,72E-06 | -1,8000294 |
| AT4G16270 | NA | Peroxidase su | 2,82E-06 | -1,72828358 |
| AT5G43110 | APUM14 | pumilio 14 | 2,95E-06 | -5,2637121 |
| AT1G02510 | ATKCO4 | Outward recti | 2,95E-06 | -3,86929683 |
| AT3G07450 | NA | Bifunctional ir | 2,95E-06 | -3,45532246 |
| AT5G08250 | NA | Cytochrome P | 2,95E-06 | -2,79304504 |
| AT5G44700 | EDA23 | Leucine-rich r | 2,95E-06 | -2,76902369 |
| AT5G56910 | NA | Proteinase inh | 2,95E-06 | -2,73546452 |

| | | | | |
|---|---|---|---|---|
| AT2G24800 | NA | Peroxidase su | 2,95E-06 | -2,5162442 |
| AT4G34470 | ASK12 | SKP1-like 12 | 2,96E-06 | -10,1492656 |
| AT5G37400 | NA | Family of unkr | 2,96E-06 | -3,1269464 |
| AT3G10430 | NA | F-box and ass | 2,96E-06 | -3,00256964 |
| AT4G15910 | ATDI21 | drought-induc | 3,04E-06 | -2,79634569 |
| AT5G43330 | NA | Lactate/malat | 3,04E-06 | -1,7132244 |
| AT2G33000 | NA | ubiquitin-asso | 3,19E-06 | -2,47879006 |
| AT5G53700 | NA | RNA-binding ( | 3,20E-06 | -5,02850247 |
| AT1G61590 | NA | Protein kinase | 3,28E-06 | -4,07308175 |
| AT4G33870 | NA | Peroxidase su | 3,28E-06 | -2,63557429 |
| AT3G49120 | ATPCB | peroxidase CB | 3,30E-06 | 1,93842923 |
| AT3G12143 | na | na | 3,30E-06 | -5,13912168 |
| AT1G61340 | NA | F-box family p | 3,30E-06 | -2,86328093 |
| AT5G55970 | NA | RING/U-box su | 3,30E-06 | -2,67573495 |
| AT5G46510 | NA | Disease resist | 3,35E-06 | -13,0556139 |
| AT1G01400 | na | na | 3,39E-06 | -3,75932345 |
| AT3G60010 | ASK13 | SKP1-like 13 | 3,39E-06 | -3,58257124 |
| AT1G49330 | NA | hydroxyprolin | 3,39E-06 | -2,78827183 |
| AT5G43420 | NA | RING/U-box su | 3,47E-06 | -4,69244116 |
| AT2G17170 | NA | Protein kinase | 3,48E-06 | -3,07295752 |
| AT5G23700 | na | na | 3,48E-06 | -3,03302928 |
| AT1G70440 | SRO3 | similar to RCD | 3,49E-06 | -3,37646325 |
| AT1G76250 | na | na | 3,59E-06 | -2,77223809 |
| AT5G28626.1 | na | na | 3,62E-06 | 2,20258544 |
| AT1G05260 | RCI3 | Peroxidase su | 3,62E-06 | -2,33597645 |
| AT1G02260 | NA | Divalent ion sy | 3,62E-06 | -2,01350206 |
| AT2G35550 | ATBPC7 | basic pentacys | 3,71E-06 | -2,83475524 |
| AT5G61620 | NA | myb-like trans | 3,74E-06 | -4,79387909 |
| AT4G14250 | NA | structural con | 3,74E-06 | -4,46106279 |
| AT3G54450 | NA | Major facilitat | 3,77E-06 | -5,40354679 |
| AT1G58602 | NA | LRR and NB-A | 3,77E-06 | -2,98678964 |
| AT3G20990.1 | na | na | 3,80E-06 | -4,2360936 |
| AT3G55570 | na | na | 3,80E-06 | -4,19538262 |

| | | | | |
|---|---|---|---|---|
| AT3G19580 | AZF2 | zinc-finger prc | 3,80E-06 | -2,81657499 |
| AT1G71770 | PAB5 | poly(A)-bindir | 3,80E-06 | -2,46340921 |
| AT1G63540 | NA | hydroxyprolin | 3,82E-06 | 10,0894271 |
| AT1G02770 | NA | Protein of unk | 3,82E-06 | -5,63832633 |
| AT1G62570 | FMO GS-OX4 | flavin-monoox | 3,82E-06 | -2,40193333 |
| AT1G77110 | PIN6 | Auxin efflux ca | 3,84E-06 | 1,19745371 |
| AT5G37130 | NA | Protein preny | 3,84E-06 | 1,14652237 |
| AT5G56380 | NA | F-box/RNI-like | 3,86E-06 | -12,7417315 |
| AT1G80990 | NA | XH domain-co | 3,91E-06 | -3,72494503 |
| AT5G21150 | AGO9 | Argonaute far | 3,91E-06 | -2,16538087 |
| AT1G22090 | emb2204 | Protein of unk | 3,94E-06 | -4,62023201 |
| AT4G08110.1 | na | na | 3,99E-06 | -12,5968221 |
| AT1G76550 | NA | Phosphofructo | 3,99E-06 | -1,13785133 |
| AT2G46450 | ATCNGC12 | cyclic nucleoti | 4,03E-06 | -2,19852328 |
| AT3G30122 | na | na | 4,04E-06 | 3,20012729 |
| AT1G69480 | NA | EXS (ERD1/XP | 4,04E-06 | 2,18160434 |
| AT5G06839 | bZIP65 | bZIP transcrip | 4,08E-06 | -1,83615988 |
| AT4G35110 | NA | Arabidopsis pl | 4,12E-06 | -2,51745615 |
| AT5G40950 | RPL27 | ribosomal pro | 4,14E-06 | -1,04361187 |
| AT5G46380 | NA | Kinase-relatec | 4,54E-06 | -3,78789494 |
| AT5G15480 | NA | C2H2-type zin | 4,54E-06 | -3,00436137 |
| AT2G13547 | na | na | 4,64E-06 | 9,72547685 |
| AT1G61430 | NA | S-locus lectin | 4,66E-06 | 1,8498899 |
| AT5G09570 | NA | Cox19-like CH | 4,66E-06 | -4,23126393 |
| AT2G35080 | NA | nucleotide bir | 4,72E-06 | -3,4089863 |
| AT3G58780 | AGL1 | K-box region a | 4,72E-06 | -2,22869916 |
| AT5G28350 | NA | Quinoprotein | 4,82E-06 | -5,50946878 |
| AT4G18440 | NA | L-Aspartase-lil | 4,87E-06 | 1,25215141 |
| AT5G28919 | na | na | 4,87E-06 | -12,5829236 |
| AT2G44590 | ADL1D | DYNAMIN-like | 4,87E-06 | -2,8901524 |
| AT1G58280 | NA | Phosphoglyce | 4,87E-06 | -2,78050918 |
| AT1G28375 | na | na | 4,92E-06 | -3,51459839 |
| AT2G42470 | NA | TRAF-like fam | 4,95E-06 | -10,0276973 |

| | | | | |
|---|---|---|---|---|
| AT4G15620 | NA | Uncharacteris | 4,95E-06 | -8,48258503 |
| AT5G63750 | ARI13 | RING/U-box su | 4,95E-06 | -2,95101885 |
| AT1G52260 | ATPDI3 | PDI-like 1-5 | 4,95E-06 | -2,83994961 |
| AT4G16710 | NA | glycosyltransf | 4,95E-06 | -2,44937175 |
| AT3G29156.1 | na | na | 4,96E-06 | -5,24864263 |
| AT1G44800 | NA | nodulin MtN2 | 5,10E-06 | -1,9771692 |
| AT5G15720 | GLIP7 | GDSL-motif lip | 5,16E-06 | -2,02334548 |
| AT2G32750 | NA | Exostosin fam | 5,19E-06 | -3,55558393 |
| AT1G69120 | AGL7 | K-box region a | 5,27E-06 | 1,09876623 |
| AT1G61370 | NA | S-locus lectin | 5,31E-06 | 1,93997302 |
| AT1G62305 | NA | Core-2/I-bran | 5,37E-06 | -2,93472549 |
| AT5G17860 | CAX7 | calcium excha | 5,37E-06 | -2,49259842 |
| AT2G40960 | NA | Single-strande | 5,47E-06 | 2,18022192 |
| AT1G18960 | NA | myb-like HTH | 5,47E-06 | -3,67816534 |
| AT2G20470 | NA | AGC (cAMP-de | 5,63E-06 | -3,52781104 |
| AT5G06520 | NA | SWAP (Suppre | 5,66E-06 | -5,14325546 |
| AT1G19190 | NA | alpha/beta-Hy | 5,84E-06 | -3,30837421 |
| AT1G66060 | NA | Family of unkr | 5,90E-06 | -4,58115427 |
| AT1G62660 | NA | Glycosyl hydro | 5,98E-06 | 1,46655249 |
| AT2G46660 | CYP78A6 | cytochrome P | 6,07E-06 | -3,07305616 |
| AT4G14905 | NA | Galactose oxic | 6,15E-06 | -12,6712972 |
| AT3G12145 | FLOR1 | Leucine-rich re | 6,15E-06 | -1,60289791 |
| AT5G23820 | NA | MD-2-related | 6,17E-06 | 1,63041039 |
| AT5G57720 | NA | AP2/B3-like tr | 6,17E-06 | -2,20129018 |
| AT1G52030 | F-ATMBP | myrosinase-bi | 6,32E-06 | -1,70498795 |
| AT3G58340 | NA | TRAF-like fam | 6,34E-06 | -5,34619854 |
| AT5G25040 | NA | Major facilitat | 6,34E-06 | -3,92698841 |
| AT1G23810 | NA | Paired amphip | 6,36E-06 | -3,12677098 |
| AT4G10980.1 | na | na | 6,59E-06 | 9,62605496 |
| AT5G44330 | NA | Tetratricopept | 6,62E-06 | -2,94293586 |
| AT5G23190 | CYP86B1 | cytochrome P | 6,64E-06 | -3,29283609 |
| AT5G28913.1 | na | na | 6,92E-06 | -12,48869 |
| AT5G25820 | NA | Exostosin fam | 6,94E-06 | -2,98240075 |

| | | | | |
|---|---|---|---|---|
| AT4G30720 | NA | FAD/NAD(P)-b | 6,95E-06 | 1,2339931 |
| AT4G20340 | NA | Transcription | 6,95E-06 | -1,49723328 |
| AT2G24960 | na | na | 7,03E-06 | 1,31083804 |
| AT4G06598 | na | na | 7,33E-06 | 1,03934082 |
| AT1G68200 | NA | Zinc finger C-x | 7,33E-06 | -2,54657499 |
| AT2G35310 | NA | Transcriptiona | 7,33E-06 | -1,47976706 |
| AT1G75110 | RRA2 | Nucleotide-di | 7,34E-06 | -1,51660028 |
| AT3G57960 | NA | Emsy N Termi | 7,48E-06 | -3,16866958 |
| AT4G19230 | CYP707A1 | cytochrome P | 7,51E-06 | -1,46594419 |
| AT1G23800 | ALDH2B | aldehyde deh | 7,54E-06 | -1,87452685 |
| AT3G29644 | na | na | 7,62E-06 | 2,2057728 |
| AT5G56870 | BGAL4 | beta-galactosi | 7,71E-06 | -1,94179026 |
| AT5G02100 | ORP3A | Oxysterol-bin | 7,71E-06 | -1,73159238 |
| AT3G57390 | AGL18 | AGAMOUS-lik | 7,74E-06 | -1,91956264 |
| AT4G24420 | NA | RNA-binding ( | 7,96E-06 | 2,75913523 |
| AT2G47485 | na | na | 7,96E-06 | -2,77847731 |
| AT5G47260 | NA | ATP binding;G | 8,01E-06 | 10,3910297 |
| AT3G58140 | NA | phenylalanyl-t | 8,01E-06 | 1,26924423 |
| AT1G27045 | NA | Homeobox-leu | 8,01E-06 | -4,86414703 |
| AT4G09455.1 | na | na | 8,01E-06 | -4,78030812 |
| AT1G63230 | NA | Tetratricopep | 8,01E-06 | -3,89342991 |
| AT1G51670 | na | na | 8,01E-06 | -3,46722844 |
| AT3G08750 | NA | F-box and ass | 8,01E-06 | -3,46435861 |
| AT5G56510 | APUM12 | pumilio 12 | 8,01E-06 | -2,93279903 |
| AT1G67810 | SUFE2 | sulfur E2 | 8,19E-06 | -3,72323767 |
| AT4G24130 | NA | Protein of unk | 8,26E-06 | -1,89315529 |
| AT2G32235 | na | na | 8,61E-06 | -1,68885484 |
| AT3G10580 | NA | Homeodomai | 8,81E-06 | -3,56789248 |
| AT1G62540 | FMO GS-OX2 | flavin-monoo | 8,81E-06 | -1,5125804 |
| AT3G13890 | ATMYB26 | myb domain p | 8,83E-06 | -4,91955513 |
| AT1G15190 | NA | Fasciclin-like a | 8,83E-06 | -1,95974227 |
| AT4G24120 | ATYSL1 | YELLOW STRIP | 8,96E-06 | -1,6862912 |
| AT5G37420 | NA | Family of unk | 9,02E-06 | -2,90954862 |

| | | | | |
|---|---|---|---|---|
| AT2G34220 | MEE20 | Protein with d | 9,17E-06 | -2,19703195 |
| AT5G55320 | NA | MBOAT (mem | 9,40E-06 | -5,60556914 |
| AT3G50940 | NA | P-loop contair | 9,42E-06 | -4,77003425 |
| AT4G10690.1 | na | na | 9,65E-06 | -12,2805344 |
| AT5G44410 | NA | FAD-binding B | 9,75E-06 | -1,56008549 |
| AT4G15440 | CYP74B2 | hydroperoxide | 9,82E-06 | 1,68112717 |
| AT2G29770 | NA | Galactose oxid | 1,02E-05 | -4,2754781 |
| AT4G16230 | NA | GDSL-like Lipa | 1,03E-05 | -4,83915736 |
| AT5G44500 | NA | Small nuclear | 1,03E-05 | -1,68312667 |
| AT2G23510 | SDT | spermidine di: | 1,05E-05 | -4,47974326 |
| AT2G42140 | NA | VQ motif-cont | 1,05E-05 | -3,27804291 |
| AT5G52000 | IMPA-8 | importin alpha | 1,06E-05 | -4,02389624 |
| AT2G34210 | NA | Transcription | 1,06E-05 | -4,01513018 |
| AT1G19470 | NA | Galactose oxid | 1,06E-05 | -3,7968627 |
| AT1G47250 | PAF2 | 20S proteasor | 1,06E-05 | -1,15071827 |
| AT1G56030 | NA | RING/U-box su | 1,08E-05 | -5,80518106 |
| AT5G14830.1 | na | na | 1,08E-05 | -5,75739386 |
| AT1G61440 | NA | S-locus lectin | 1,14E-05 | 3,71699987 |
| AT3G28007 | NA | Nodulin MtN3 | 1,17E-05 | -2,61024308 |
| AT1G58848 | NA | Disease resista | 1,21E-05 | -7,78627057 |
| AT1G24230 | NA | Paired amphip | 1,22E-05 | -4,02374109 |
| AT5G42120 | NA | Concanavalin | 1,24E-05 | -2,92903537 |
| AT5G48660 | NA | B-cell recepto | 1,24E-05 | -1,82071186 |
| AT1G65560 | NA | Zinc-binding d | 1,24E-05 | -1,62907524 |
| AT1G62600 | NA | Flavin-binding | 1,24E-05 | -1,51786029 |
| AT3G58280 | NA | Arabidopsis pl | 1,27E-05 | -3,3430381 |
| AT4G24860 | NA | P-loop contair | 1,27E-05 | -2,02493776 |
| AT3G17260.1 | na | na | 1,30E-05 | -6,3904874 |
| AT5G09560 | NA | RNA-binding K | 1,30E-05 | -3,12245346 |
| AT5G08600 | NA | U3 ribonucleo | 1,32E-05 | -3,86254294 |
| AT4G13493 | na | na | 1,34E-05 | 3,55921763 |
| AT1G29640 | NA | Protein of unk | 1,34E-05 | -3,16966071 |
| AT3G21950 | NA | S-adenosyl-L-r | 1,35E-05 | 1,90958047 |

| | | | | |
|---|---|---|---|---|
| AT5G41890 | NA | GDSL-like Lipa | 1,35E-05 | -2,02470142 |
| AT1G74210 | NA | PLC-like phosp | 1,35E-05 | -1,52538718 |
| AT4G36515 | na | na | 1,37E-05 | -7,17636913 |
| AT1G28695 | NA | Nucleotide-dip | 1,40E-05 | -3,57123872 |
| AT5G53950 | ANAC098 | NAC (No Apica | 1,40E-05 | -1,69955985 |
| AT5G56150 | UBC30 | ubiquitin-conj | 1,44E-05 | -1,51329424 |
| AT5G39010 | na | na | 1,47E-05 | -4,9651113 |
| AT5G24920 | AtGDU5 | glutamine dur | 1,55E-05 | -5,16280887 |
| AT1G66590 | ATCOX19-1 | cytochrome c | 1,56E-05 | -5,06699255 |
| AT1G66110 | NA | Family of unkr | 1,56E-05 | -3,79407021 |
| AT5G20370 | NA | serine-rich pro | 1,59E-05 | -4,18749587 |
| AT3G21720 | ICL | isocitrate lyas | 1,60E-05 | -7,88981077 |
| AT1G51965 | ABO5 | ABA Overly-Se | 1,60E-05 | -1,61016655 |
| AT2G32460 | ATM1 | myb domain p | 1,63E-05 | -10,7362691 |
| AT3G59510 | NA | Leucine-rich re | 1,63E-05 | -4,17682422 |
| AT4G07507.1 | na | na | 1,67E-05 | -11,9635312 |
| AT4G26280 | NA | P-loop contair | 1,67E-05 | -4,39833025 |
| AT5G52490 | NA | Fibrillarin fam | 1,73E-05 | -8,58121389 |
| AT3G61680 | NA | alpha/beta-Hy | 1,73E-05 | -2,32974944 |
| AT2G34850 | MEE25 | NAD(P)-bindir | 1,77E-05 | 1,42168945 |
| AT1G67800 | NA | Copine (Calciu | 1,77E-05 | 1,29938789 |
| AT2G26440 | NA | Plant invertase | 1,77E-05 | -1,85940249 |
| AT3G11480 | ATBSMT1 | S-adenosyl-L-r | 1,78E-05 | -5,26847268 |
| AT4G22960 | NA | Protein of unk | 1,79E-05 | -2,07665575 |
| AT2G40920 | NA | F-box and asso | 1,79E-05 | -1,49649711 |
| AT3G58360 | NA | TRAF-like fam | 1,80E-05 | -5,94869003 |
| AT5G26730 | NA | Fasciclin-like a | 1,80E-05 | -4,18988293 |
| AT3G58440 | NA | TRAF-like supe | 1,80E-05 | -3,86595793 |
| AT1G69560 | ATMYB105 | myb domain p | 1,80E-05 | -2,49033068 |
| AT2G19110 | ATHMA4 | heavy metal a | 1,80E-05 | -2,4856838 |
| AT2G43470 | NA | Protein of unk | 1,83E-05 | -3,30484557 |
| AT2G18460 | LCV3 | like COV 3 | 1,83E-05 | -2,48042422 |
| AT4G33270 | CDC20.1 | Transducin far | 1,84E-05 | -1,401665 |

| | | | | |
|---|---|---|---|---|
| AT1G15320 | na | na | 1,86E-05 | -2,8100896 |
| AT3G20975.1 | na | na | 1,86E-05 | -2,44961136 |
| AT3G49830 | NA | P-loop contair | 1,89E-05 | -2,70601722 |
| AT5G23210 | SCPL34 | serine carboxy | 1,89E-05 | -1,33076234 |
| AT4G26255 | na | na | 1,93E-05 | 1,48837604 |
| AT3G58180 | NA | ARM repeat su | 1,93E-05 | 1,14068992 |
| AT2G03160 | ASK19 | SKP1-like 19 | 1,93E-05 | -4,66402678 |
| AT3G06560 | PAPS3 | poly(A) polym | 1,93E-05 | -3,19139061 |
| AT3G30120 | na | na | 1,93E-05 | -2,83986823 |
| AT3G58960 | NA | F-box/RNI-like | 1,94E-05 | -4,33357578 |
| AT5G51620 | NA | Uncharacteris | 1,95E-05 | -11,772339 |
| AT4G02970 | AT7SL-1 | 7SL RNA1 | 1,95E-05 | -2,26402461 |
| AT2G21640 | na | na | 2,00E-05 | -1,71113792 |
| AT3G08900 | RGP | reversibly glyc | 2,01E-05 | -2,43674023 |
| AT4G11320 | NA | Papain family | 2,03E-05 | 1,43913083 |
| AT1G11150.1 | na | na | 2,03E-05 | -10,4674253 |
| AT4G39590 | NA | Galactose oxic | 2,04E-05 | -5,9780989 |
| AT4G15630 | NA | Uncharacteris | 2,04E-05 | -3,25484344 |
| AT5G09750 | HEC3 | basic helix-loc | 2,04E-05 | -3,13663438 |
| AT1G76410 | ATL8 | RING/U-box su | 2,04E-05 | -1,96052195 |
| AT5G41700 | ATUBC8 | ubiquitin conj | 2,04E-05 | -1,12394916 |
| AT1G07660 | NA | Histone super | 2,05E-05 | 1,71433309 |
| AT2G03170 | ASK14 | SKP1-like 14 | 2,07E-05 | -5,29609458 |
| AT3G19390 | NA | Granulin repe | 2,07E-05 | -4,07850411 |
| AT1G64210 | NA | Leucine-rich re | 2,07E-05 | -3,68271903 |
| AT2G21680 | NA | Galactose oxic | 2,10E-05 | -3,55782938 |
| AT2G45750 | NA | S-adenosyl-L-r | 2,10E-05 | -2,67007919 |
| AT2G37770 | NA | NAD(P)-linked | 2,10E-05 | -2,18617027 |
| AT3G32092.1 | na | na | 2,11E-05 | 6,00754478 |
| AT5G51480 | SKS2 | SKU5 similar 2 | 2,17E-05 | -1,62821338 |
| AT4G16870.1 | na | na | 2,18E-05 | -9,68139012 |
| AT1G65160 | NA | Ubiquitin carb | 2,18E-05 | -3,72568876 |
| AT3G09680 | NA | Ribosomal pro | 2,18E-05 | -3,6637585 |

| | | | | |
|---|---|---|---|---|
| AT5G61320 | CYP89A3 | cytochrome P | 2,18E-05 | -3,25304848 |
| AT5G04500 | NA | glycosyltransf | 2,18E-05 | -2,83213465 |
| AT5G48650 | NA | Nuclear trans | 2,18E-05 | -1,00434443 |
| AT4G30110 | ATHMA2 | heavy metal a | 2,25E-05 | -1,57015029 |
| AT3G16415 | na | na | 2,27E-05 | -5,87426548 |
| AT5G18000 | VDD | VERDANDI | 2,27E-05 | -2,95662935 |
| AT1G50400 | NA | Eukaryotic po | 2,30E-05 | -2,66965653 |
| AT4G20900 | MS5 | Tetratricopep | 2,30E-05 | -2,51520017 |
| AT1G63630 | NA | Tetratricopep | 2,30E-05 | -2,18957134 |
| AT4G00750 | NA | S-adenosyl-L- | 2,30E-05 | -1,5347152 |
| AT1G79780 | NA | Uncharacteris | 2,32E-05 | -3,26135415 |
| AT1G06030 | NA | pfkB-like carb | 2,32E-05 | -2,62463814 |
| AT5G43050 | NPQ6 | Protein of unk | 2,34E-05 | -2,75237441 |
| AT4G28580 | ATMGT5 | magnesium tr | 2,35E-05 | -4,06515872 |
| AT2G36660 | PAB7 | poly(A) bindin | 2,38E-05 | -1,95720379 |
| AT1G56510 | ADR2 | Disease resist | 2,39E-05 | -11,263581 |
| AT2G26960 | AtMYB81 | myb domain p | 2,39E-05 | -3,98625148 |
| AT2G27035 | AtENODL20 | early nodulin- | 2,42E-05 | -1,3652351 |
| AT2G20970 | na | na | 2,46E-05 | -6,55836265 |
| AT5G13210 | NA | Uncharacteris | 2,46E-05 | -4,16142394 |
| AT1G23700 | NA | Protein kinase | 2,46E-05 | -2,39353183 |
| AT5G56430 | NA | F-box/FBD-lik | 2,49E-05 | -8,82287089 |
| AT5G46050 | ATPTR3 | peptide trans | 2,49E-05 | -1,57601978 |
| AT5G10300 | ATMES5 | methyl estera | 2,50E-05 | -1,39089391 |
| AT5G03000 | NA | Galactose oxi | 2,51E-05 | -5,22322672 |
| AT4G22080 | RHS14 | root hair spec | 2,51E-05 | -4,12657362 |
| AT3G22860 | ATEIF3C-2 | eukaryotic tra | 2,51E-05 | -3,81372448 |
| AT1G63210 | NA | Transcription | 2,54E-05 | -6,62573162 |
| AT5G59490 | NA | Haloacid deha | 2,54E-05 | -2,47779956 |
| AT5G43140 | NA | Peroxisomal n | 2,54E-05 | -1,61587534 |
| AT1G44770 | na | na | 2,54E-05 | -1,15491569 |
| AT1G54230 | NA | Winged helix- | 2,61E-05 | -3,05352964 |
| AT2G44790 | UCC2 | uclacyanin 2 | 2,62E-05 | -2,11221489 |

| | | | | |
|---|---|---|---|---|
| AT5G09610 | APUM21 | pumilio 21 | 2,66E-05 | -3,14677703 |
| AT3G50580 | na | na | 2,68E-05 | -3,82238028 |
| AT5G17870 | PSRP6 | plastid-specifi | 2,68E-05 | -1,24284123 |
| AT4G34900 | ATXDH2 | xanthine dehy | 2,68E-05 | -1,21947962 |
| AT3G16300 | NA | Uncharacteris | 2,72E-05 | -4,91829098 |
| AT5G15800 | AGL2 | K-box region a | 2,72E-05 | -4,14510314 |
| AT3G54160 | NA | RNI-like super | 2,74E-05 | -1,93598515 |
| AT3G41979 | na | na | 2,76E-05 | -4,72421886 |
| AT5G23530 | AtCXE18 | carboxyestera | 2,76E-05 | -4,18345999 |
| AT2G28120 | NA | Major facilitat | 2,76E-05 | -2,6122564 |
| AT3G02030 | NA | transferases t | 2,76E-05 | -2,55312695 |
| AT1G04980 | ATPDI10 | PDI-like 2-2 | 2,76E-05 | -1,41333976 |
| AT3G58110 | na | na | 2,77E-05 | 1,03180538 |
| AT2G22140 | ATEME1B | essential meio | 2,77E-05 | -2,86002665 |
| AT1G05020 | NA | ENTH/ANTH/V | 2,77E-05 | -2,6622902 |
| AT5G53680 | NA | RNA-binding ( | 2,78E-05 | -3,73465666 |
| AT3G42960 | ASD | TAPETUM 1 | 2,79E-05 | -3,65163172 |
| AT2G35930 | PUB23 | plant U-box 23 | 2,79E-05 | -3,12939008 |
| AT4G07825 | na | na | 2,79E-05 | -1,07007597 |
| AT4G23600 | CORI3 | Tyrosine trans | 2,86E-05 | 1,56244723 |
| AT4G21590 | ENDO3 | endonuclease | 2,88E-05 | -1,41289955 |
| AT2G46455 | NA | OxaA/YidC-like | 2,90E-05 | -1,83477049 |
| AT5G16190 | ATCSLA11 | cellulose synth | 2,91E-05 | -1,45196215 |
| AT1G54050 | NA | HSP20-like cha | 2,95E-05 | -1,85727728 |
| AT1G75530 | NA | Forkhead-asso | 2,97E-05 | -4,73422162 |
| AT1G35320 | na | na | 2,98E-05 | -11,5803759 |
| AT5G26800 | na | na | 2,98E-05 | -1,24657087 |
| AT1G02400 | ATGA2OX4 | gibberellin 2-o | 3,02E-05 | -5,06173763 |
| AT5G60740 | NA | ABC transport | 3,02E-05 | -3,76877567 |
| AT5G26000 | BGLU38 | thioglucoside | 3,08E-05 | 1,45261634 |
| AT2G30650 | NA | ATP-depender | 3,08E-05 | -4,16084158 |
| AT5G11080 | NA | Ubiquitin-like | 3,12E-05 | -5,35356744 |
| AT1G50580 | NA | UDP-Glycosylt | 3,16E-05 | -4,74831264 |

| | | | | |
|---|---|---|---|---|
| AT1G62630 | NA | Disease resista | 3,16E-05 | -4,36070383 |
| AT3G56530 | anac064 | NAC domain c | 3,16E-05 | -3,93573611 |
| AT5G35600 | HDA7 | histone deace | 3,16E-05 | -3,08488608 |
| AT2G34270 | na | na | 3,21E-05 | -3,55039616 |
| AT2G27630 | NA | Ubiquitin carb | 3,21E-05 | -2,33757195 |
| AT1G63360 | NA | Disease resista | 3,21E-05 | -2,23188746 |
| AT4G12460 | ORP2B | OSBP(oxyster | 3,21E-05 | -1,02949056 |
| AT5G26220 | NA | ChaC-like fam | 3,24E-05 | -2,90572381 |
| AT1G61630 | ATENT7 | equilibrative r | 3,26E-05 | -5,83884446 |
| AT1G60300 | NA | NAC (No Apica | 3,26E-05 | -3,44631413 |
| AT1G35612.1 | na | na | 3,27E-05 | -8,37303842 |
| AT2G18190 | NA | P-loop contair | 3,27E-05 | -4,10426528 |
| AT1G63330 | NA | Pentatricopep | 3,32E-05 | 6,2641192 |
| AT1G62710 | BETA-VPE | beta vacuolar | 3,38E-05 | -2,61620972 |
| AT2G45120 | NA | C2H2-like zinc | 3,38E-05 | -2,15592308 |
| AT4G39580 | NA | Galactose oxid | 3,39E-05 | -2,77304445 |
| AT3G54130 | NA | Josephin fami | 3,41E-05 | -1,37304471 |
| AT1G66460 | NA | Protein kinase | 3,43E-05 | -4,3511224 |
| AT5G18330 | NA | ARM repeat su | 3,43E-05 | -3,63275779 |
| AT1G49160 | WNK7 | Protein kinase | 3,43E-05 | -3,30642154 |
| AT4G34210 | ASK11 | SKP1-like 11 | 3,43E-05 | -3,25083431 |
| AT3G12850 | NA | COP9 signalos | 3,43E-05 | -3,0050214 |
| AT4G14490 | NA | SMAD/FHA dc | 3,43E-05 | -1,94751667 |
| AT5G61120 | na | na | 3,43E-05 | -1,80252728 |
| AT1G65250 | NA | Protein kinase | 3,44E-05 | -3,82868329 |
| AT1G24220 | NA | paired amphip | 3,46E-05 | -4,10156742 |
| AT2G01520 | MLP328 | MLP-like prote | 3,48E-05 | 1,11644199 |
| AT2G39350 | NA | ABC-2 type tra | 3,48E-05 | -2,36354226 |
| AT1G22000 | NA | FBD F-box and | 3,52E-05 | -3,60725467 |
| AT1G21460 | NA | Nodulin MtN3 | 3,53E-05 | -1,60832956 |
| AT5G03480 | NA | RNA-binding ( | 3,56E-05 | -5,44338547 |
| AT1G49360 | NA | F-box family p | 3,56E-05 | -1,37979808 |
| AT1G71160 | KCS7 | 3-ketoacyl-Co | 3,58E-05 | -3,92911799 |

| | | | | |
|---|---|---|---|---|
| AT4G40020 | NA | Myosin heavy | 3,58E-05 | -2,77597615 |
| AT5G37460 | NA | Family of unkı | 3,62E-05 | -3,06590221 |
| AT1G12080 | NA | Vacuolar calci | 3,62E-05 | -1,30545946 |
| AT4G13345 | MEE55 | Serinc-domair | 3,64E-05 | -2,27416555 |
| AT1G19030.1 | na | na | 3,65E-05 | -9,06105785 |
| AT2G25700 | ASK3 | SKP1-like 3 | 3,65E-05 | -8,91594709 |
| AT1G67920 | na | na | 3,65E-05 | -3,16586928 |
| AT5G17190 | na | na | 3,65E-05 | -1,65447345 |
| AT1G74150 | NA | Galactose oxic | 3,65E-05 | -1,62278261 |
| AT4G14040 | EDA38 | selenium-binc | 3,69E-05 | 1,09550099 |
| AT5G02350 | NA | Cysteine/Histi | 3,69E-05 | -6,12818653 |
| AT4G16460 | na | na | 3,69E-05 | -4,55189981 |
| AT2G32310 | NA | CCT motif fam | 3,69E-05 | -3,53614154 |
| AT5G49340 | TBL4 | TRICHOME BII | 3,69E-05 | -3,08125216 |
| AT4G15920 | NA | Nodulin MtN3 | 3,69E-05 | -1,1936573 |
| AT1G54930 | NA | GRF zinc finge | 3,70E-05 | -3,38496328 |
| AT2G28200 | NA | C2H2-type zin | 3,70E-05 | -2,46394798 |
| AT5G38820 | NA | Transmembra | 3,72E-05 | -2,35509313 |
| AT4G04480 | na | na | 3,75E-05 | -3,20501106 |
| AT5G05590 | PAI2 | phosphoribos | 3,77E-05 | 1,34436779 |
| AT3G20710 | NA | F-box family p | 3,77E-05 | -8,98880114 |
| AT1G62590 | NA | pentatricopep | 3,77E-05 | -3,0817739 |
| AT2G20825 | ULT2 | Development | 3,78E-05 | -4,4656851 |
| AT1G30780 | NA | F-box associat | 3,81E-05 | -6,03228653 |
| AT1G34180 | anac016 | NAC domain c | 3,84E-05 | -3,02698121 |
| AT1G66400 | CML23 | calmodulin lik | 3,87E-05 | -3,22984203 |
| AT5G07010 | ATST2A | sulfotransfera | 3,88E-05 | -3,63830112 |
| AT5G22200 | NA | Late embryog | 3,89E-05 | -3,9751732 |
| AT5G13790 | AGL15 | AGAMOUS-lik | 3,89E-05 | -2,17545814 |
| AT5G08730 | ARI16 | IBR domain-cc | 3,92E-05 | -2,67769309 |
| AT5G62960 | na | na | 3,92E-05 | -1,05653376 |
| AT5G48790 | NA | Domain of unl | 3,92E-05 | -1,04661721 |
| AT5G47740 | NA | Adenine nucle | 3,94E-05 | -4,77417151 |

| | | | | |
|---|---|---|---|---|
| AT3G21000 | NA | Gag-Pol-relate | 3,97E-05 | -2,80673095 |
| AT1G61065 | NA | Protein of unk | 3,97E-05 | -1,68037607 |
| AT1G26515 | NA | F-box and ass | 4,00E-05 | -8,98581592 |
| AT3G20520 | SVL3 | SHV3-like 3 | 4,00E-05 | -3,10786896 |
| AT1G24600 | na | na | 4,00E-05 | -2,42943838 |
| AT5G10140 | AGL25 | K-box region a | 4,02E-05 | -4,36523806 |
| AT1G64950 | CYP89A5 | cytochrome P | 4,02E-05 | -1,49897823 |
| AT2G26740 | ATSEH | soluble epoxic | 4,04E-05 | -2,57224952 |
| AT1G60340 | NA | NAC (No Apica | 4,05E-05 | -2,99321757 |
| AT5G51060 | ATRBOHC | NADPH/respir | 4,06E-05 | -3,52399864 |
| AT4G04410.1 | na | na | 4,07E-05 | 2,98095919 |
| AT4G01360 | na | na | 4,07E-05 | -2,46949946 |
| AT2G16450 | NA | F-box and ass | 4,10E-05 | -6,19944545 |
| AT4G09450 | NA | Duplicated ho | 4,10E-05 | -4,24735579 |
| AT5G65790 | ATMYB68 | myb domain p | 4,13E-05 | -2,23363127 |
| AT2G34100 | na | na | 4,15E-05 | -5,71836867 |
| AT3G03650 | EDA5 | Exostosin fam | 4,15E-05 | -3,32579946 |
| AT5G03350 | NA | Legume lectin | 4,15E-05 | -2,6863475 |
| AT3G28570 | NA | P-loop contair | 4,16E-05 | -2,22444332 |
| AT1G15310 | ATHSRP54A | signal recogni | 4,16E-05 | -1,31803956 |
| AT5G50060 | NA | Plant invertas | 4,20E-05 | -4,86058282 |
| AT1G05920 | NA | Domain of unl | 4,20E-05 | -4,06383397 |
| AT4G37780 | ATMYB87 | myb domain p | 4,20E-05 | -2,76622209 |
| AT5G25400 | NA | Nucleotide-su | 4,20E-05 | -2,32547695 |
| AT2G38530 | cdf3 | lipid transfer p | 4,20E-05 | -1,83309953 |
| AT5G52340 | ATEXO70A2 | exocyst subur | 4,24E-05 | -1,6622331 |
| AT5G42730 | na | na | 4,25E-05 | -10,8541681 |
| AT4G37520 | NA | Peroxidase su | 4,25E-05 | -4,12163536 |
| AT5G18340 | NA | ARM repeat su | 4,28E-05 | -3,27561296 |
| AT3G55780 | NA | Glycosyl hydro | 4,28E-05 | -1,91151151 |
| AT1G69550 | NA | disease resista | 4,33E-05 | -2,0179357 |
| AT3G19070 | NA | Homeodomai | 4,35E-05 | -5,29547574 |
| AT1G02050 | LAP6 | Chalcone and | 4,36E-05 | -3,80200996 |

| | | | | |
|---|---|---|---|---|
| AT5G36930 | NA | Disease resista | 4,42E-05 | -11,3124755 |
| AT4G12920 | NA | Eukaryotic asp | 4,42E-05 | -5,45843622 |
| AT1G11510 | NA | DNA-binding s | 4,42E-05 | -4,01447043 |
| AT5G42280 | NA | Cysteine/Histi | 4,42E-05 | -1,33653098 |
| AT4G31330 | NA | Protein of unk | 4,50E-05 | -2,95221595 |
| AT5G39790 | NA | 5prime-AMP-a | 4,50E-05 | -1,27820174 |
| AT5G59230 | NA | transcription f | 4,56E-05 | -2,72593658 |
| AT2G35070 | na | na | 4,63E-05 | -5,39679604 |
| AT5G40940 | FLA20 | putative fascic | 4,63E-05 | -2,57065779 |
| AT1G61610 | NA | S-locus lectin p | 4,70E-05 | 3,49424043 |
| AT2G40130 | NA | Double Clp-N | 4,71E-05 | 1,08067121 |
| AT2G45430 | AHL22 | AT-hook motif | 4,71E-05 | -4,20562011 |
| AT4G00610 | NA | DNA-binding s | 4,72E-05 | -3,95646582 |
| AT5G37980 | NA | Zinc-binding d | 4,75E-05 | 2,87617218 |
| AT3G13640 | ATRLI1 | RNAse l inhibit | 4,79E-05 | -1,30828349 |
| AT5G41800 | NA | Transmembra | 4,87E-05 | -2,63518058 |
| AT4G34881 | na | na | 4,89E-05 | -10,2796157 |
| AT3G50330 | HEC2 | basic helix-loc | 4,93E-05 | -4,40689186 |
| AT3G13220 | ABCG26 | ABC-2 type tra | 5,04E-05 | -4,01651019 |
| AT1G35440 | CYCT1;1 | cyclin T1;1 | 5,05E-05 | -6,00000487 |
| AT3G07970 | QRT2 | Pectin lyase-li | 5,05E-05 | -3,87790566 |
| AT1G66990 | na | na | 5,09E-05 | 4,83868662 |
| AT3G10116 | NA | COBRA-like ex | 5,11E-05 | -10,2121616 |
| AT5G15620 | NA | F-box/RNI-like | 5,12E-05 | -3,68069833 |
| AT1G47520.1 | na | na | 5,14E-05 | -11,206851 |
| AT5G60142 | NA | AP2/B3-like tr | 5,15E-05 | -2,45134781 |
| AT5G37920 | NA | Family of unkn | 5,19E-05 | -5,16397794 |
| AT3G62570 | NA | Tetratricopept | 5,29E-05 | -1,31385128 |
| AT4G02700 | SULTR3;2 | sulfate transp | 5,31E-05 | -2,12443177 |
| AT5G55896.1 | na | na | 5,40E-05 | -3,06730862 |
| AT4G35070 | NA | SBP (S-ribonuc | 5,43E-05 | -1,36951313 |
| AT1G79450 | ALIS5 | ALA-interactin | 5,49E-05 | -3,0642631 |
| AT3G21230 | 4CL5 | 4-coumarate:( | 5,49E-05 | -1,80340233 |

| | | | | |
|---|---|---|---|---|
| AT4G04690 | NA | F-box and ass | 5,56E-05 | -2,74618317 |
| AT5G37470 | NA | Family of unk | 5,57E-05 | -2,60816988 |
| AT1G05490 | chr31 | chromatin rer | 5,57E-05 | -1,16074095 |
| AT2G39380 | ATEXO70H2 | exocyst subur | 5,71E-05 | -10,3602238 |
| AT1G43590.1 | na | na | 5,77E-05 | -8,05280691 |
| AT4G22600 | na | na | 5,88E-05 | -3,5829781 |
| AT3G61340 | NA | F-box and ass | 5,88E-05 | -3,46114029 |
| AT5G15060 | NA | Lateral organ | 5,88E-05 | -3,06228143 |
| AT3G58260 | NA | TRAF-like fam | 5,90E-05 | -6,26163647 |
| AT4G38280 | na | na | 5,92E-05 | -2,36545077 |
| AT5G25950 | NA | Protein of Unl | 6,01E-05 | -1,74822668 |
| AT1G60270 | BGLU6 | beta glucosida | 6,02E-05 | -3,06363798 |
| AT3G04620 | NA | Alba DNA/RNA | 6,02E-05 | -1,54927478 |
| AT2G40010 | NA | Ribosomal prc | 6,26E-05 | -11,1481013 |
| AT5G43070 | WPP1 | WPP domain p | 6,26E-05 | -1,42961748 |
| AT4G30030 | NA | Eukaryotic asp | 6,27E-05 | -3,75222753 |
| AT5G56747.1 | na | na | 6,34E-05 | -6,31346529 |
| AT1G45248 | NA | Nucleolar hist | 6,34E-05 | -5,75529269 |
| AT3G61810 | NA | Glycosyl hydrc | 6,38E-05 | -1,4798649 |
| AT1G23770 | NA | F-box family p | 6,48E-05 | -3,03085531 |
| AT4G39760 | NA | Galactose oxic | 6,49E-05 | -8,44092577 |
| AT5G25230 | NA | Ribosomal prc | 6,49E-05 | -2,50578976 |
| AT4G17200 | NA | F-box and ass | 6,60E-05 | -3,55716013 |
| AT3G09405 | NA | Pectinacetyles | 6,61E-05 | -2,59413284 |
| AT1G01670 | NA | RING/U-box su | 6,65E-05 | 1,39615575 |
| AT5G49240 | APRR4 | pseudo-respo | 6,65E-05 | -6,60126914 |
| AT2G40180 | ATHPP2C5 | phosphatase 2 | 6,81E-05 | -2,23424544 |
| AT5G38490 | NA | Domain of unl | 6,91E-05 | -5,89556476 |
| AT5G14300 | ATPHB5 | prohibitin 5 | 6,93E-05 | -4,6943504 |
| AT1G78720 | NA | SecY protein t | 7,01E-05 | -3,97187961 |
| AT2G26400 | ARD | acireductone | 7,06E-05 | 1,12005513 |
| AT4G01023 | NA | RING/U-box su | 7,08E-05 | -3,10103834 |
| AT4G23990 | ATCSLG3 | cellulose syntl | 7,19E-05 | 1,02652114 |

| | | | | |
|---|---|---|---|---|
| AT5G54064 | na | na | 7,19E-05 | -5,11008284 |
| AT3G24230 | NA | Pectate lyase | 7,19E-05 | -2,94413889 |
| AT4G05020 | NDB2 | NAD(P)H dehy | 7,19E-05 | -1,1913598 |
| AT5G52890 | NA | AT hook motif | 7,19E-05 | -1,17256497 |
| AT4G01500 | NGA4 | AP2/B3-like tr | 7,20E-05 | -2,52292228 |
| AT5G01180 | ATPTR5 | peptide trans | 7,25E-05 | -2,743952 |
| AT2G42960 | NA | Protein kinase | 7,26E-05 | 1,04594563 |
| AT1G03620 | NA | ELMO/CED-12 | 7,26E-05 | -5,2177183 |
| AT4G09960 | AGL11 | K-box region a | 7,37E-05 | -7,46298895 |
| AT5G51830 | NA | pfkB-like carb | 7,44E-05 | -1,08512019 |
| AT1G50680 | NA | AP2/B3 transc | 7,51E-05 | -3,67729135 |
| AT2G34870 | MEE26 | hydroxyprolin | 7,54E-05 | -2,1669109 |
| AT4G05310 | NA | Ubiquitin-like | 7,55E-05 | -3,35251767 |
| AT1G74220 | na | na | 7,55E-05 | -2,48713536 |
| AT5G54330 | NA | Protein of unk | 7,60E-05 | -5,90175003 |
| AT5G14180 | MPL1 | Myzus persica | 7,63E-05 | -2,362592 |
| AT1G70020 | NA | Protein of unk | 7,69E-05 | -5,88964813 |
| AT1G21320 | NA | nucleotide bir | 7,77E-05 | -2,43949935 |
| AT3G59160 | NA | F-box/RNI-like | 7,84E-05 | -3,40618751 |
| AT1G80160 | NA | Lactoylglutath | 7,84E-05 | -2,73200921 |
| AT5G07230 | NA | Bifunctional ir | 7,87E-05 | -4,16150828 |
| AT1G62200 | NA | Major facilitat | 7,96E-05 | 1,07923328 |
| AT3G08860 | PYD4 | PYRIMIDINE 4 | 7,96E-05 | -3,24651627 |
| AT5G39940 | NA | FAD/NAD(P)-b | 8,02E-05 | -1,05906416 |
| AT5G35796 | na | na | 8,03E-05 | 8,58614564 |
| AT4G35380 | NA | SEC7-like guar | 8,05E-05 | -4,61886611 |
| AT1G58340 | ZF14 | MATE efflux fa | 8,34E-05 | 1,85229328 |
| AT4G35900 | atbzip14 | Basic-leucine z | 8,35E-05 | 1,16267436 |
| AT1G70220 | NA | RNA-processir | 8,37E-05 | -2,13786933 |
| AT4G23700 | ATCHX17 | cation/H+ exc | 8,41E-05 | -2,81294363 |
| AT2G25600 | AKT6 | Shaker pollen | 8,41E-05 | -2,67022274 |
| AT2G32470 | NA | F-box associat | 8,48E-05 | -6,21131687 |
| AT1G11920 | NA | Pectin lyase-li | 8,54E-05 | -3,79204892 |

| | | | | |
|---|---|---|---|---|
| AT1G44740 | na | na | 8,54E-05 | -2,92349326 |
| AT2G05960.1 | na | na | 8,56E-05 | -4,53312019 |
| AT4G12580 | na | na | 8,59E-05 | -3,69004894 |
| AT3G09160 | NA | RNA-binding ( | 8,67E-05 | -4,08431822 |
| AT5G36240 | NA | zinc knuckle (( | 8,73E-05 | 3,02307965 |
| AT4G28130 | ATDGK6 | diacylglycerol | 8,73E-05 | -3,77338243 |
| AT3G45560 | NA | zinc finger (C3 | 8,74E-05 | -6,33542906 |
| AT1G75030 | ATLP-3 | thaumatin-like | 8,79E-05 | -3,25690528 |
| AT3G51190 | NA | Ribosomal pro | 8,87E-05 | -5,64615176 |
| AT4G21010 | NA | Transcription | 8,94E-05 | -4,09081649 |
| AT3G55730 | AtMYB109 | myb domain p | 8,96E-05 | -1,41389977 |
| AT5G56400 | NA | FBD F-box Skp | 9,04E-05 | -4,34695265 |
| AT2G27780 | NA | Transcription | 9,16E-05 | -3,63675523 |
| AT4G16880 | NA | Leucine-rich r | 9,19E-05 | -7,62598011 |
| AT1G69180 | CRC | Plant-specific | 9,25E-05 | -2,00139835 |
| AT3G62430 | NA | Protein with R | 9,31E-05 | -1,66289722 |
| AT4G20460 | NA | NAD(P)-bindir | 9,39E-05 | -2,19580083 |
| AT1G51538 | NA | Aminotransfer | 9,43E-05 | 1,32123152 |
| AT2G42830 | AGL5 | K-box region a | 9,48E-05 | -4,05496959 |
| AT4G36850 | NA | PQ-loop repea | 9,49E-05 | -1,46540151 |
| AT1G25550 | NA | myb-like trans | 9,50E-05 | -1,98287167 |
| AT1G74460 | NA | GDSL-like Lipa | 9,50E-05 | -1,79934044 |
| AT5G41910 | MED10A | Mediator com | 9,54E-05 | -1,66437309 |
| AT4G08800 | NA | Protein kinase | 9,57E-05 | -8,14959442 |
| AT1G52618 | na | na | 9,59E-05 | -10,9106069 |
| AT5G43470 | HRT | Disease resista | 9,59E-05 | -2,41832253 |
| AT4G13885 | NA | Polynucleotidy | 9,61E-05 | 1,7927654 |
| AT5G28080 | WNK9 | Protein kinase | 9,61E-05 | -4,83930123 |
| AT3G26440 | NA | Protein of unk | 9,62E-05 | -2,63322539 |
| AT2G21100 | NA | Disease resista | 9,62E-05 | -2,62619154 |
| AT1G51420 | ATSPP1 | sucrose-phosp | 9,62E-05 | -2,25636059 |
| AT3G21010.1 | na | na | 9,62E-05 | -1,36280831 |
| AT1G03490 | ANAC006 | NAC domain c | 9,63E-05 | -5,48618618 |

| | | | | |
|---|---|---|---|---|
| AT4G22120 | NA | ERD (early-res | 9,68E-05 | -1,07224516 |
| AT4G36830 | HOS3-1 | GNS1/SUR4 m | 9,73E-05 | 1,94071296 |
| AT3G46070 | NA | C2H2-type zin | 9,73E-05 | -8,95934568 |
| AT1G01280 | CYP703 | cytochrome P | 9,73E-05 | -4,30900739 |
| AT2G18260 | ATSYP112 | syntaxin of pla | 9,73E-05 | -2,97912076 |
| AT1G48700 | NA | 2-oxoglutarate | 9,75E-05 | 5,4221802 |
| AT4G33930 | NA | Cupredoxin su | 9,75E-05 | -9,02421186 |
| AT2G26940 | NA | C2H2-type zin | 9,78E-05 | -4,72842245 |
| AT3G28540 | NA | P-loop contair | 9,81E-05 | -2,08957723 |
| AT4G39010 | AtGH9B18 | glycosyl hydrc | 9,82E-05 | -1,45287746 |
| AT3G21120 | NA | F-box and asso | 9,90E-05 | -8,5798787 |
| AT5G27940 | WPP3 | WPP domain p | 9,91E-05 | -3,10431248 |
| AT4G19460 | NA | UDP-Glycosylt | 9,96E-05 | -3,87599695 |
| AT3G21030.1 | na | na | 9,99E-05 | -2,65929293 |
| AT5G17200 | NA | Pectin lyase-li | 0,00010039 | -5,09020164 |
| AT5G16010 | NA | 3-oxo-5-alpha | 0,00010097 | 1,25186457 |
| AT1G53887 | na | na | 0,00010267 | -3,73028133 |
| AT5G39610 | ANAC092 | NAC domain c | 0,00010267 | -2,80518382 |
| AT1G77850 | ARF17 | auxin respons | 0,00010267 | -1,08621449 |
| AT1G66170 | MMD1 | RING/FYVE/PH | 0,0001029 | -2,5308146 |
| AT5G35777.1 | na | na | 0,00010295 | -4,49083263 |
| AT5G05440 | PYL5 | Polyketide cyc | 0,00010321 | -2,29149511 |
| AT3G17010 | NA | AP2/B3-like tr | 0,00010339 | -1,66553552 |
| AT3G53480 | ABCG37 | pleiotropic dru | 0,00010458 | -2,5068087 |
| AT2G28140 | NA | Protein of unk | 0,00010465 | -2,4255916 |
| AT5G61850 | LFY | floral meristei | 0,0001049 | 1,29164934 |
| AT1G06250 | NA | alpha/beta-Hy | 0,0001049 | -1,78194954 |
| AT3G44770 | NA | Protein of unk | 0,00010511 | -1,28853502 |
| AT5G38000 | NA | Zinc-binding d | 0,0001053 | 3,01781935 |
| AT5G01610 | NA | Protein of unk | 0,00010597 | -2,1038813 |
| AT3G21620 | NA | ERD (early-res | 0,00010606 | -2,94534635 |
| AT1G02470 | NA | Polyketide cyc | 0,00010643 | -3,56010213 |
| AT4G19560 | CYCT1;2 | Cyclin family p | 0,00010792 | -2,21619685 |

| | | | | |
|---|---|---|---|---|
| AT1G76830 | NA | F-box and ass | 0,00010799 | -3,19440889 |
| AT1G21540 | NA | AMP-depende | 0,0001081 | -1,09567291 |
| AT2G17950 | PGA6 | Homeodomai | 0,00010844 | -2,26430693 |
| AT1G75750 | GASA1 | GAST1 protei | 0,00010931 | -1,65496193 |
| AT5G17880 | CSA1 | disease resist | 0,00010993 | -10,9121652 |
| AT3G23870 | NA | Protein of unk | 0,00010993 | -2,572021 |
| AT5G49200 | NA | WD-40 repeat | 0,00011009 | -4,01748967 |
| AT4G03320 | tic20-IV | translocon at | 0,00011013 | -1,65068512 |
| AT2G30080 | ATZIP6 | ZIP metal ion | 0,00011026 | -1,85551928 |
| AT1G78000 | SEL1 | sulfate transp | 0,00011073 | -3,45977573 |
| AT1G69600 | ATHB29 | zinc finger hor | 0,00011073 | -1,06984247 |
| AT1G16510 | NA | SAUR-like aux | 0,00011116 | -2,89441633 |
| AT4G18680 | na | na | 0,00011131 | -2,40653592 |
| AT1G53020 | na | na | 0,0001114 | -4,2510853 |
| AT5G51630 | NA | Disease resist | 0,00011231 | -10,7513751 |
| AT5G56570 | NA | Leucine-rich r | 0,00011438 | -6,17455625 |
| AT2G31760 | ARI10 | RING/U-box su | 0,00011438 | -5,03941774 |
| AT1G53660 | NA | nodulin MtN2 | 0,00011463 | 2,68595153 |
| AT2G45560 | CYP76C1 | cytochrome P | 0,00011477 | 1,85767356 |
| AT5G45150 | RTL3 | RNAse THREE- | 0,00011522 | 1,28639972 |
| AT2G25790 | NA | Leucine-rich r | 0,00011537 | 1,30669007 |
| AT1G14580 | NA | C2H2-like zinc | 0,00011585 | 1,21748143 |
| AT4G11070 | AtWRKY41 | WRKY family t | 0,00011585 | -10,6980681 |
| AT4G09300 | NA | LisH and RanB | 0,00011637 | -10,6764525 |
| AT1G66500 | NA | Pre-mRNA cle | 0,00011666 | -2,77645848 |
| AT1G50820 | NA | Aminotransfe | 0,00011808 | -2,30408962 |
| AT4G21850 | ATMSRB9 | methionine su | 0,00011831 | 2,29137185 |
| AT1G22010 | na | na | 0,00011871 | -4,84700469 |
| AT1G61180 | NA | LRR and NB-A | 0,00011902 | -1,62617363 |
| AT3G06100 | NIP7;1 | NOD26-like in | 0,00011917 | -3,58951988 |
| AT1G52430 | NA | Ubiquitin carb | 0,00011952 | -2,08189125 |
| AT5G55590 | QRT1 | Pectin lyase-li | 0,0001216 | -4,04423628 |
| AT5G27750 | NA | F-box/FBD-lik | 0,00012206 | -1,46071322 |

| | | | | |
|---|---|---|---|---|
| AT1G78640 | na | na | 0,00012664 | -3,74425336 |
| AT5G60650 | na | na | 0,00012731 | -1,59541826 |
| AT4G03350 | NA | ubiquitin fami | 0,00012766 | -3,30163395 |
| AT5G43660 | NA | 2-oxoglutarat | 0,00012814 | -2,2081825 |
| AT1G03850 | NA | Glutaredoxin f | 0,00012814 | -2,13778793 |
| AT5G37890 | NA | Protein with R | 0,00012814 | -1,06938516 |
| AT5G18180 | NA | H/ACA ribonu | 0,00012875 | -1,97971029 |
| AT4G19990 | FRS1 | FAR1-related s | 0,00012903 | 1,21285665 |
| AT4G21680 | NRT1.8 | NITRATE TRAN | 0,00012986 | -1,93525929 |
| AT1G70640 | NA | octicosapeptid | 0,00013053 | 2,76683797 |
| AT5G25130 | CYP71B12 | cytochrome P | 0,00013077 | 1,65908661 |
| AT1G78160 | APUM7 | pumilio 7 | 0,00013114 | -4,76048375 |
| AT1G09930 | ATOPT2 | oligopeptide t | 0,00013119 | -2,78734492 |
| AT5G24910 | CYP714A1 | cytochrome P | 0,0001314 | 1,63393658 |
| AT5G03795 | NA | Exostosin fam | 0,00013278 | -3,10988378 |
| AT5G37440 | NA | Chaperone Dr | 0,00013431 | -10,0488193 |
| AT3G11165 | na | na | 0,00013456 | -4,62270126 |
| AT2G27120 | POL2B | DNA polymera | 0,00013605 | -3,25671961 |
| AT5G16090 | na | na | 0,00013707 | -3,85072517 |
| AT1G50970 | NA | Membrane tra | 0,00013726 | -1,5081207 |
| AT1G52770 | NA | Phototropic-re | 0,00013947 | 2,82373324 |
| AT4G16807 | na | na | 0,00013947 | -1,86488602 |
| AT1G58160 | NA | Mannose-binc | 0,00013974 | -9,2709684 |
| AT3G10410 | CPY | SERINE CARBC | 0,00013999 | -1,56782148 |
| AT4G22870 | NA | 2-oxoglutarat | 0,00014026 | -4,86490612 |
| AT3G44760 | na | na | 0,0001411 | -4,26303697 |
| AT5G47150 | NA | YDG/SRA dom | 0,00014228 | -3,76610417 |
| AT5G56370 | NA | F-box/RNI-like | 0,00014355 | -10,827267 |
| AT5G63560 | NA | HXXXD-type a | 0,00014447 | -4,72552859 |
| AT1G30760 | NA | FAD-binding B | 0,00014447 | -1,23788206 |
| AT5G50800 | NA | Nodulin MtN3 | 0,00014569 | -3,59426989 |
| AT3G07250 | NA | nuclear transp | 0,00014686 | -2,69673863 |
| AT5G23660 | MTN3 | homolog of M | 0,00014795 | -2,50562905 |

| | | | | |
|---|---|---|---|---|
| AT3G15950 | NAI2 | DNA topoisom | 0,00014831 | 1,48059861 |
| AT1G18265 | NA | Protein of unk | 0,00014833 | -3,29422149 |
| AT5G41120 | NA | Esterase/lipas | 0,00014833 | -1,05042036 |
| AT1G58590 | na | na | 0,00014969 | -10,5282309 |
| AT5G10170 | ATMIPS3 | myo-inositol-1 | 0,00015038 | -1,1808472 |
| AT2G21140 | ATPRP2 | proline-rich pr | 0,00015051 | -1,72811622 |
| AT3G04660 | NA | F-box and ass | 0,00015104 | -5,4062682 |
| AT5G02140 | NA | Pathogenesis- | 0,00015139 | 2,00854709 |
| AT1G05080 | NA | F-box/RNI-like | 0,00015362 | -4,97699467 |
| AT5G65090 | BST1 | DNAse I-like su | 0,00015396 | -2,29876704 |
| AT3G02000 | ROXY1 | Thioredoxin su | 0,00015439 | 1,47985223 |
| AT3G10525 | LGO | LOSS OF GIAN | 0,00015608 | -1,76440414 |
| AT5G60610 | NA | F-box/RNI-like | 0,00015617 | -3,69661148 |
| AT4G31520 | NA | SDA1 family p | 0,00015617 | -2,21881122 |
| AT4G20420 | NA | Tapetum spec | 0,00015989 | -5,61331213 |
| AT5G51270 | NA | U-box domain | 0,00015999 | -1,79379353 |
| AT4G20350 | NA | oxidoreductas | 0,00016296 | -2,80451137 |
| AT5G09520 | NA | hydroxyprolin | 0,00016602 | -5,06191563 |
| AT1G67481 | na | na | 0,00016697 | -1,83510914 |
| AT2G48130 | NA | Bifunctional ir | 0,00016725 | -4,02226763 |
| AT3G56560 | anac065 | NAC domain c | 0,00016921 | -5,05981723 |
| AT2G29810 | NA | Galactose oxi | 0,00016927 | -4,43819518 |
| AT2G01960 | TET14 | tetraspanin14 | 0,00016952 | -8,16961881 |
| AT5G48620 | NA | Disease resista | 0,00016974 | -1,13560621 |
| AT5G52330 | NA | TRAF-like supe | 0,00017281 | -5,3712346 |
| AT2G24000 | scpl22 | serine carboxy | 0,00017339 | -1,78625588 |
| AT5G43403 | na | na | 0,00017437 | -10,4129486 |
| AT4G10220 | NA | Protein of Unk | 0,00017449 | 8,0701453 |
| AT1G63340 | NA | Flavin-contain | 0,00017537 | 5,02600268 |
| AT4G19030 | AT-NLM1 | NOD26-like m | 0,00017537 | 3,22601669 |
| AT1G70820 | NA | phosphogluco | 0,00017537 | 1,01859015 |
| AT1G43140 | NA | Cullin family p | 0,00017537 | -6,05305702 |
| AT3G33235.1 | na | na | 0,00017537 | -3,52181683 |

| | | | | |
|---|---|---|---|---|
| AT5G07060 | NA | CCCH-type zin | 0,00017537 | -2,48987497 |
| AT5G25470 | NA | AP2/B3-like tr | 0,00017537 | -2,35853061 |
| AT4G19020 | CMT2 | chromomethy | 0,00017715 | 1,24092422 |
| AT5G53660 | AtGRF7 | growth-regula | 0,00017715 | 1,03874697 |
| AT4G30040 | NA | Eukaryotic asp | 0,00017715 | -4,59309091 |
| AT1G35290 | NA | Thioesterase s | 0,00017715 | -1,83147185 |
| AT1G61320 | NA | FBD / Leucine | 0,00017739 | 8,52603083 |
| AT1G80820 | ATCCR2 | cinnamoyl coa | 0,00017739 | -1,50850557 |
| AT1G61450 | na | na | 0,00017771 | -1,70678296 |
| AT1G20620 | ATCAT3 | catalase 3 | 0,00017906 | 1,18865948 |
| AT1G02640 | ATBXL2 | beta-xylosidas | 0,00017943 | -1,18474063 |
| AT2G26290 | ARSK1 | root-specific k | 0,00018171 | -3,03742476 |
| AT3G49270 | na | na | 0,00018289 | -4,08214888 |
| AT2G29820 | NA | Galactose oxic | 0,00018442 | -3,30992182 |
| AT5G52900 | na | na | 0,00018527 | 1,75798883 |
| AT2G01810 | NA | RING/FYVE/PH | 0,00018527 | -2,5925724 |
| AT5G38740 | AGL77 | AGAMOUS-lik | 0,0001859 | -10,3656697 |
| AT3G03930 | NA | protein kinase | 0,00018688 | -8,15822462 |
| AT5G49260 | na | na | 0,00018721 | -6,45453503 |
| AT5G13170 | SAG29 | senescence-as | 0,0001876 | -2,2692653 |
| AT1G14760 | KNATM | KNOX Arabido | 0,00018763 | -2,87740236 |
| AT5G41820 | ATRGTA2 | RAB geranylge | 0,00018789 | -5,81214321 |
| AT4G12000 | NA | SNARE associa | 0,00018789 | -1,04797191 |
| AT1G60730 | NA | NAD(P)-linked | 0,00018796 | -2,78971285 |
| AT5G04770 | ATCAT6 | cationic amino | 0,00018796 | -1,6407663 |
| AT2G02061 | NA | Nucleotide-di | 0,00018853 | -1,45471533 |
| AT1G77410 | BGAL16 | beta-galactosi | 0,00019039 | -1,46851618 |
| AT5G07475 | NA | Cupredoxin su | 0,00019234 | -1,88709388 |
| AT3G29380 | NA | Cyclin-like fam | 0,00019551 | -3,44853362 |
| AT3G44757 | na | na | 0,00019591 | -8,00704608 |
| AT1G63860 | NA | Disease resista | 0,00019807 | -10,5813318 |
| AT4G33020 | ATZIP9 | ZIP metal ion | 0,00019807 | -2,04507634 |
| AT4G34890 | ATXDH1 | xanthine dehy | 0,00019904 | 1,18733932 |

| | | | | |
|---|---|---|---|---|
| AT5G25640 | NA | Rhomboid-rel | 0,00019916 | 3,80506683 |
| AT4G32510 | NA | HCO3- transpo | 0,00019916 | -2,88307756 |
| AT2G23290 | AtMYB70 | myb domain p | 0,00019916 | -1,09726218 |
| AT2G27750 | NA | Surfeit locus p | 0,00020037 | -8,34704489 |
| AT1G52827 | ATCDT1 | cadmium tole | 0,00020037 | -2,64438513 |
| AT3G24840 | NA | Sec14p-like ph | 0,00020037 | -1,01209689 |
| AT4G16030 | NA | Ribosomal pro | 0,00020165 | -8,51459842 |
| AT2G27650 | NA | Ubiquitin carb | 0,00020246 | -2,5077585 |
| AT2G29830 | NA | Galactose oxid | 0,00020372 | -3,05721925 |
| AT3G15570 | NA | Phototropic-re | 0,00020373 | 1,15559512 |
| AT4G23020 | na | na | 0,00020498 | 1,29963958 |
| AT1G63206 | NA | Cystatin/mono | 0,00020498 | -3,57528586 |
| AT4G18900 | NA | Transducin/W | 0,00020589 | -5,78786369 |
| AT5G13130 | NA | Histidine kinas | 0,00020637 | -4,62213011 |
| AT1G02610 | NA | RING/FYVE/PH | 0,00020676 | -4,83572577 |
| AT4G26370 | NA | antiterminatio | 0,0002074 | 1,00802414 |
| AT1G72980 | LBD7 | LOB domain-c | 0,00020775 | -5,57548345 |
| AT5G44440 | NA | FAD-binding B | 0,00020875 | -4,67083576 |
| AT2G16190 | na | na | 0,00020907 | -9,35437198 |
| AT5G35450 | NA | Disease resista | 0,00021217 | -3,35505439 |
| AT5G42700 | NA | AP2/B3-like tr | 0,00021217 | -1,60065012 |
| AT5G22160 | na | na | 0,0002177 | -9,39829022 |
| AT3G15700 | NA | P-loop contair | 0,0002177 | -4,98867035 |
| AT4G14780 | NA | Protein kinase | 0,0002177 | -4,62548859 |
| AT2G21730 | ATCAD2 | cinnamyl alcol | 0,0002177 | -4,13445913 |
| AT4G24350 | NA | Phosphorylase | 0,00022032 | 1,37300064 |
| AT1G07750 | NA | RmlC-like cupi | 0,00022123 | -1,03880424 |
| AT3G52957 | na | na | 0,00022148 | -8,15198505 |
| AT1G61490 | NA | S-locus lectin | 0,00022228 | 1,48728682 |
| AT4G12990 | na | na | 0,00022279 | -10,2632985 |
| AT4G03170 | NA | AP2/B3-like tr | 0,00022279 | -3,21322559 |
| AT5G03790 | ATHB51 | homeobox 51 | 0,00022518 | 1,42927598 |
| AT2G02060 | NA | Homeodomai | 0,00022565 | 1,52569089 |

| | | | | | |
|---|---|---|---|---|---|
| AT3G17530 | NA | | F-box and ass | 0,00022665 | -4,31126797 |
| AT4G00250 | NA | | DNA-binding s | 0,00022707 | -4,55042667 |
| AT2G39240 | NA | | RNA polymera | 0,00022767 | -4,80492057 |
| AT1G27280 | NA | | Paired amphip | 0,00022856 | -2,10663683 |
| AT3G16380 | PAB6 | | poly(A) bindin | 0,00023072 | -1,51979977 |
| AT5G22020 | NA | | Calcium-depe | 0,00023079 | -1,23636618 |
| AT5G43740 | NA | | Disease resist | 0,00023343 | -5,05999802 |
| AT2G33880 | HB-3 | | homeobox-3 | 0,00023472 | -2,0226283 |
| AT1G24250 | NA | | Paired amphip | 0,00023506 | -3,38606868 |
| AT3G23770 | NA | | O-Glycosyl hy | 0,00023524 | -3,81844178 |
| AT1G05930 | NA | | Domain of unl | 0,00023765 | -2,55059521 |
| AT5G49420 | NA | | MADS-box tra | 0,00023771 | -4,57079827 |
| AT2G31460 | NA | | Domain of unl | 0,00023793 | -2,85815756 |
| AT3G13610 | NA | | 2-oxoglutarat | 0,00023833 | -1,76439082 |
| AT5G51730 | NA | | RNA-binding ( | 0,00023879 | -1,03937523 |
| AT1G60250 | NA | | B-box zinc fing | 0,00024116 | -8,55994029 |
| AT1G65342 | na | | na | 0,00024126 | -9,22772638 |
| AT4G21720 | na | | na | 0,00024172 | -1,42157745 |
| AT2G32350 | NA | | Ubiquitin-like | 0,00024329 | -1,56588819 |
| AT4G05631 | na | | na | 0,00024366 | -10,1797556 |
| AT4G00160 | NA | | F-box/RNI-like | 0,00024729 | -9,25564043 |
| AT4G09250 | NA | | SPla/RYanodir | 0,00024818 | -10,279744 |
| AT2G29800 | NA | | Galactose oxi | 0,00024836 | -3,44227293 |
| AT3G13900 | NA | | ATPase E1-E2 | 0,00024964 | -1,85504808 |
| AT4G11730 | NA | | Cation transp | 0,00024997 | -5,72558831 |
| AT3G57950 | na | | na | 0,00025059 | -3,36321305 |
| AT1G49070.1 | na | | na | 0,00025068 | 8,48679648 |
| AT4G01593 | na | | na | 0,00025073 | -10,2527745 |
| AT1G73120 | na | | na | 0,00025202 | 4,17276922 |
| AT2G39820 | NA | | Translation ini | 0,00025404 | -2,22952761 |
| AT1G73220 | | 01-oct | organic cation | 0,00025508 | -4,44143559 |
| AT2G27060 | NA | | Leucine-rich r | 0,00025613 | 1,07332414 |
| AT4G38932 | na | | na | 0,00025638 | 1,00620471 |

| | | | | |
|---|---|---|---|---|
| AT5G47635 | NA | Pollen Ole e 1 | 0,0002565 | -4,32827985 |
| AT5G09210 | NA | GC-rich seque | 0,00026175 | -3,86509153 |
| AT4G37310 | CYP81H1 | cytochrome P | 0,00026582 | -1,26318092 |
| AT1G67160 | NA | F-box family p | 0,00026649 | -5,35101352 |
| AT5G28330 | na | na | 0,00026659 | -5,14110971 |
| AT5G38393 | na | na | 0,00026662 | -8,58785747 |
| AT3G45970 | ATEXLA1 | expansin-like A | 0,00026712 | -2,73926408 |
| AT3G08810 | NA | Galactose oxid | 0,00026787 | -2,7232095 |
| AT4G13572 | na | na | 0,00026875 | 5,78693478 |
| AT3G56230 | NA | BTB/POZ dom | 0,00026909 | -3,60497847 |
| AT2G12480 | SCPL43 | serine carboxy | 0,00026977 | -6,36875985 |
| AT3G19610 | NA | Plant protein ( | 0,00027005 | -5,24237701 |
| AT2G14710 | NA | F-box family p | 0,00027057 | -5,36618459 |
| AT4G23310 | CRK23 | cysteine-rich R | 0,00027093 | 9,36963739 |
| AT4G08991 | na | na | 0,00027136 | -10,103996 |
| AT4G10680 | NA | transcription f | 0,00027136 | -4,14179861 |
| AT1G65140 | NA | Ubiquitin carb | 0,00027273 | -2,31503061 |
| AT5G13700 | APAO | polyamine oxi | 0,00027317 | -2,11144101 |
| AT5G24860 | ATFPF1 | flowering prom | 0,00027432 | 1,17755003 |
| AT4G14805 | NA | Bifunctional in | 0,00027579 | -2,32278539 |
| AT1G33770 | NA | Protein kinase | 0,00027857 | -3,80253117 |
| AT5G39330 | NA | Protein of unk | 0,00028021 | -5,34631095 |
| AT5G46520 | NA | Disease resista | 0,00028194 | -9,56295196 |
| AT3G46770 | NA | AP2/B3-like tr | 0,00028289 | -4,52457731 |
| AT3G11980 | FAR2 | Jojoba acyl Co | 0,00028374 | -4,61589391 |
| AT1G72800 | NA | RNA-binding ( | 0,00028374 | -2,49289475 |
| AT1G62150 | NA | Mitochondrial | 0,00028374 | -1,33051972 |
| AT5G64870 | NA | SPFH/Band 7/ | 0,00028588 | -2,13193757 |
| AT5G53030 | na | na | 0,00028898 | 1,98973132 |
| AT1G51380 | NA | DEA(D/H)-box | 0,00028913 | 1,11128311 |
| AT4G20725.1 | na | na | 0,00029241 | 8,64523357 |
| AT3G16400 | ATMLP-470 | nitrile specifie | 0,00029241 | 1,10470155 |
| AT1G56410 | ERD2 | heat shock pro | 0,00029241 | -3,37883426 |

| | | | | |
|---|---|---|---|---|
| AT4G11830 | PLDGAMMA2 | phospholipase | 0,0002935 | -1,15782687 |
| AT4G25220 | RHS15 | root hair spec | 0,00029448 | -4,0227424 |
| AT2G05410 | NA | TRAF-like fam | 0,00029483 | -8,41270792 |
| AT4G06744 | NA | Leucine-rich r | 0,00029483 | -1,1919645 |
| AT5G49800 | NA | Polyketide cyc | 0,00029517 | -1,47594118 |
| AT3G59220 | ATPIRIN1 | pirin | 0,00029585 | -3,28759351 |
| AT4G16140 | NA | proline-rich fa | 0,00029598 | -1,58128786 |
| AT5G50030 | NA | Plant invertas | 0,00029787 | -3,53475887 |
| AT1G27860 | NA | Protein of unk | 0,00030054 | -4,89881603 |
| AT1G32450 | NRT1.5 | nitrate transp | 0,00030099 | -2,70266318 |
| AT5G37860 | NA | Heavy metal t | 0,00030115 | -3,38888633 |
| AT4G23450 | NA | RING/U-box su | 0,00030282 | -8,66343077 |
| AT3G53640 | NA | Protein kinase | 0,00030371 | -3,36380513 |
| AT5G49550 | na | na | 0,00030434 | -1,22452741 |
| AT4G38870 | NA | F-box and ass | 0,00030491 | -7,73719853 |
| AT4G16050 | NA | Aminotransfer | 0,00030788 | -2,00928026 |
| AT2G34700 | NA | Pollen Ole e 1 | 0,0003079 | 3,48697557 |
| AT5G17240 | SDG40 | SET domain gr | 0,00030973 | -1,38839578 |
| AT5G38970 | ATBR6OX | brassinosteroi | 0,0003123 | -2,82807848 |
| AT1G61560 | ATMLO6 | Seven transm | 0,00031263 | 2,93128457 |
| AT2G30900 | TBL43 | TRICHOME BII | 0,00031263 | -1,7558103 |
| AT3G22920 | NA | Cyclophilin-lik | 0,00031265 | -4,56022698 |
| AT1G47480 | NA | alpha/beta-Hy | 0,00032338 | -3,19451505 |
| AT5G17165 | na | na | 0,00032945 | 1,60782595 |
| AT5G14810.1 | na | na | 0,00033153 | -5,91351463 |
| AT1G49900 | NA | C2H2 type zin | 0,00033153 | -5,30178593 |
| AT2G46480 | GAUT2 | galacturonosy | 0,00033153 | -2,42653498 |
| AT1G52155 | na | na | 0,00033378 | -1,67119271 |
| AT5G14000 | anac084 | NAC domain c | 0,00033411 | -2,19458712 |
| AT1G08050 | NA | Zinc finger (C3 | 0,00033448 | -1,38196589 |
| AT5G41130 | NA | Esterase/lipas | 0,00033531 | -1,62385586 |
| AT5G50480 | NF-YC6 | nuclear factor | 0,00033672 | -5,49172497 |
| AT5G02540 | NA | NAD(P)-bindir | 0,00033687 | -3,50907724 |

| | | | | |
|---|---|---|---|---|
| AT4G09200 | NA | SPla/RYanodir | 0,00034282 | -10,0041627 |
| AT1G66725 | na | na | 0,00034282 | -4,33087893 |
| AT1G80970 | NA | XH domain-co | 0,00034282 | -1,23866105 |
| AT1G25230 | NA | Calcineurin-lik | 0,00034282 | -1,0870426 |
| AT3G17570 | NA | F-box and ass | 0,0003446 | -7,98340528 |
| AT5G48420 | na | na | 0,0003459 | -2,41956973 |
| AT1G79800 | AtENODL7 | early nodulin- | 0,00035011 | -3,24993536 |
| AT4G30470 | NA | NAD(P)-bindir | 0,00035528 | -1,72181271 |
| AT1G08065 | ACA5 | alpha carboni | 0,00035677 | -4,30514917 |
| AT5G50120 | NA | Transducin/W | 0,00035891 | -3,09329973 |
| AT4G37690 | NA | Galactosyl tra | 0,00035891 | -1,83654733 |
| AT5G35370 | NA | S-locus lectin | 0,00035908 | 2,08323171 |
| AT5G28622.1 | na | na | 0,00035908 | -10,0047147 |
| AT2G33200 | NA | F-box family p | 0,00036115 | -4,82538438 |
| AT3G05890 | RCI2B | Low temperat | 0,00036298 | -1,88742992 |
| AT1G71110 | na | na | 0,00036298 | -1,16083909 |
| AT2G41780 | na | na | 0,00036417 | -2,176446 |
| AT4G22030 | NA | F-box family p | 0,00036429 | -5,42429333 |
| AT1G03920 | NA | Protein kinase | 0,00036486 | -1,519519 |
| AT4G13500 | na | na | 0,00036631 | -1,04130037 |
| AT1G59790 | NA | Cullin family p | 0,00036634 | -5,98974217 |
| AT4G09030 | AGP10 | arabinogalact | 0,00036912 | -1,9639137 |
| AT5G01600 | ATFER1 | ferretin 1 | 0,00036912 | -1,10143329 |
| AT1G11620 | NA | F-box and ass | 0,00036916 | -9,1136243 |
| AT3G22600 | NA | Bifunctional ir | 0,00037098 | -1,83063054 |
| AT3G60280 | UCC3 | uclacyanin 3 | 0,00037108 | -1,95164124 |
| AT4G13960 | NA | F-box/RNI-like | 0,00037756 | -8,35828805 |
| AT2G29940 | ATPDR3 | pleiotropic dru | 0,00038193 | -2,31280371 |
| AT2G23160 | NA | F-box family p | 0,0003844 | -4,57604727 |
| AT4G38880 | ASE3 | GLN phosphor | 0,00038524 | -5,01055238 |
| AT5G52170 | HDG7 | homeodomair | 0,00038713 | -1,35092677 |
| AT4G01740 | NA | Cysteine/Histi | 0,00038864 | -9,99431743 |
| AT5G27250.1 | na | na | 0,00038974 | -9,38116464 |

| | | | | |
|---|---|---|---|---|
| AT4G39290 | NA | Galactose oxid | 0,00038999 | -8,5951407 |
| AT5G12940 | NA | Leucine-rich r | 0,00039065 | -1,77069168 |
| AT1G07795 | na | na | 0,00039109 | -2,65652696 |
| AT5G10410 | NA | ENTH/ANTH/V | 0,00039395 | 2,52870242 |
| AT5G46490 | NA | Disease resist | 0,00039395 | -8,9944388 |
| AT1G66000 | NA | Family of unkn | 0,00039395 | -4,9008102 |
| AT2G39590 | NA | Ribosomal pro | 0,00039488 | -4,58457564 |
| AT1G71810 | NA | Protein kinase | 0,00039669 | -1,15197222 |
| AT4G38390 | RHS17 | root hair spec | 0,00039689 | -4,94822552 |
| AT5G34780 | NA | Thiamin diphc | 0,00039872 | -9,48100099 |
| AT4G24050 | NA | NAD(P)-bindir | 0,00039872 | -1,20327763 |
| AT2G44195 | NA | CBF1-interacti | 0,00039985 | -6,60239188 |
| AT1G71300 | NA | Vps52 / Sac2 f | 0,00040121 | -4,54542272 |
| AT2G36590 | ATPROT3 | proline transp | 0,00040386 | -1,29296113 |
| AT5G10420 | NA | MATE efflux fa | 0,00040521 | 1,80260727 |
| AT3G55540 | NA | nuclear transp | 0,00040521 | -4,86431876 |
| AT5G14070 | ROXY2 | Thioredoxin su | 0,00040521 | -2,14423039 |
| AT1G52347 | na | na | 0,00041285 | -9,84383285 |
| AT2G41640 | NA | Glycosyltransf | 0,00041285 | -2,24472501 |
| AT3G01030 | NA | C2H2 and C2H | 0,00041346 | -2,39257899 |
| AT5G58620 | NA | zinc finger (CC | 0,00041359 | -1,0302879 |
| AT1G60350 | anac024 | NAC domain c | 0,00041507 | -4,33099904 |
| AT2G35170 | NA | Histone H3 K4 | 0,00041652 | -1,14424806 |
| AT3G27620 | AOX1C | alternative ox | 0,00041673 | -2,03389678 |
| AT3G45460 | NA | IBR domain cc | 0,0004204 | -5,50506418 |
| AT2G18115 | na | na | 0,00042321 | -5,11920045 |
| AT5G24660 | LSU2 | response to lc | 0,00042321 | -4,44791118 |
| AT5G38565 | NA | F-box/FBD-like | 0,00042321 | -3,52411654 |
| AT5G08090 | na | na | 0,0004249 | -3,70492056 |
| AT2G03505 | NA | Carbohydrate- | 0,00042628 | -2,37212997 |
| AT5G07990 | CYP75B1 | Cytochrome P | 0,00043116 | -1,45699301 |
| AT1G72510 | NA | Protein of unk | 0,00043515 | -1,45798801 |
| AT4G38290 | NA | Haemolysin-II | 0,0004354 | 1,89518905 |

| | | | | |
|---|---|---|---|---|
| AT5G40040 | NA | 60S acidic ribo | 0,00043747 | -3,43721859 |
| AT1G63420 | NA | Arabidopsis th | 0,00043747 | -1,01004541 |
| AT5G48485 | DIR1 | Bifunctional ir | 0,00044174 | -1,74491781 |
| AT1G68840 | EDF2 | related to ABI | 0,00044367 | -2,59146443 |
| AT2G02990 | ATRNS1 | ribonuclease 1 | 0,00044449 | -2,63846538 |
| AT1G29570 | NA | Zinc finger C-x | 0,00044516 | -3,61993458 |
| AT5G19640 | NA | Major facilitat | 0,00044572 | -8,85853789 |
| AT4G16600 | NA | Nucleotide-di | 0,00044739 | 1,98049369 |
| AT1G25330 | NA | basic helix-loo | 0,00044739 | -5,14006552 |
| AT3G61230 | NA | GATA type zin | 0,00044739 | -3,33514061 |
| AT1G35560 | NA | TCP family tra | 0,00044739 | -1,35272969 |
| AT5G41310 | NA | P-loop nucleo | 0,00044848 | 5,09170949 |
| AT4G11900 | NA | S-locus lectin | 0,00044848 | -3,85032909 |
| AT1G13420 | ATST4B | sulfotransfera | 0,00044848 | -3,03648398 |
| AT5G58610 | NA | PHD finger tra | 0,00045159 | -1,36392719 |
| AT1G11110 | NA | LisH and RanB | 0,00045328 | -2,70209669 |
| AT5G10390 | NA | Histone super | 0,00045741 | 1,36741484 |
| AT1G58150 | na | na | 0,00045771 | -9,91638834 |
| AT3G18560 | na | na | 0,00045771 | -1,34688705 |
| AT2G17940 | NA | Plant protein | 0,00046357 | -7,99865142 |
| AT3G63095 | NA | Tetratricopep | 0,00046468 | -5,10774541 |
| AT3G59170 | NA | F-box/RNI-like | 0,00046468 | -4,7848544 |
| AT1G68460 | ATIPT1 | isopentenyltra | 0,00046493 | -2,58982461 |
| AT5G63760 | ARI15 | RING/U-box su | 0,000466 | -1,31723292 |
| AT5G38360 | NA | alpha/beta-Hy | 0,00046826 | -1,06265834 |
| AT1G77860 | KOM | Rhomboid-rel | 0,00047221 | -1,60246401 |
| AT1G62085 | NA | Mitochondrial | 0,00047276 | -1,14279197 |
| AT1G64030 | ATSRP3 | serpin 3 | 0,00047377 | -4,29293118 |
| AT5G02580 | NA | Plant protein 1 | 0,00047606 | 2,81950519 |
| AT5G05220 | na | na | 0,00047721 | -4,3236342 |
| AT2G32360 | NA | Ubiquitin-like | 0,00047855 | -2,71031853 |
| AT1G50630 | NA | Protein of unk | 0,00047855 | -1,96848742 |
| AT3G05770 | na | na | 0,00047926 | -7,97032963 |

| | | | | |
|---|---|---|---|---|
| AT5G54320 | NA | Protein of unk | 0,00047945 | -5,66160708 |
| AT4G12870 | NA | Gamma interf | 0,00048128 | 1,00937696 |
| AT2G34555 | ATGA2OX3 | gibberellin 2-c | 0,00048167 | -5,53277092 |
| AT4G18750 | DOT4 | Pentatricopep | 0,00048445 | 1,06198182 |
| AT3G59070 | NA | Cytochrome b | 0,00048445 | -5,61766823 |
| AT1G67220 | ATHPCAT1 | histone acetyl | 0,00048445 | -1,83964247 |
| AT5G54480 | NA | Protein of unk | 0,00048532 | -8,07792837 |
| AT2G35090 | NA | Protein of unk | 0,00048644 | -8,86505132 |
| AT2G36190 | AtcwINV4 | cell wall inver | 0,00048644 | -6,3456106 |
| AT3G11340 | NA | UDP-Glycosylt | 0,00048644 | -4,45228295 |
| AT4G39925 | NA | AT hook motif | 0,00048868 | -9,10091441 |
| AT3G12430 | NA | Polynucleotidy | 0,00048974 | -1,37610172 |
| AT4G28390 | AAC3 | ADP/ATP carri | 0,00048997 | -1,34561922 |
| AT1G61230 | NA | Mannose-binc | 0,00049296 | -8,85214306 |
| AT1G54940 | PGSIP4 | plant glycoger | 0,00049678 | -5,09069949 |
| AT2G24660.1 | na | na | 0,00049733 | -3,35120244 |
| AT2G44500 | NA | O-fucosyltrans | 0,00050067 | -1,74551828 |
| AT3G12420 | NA | Polynucleotidy | 0,00050193 | -1,55583657 |
| AT5G06230 | TBL9 | TRICHOME BII | 0,00050252 | -2,41857628 |
| AT3G58290 | NA | TRAF-like supe | 0,00050406 | -10,2060964 |
| AT5G42460 | NA | F-box and ass | 0,00050431 | 1,34669001 |
| AT2G05520 | ATGRP-3 | glycine-rich pr | 0,00050878 | 1,526111 |
| AT2G20800 | NDB4 | NAD(P)H dehy | 0,0005104 | -2,47553449 |
| AT1G34580 | NA | Major facilitat | 0,00051591 | 1,62309799 |
| AT1G68870 | ATSOFL2 | SOB five-like 2 | 0,00051591 | -1,46949648 |
| AT5G02200 | FHL | far-red-elonga | 0,00051786 | -2,65684153 |
| AT4G12100 | NA | Cullin family p | 0,00052084 | -3,76379216 |
| AT5G27360 | SFP2 | Major facilitat | 0,0005217 | -1,40932851 |
| AT1G49000 | na | na | 0,00052337 | -2,98197221 |
| AT2G45135 | NA | RING/U-box su | 0,00052424 | -3,1534008 |
| AT5G38096 | na | na | 0,00052479 | -8,70118316 |
| AT4G12940 | na | na | 0,00052479 | -6,79690272 |
| AT1G53790 | NA | F-box and ass | 0,00052479 | -2,33480199 |

| | | | | |
|---|---|---|---|---|
| AT3G22730 | NA | F-box and ass | 0,00052845 | -5,17355501 |
| AT3G53100 | NA | GDSL-like Lipa | 0,00053101 | -1,34085469 |
| AT4G26390 | NA | Pyruvate kina: | 0,00053143 | -3,84489071 |
| AT4G10950 | NA | SGNH hydrola | 0,00053209 | -2,12147731 |
| AT1G44890 | na | na | 0,00053633 | -1,19693155 |
| AT5G14930 | SAG101 | senescence-as | 0,00054585 | 1,45151675 |
| AT5G25250 | NA | SPFH/Band 7/ | 0,0005527 | -4,15357826 |
| AT2G03090 | ATEXP15 | expansin A15 | 0,00055359 | 1,3488231 |
| AT1G04570 | NA | Major facilitat | 0,00055509 | -1,31196357 |
| AT1G50520 | CYP705A27 | cytochrome P | 0,0005551 | -6,49144848 |
| AT3G13280 | NA | Putative endo | 0,0005551 | -4,98081811 |
| AT4G13110 | NA | BSD domain-c | 0,0005551 | -1,42097707 |
| AT5G38090 | na | na | 0,00055592 | -5,14386079 |
| AT5G42630 | ATS | Homeodomai | 0,00055592 | -2,66305005 |
| AT5G43160 | NA | Family of unk | 0,00055963 | 1,0234814 |
| AT1G52825 | na | na | 0,000561 | -2,35614725 |
| AT5G43790 | NA | Pentatricopep | 0,000561 | -1,70031737 |
| AT4G02380 | AtLEA5 | senescence-as | 0,00056295 | -2,21681208 |
| AT5G52250 | NA | Transducin/W | 0,00056494 | -2,14848877 |
| AT5G58160 | NA | actin binding | 0,0005662 | -1,03249833 |
| AT1G20925 | NA | Auxin efflux ca | 0,00056924 | -2,85509031 |
| AT3G60780 | NA | Protein of unk | 0,00057505 | -2,9138476 |
| AT3G58210 | NA | TRAF-like fam | 0,00057624 | -5,14104138 |
| AT2G26160 | NA | F-box family p | 0,00057664 | -8,12558188 |
| AT5G43860 | ATCLH2 | chlorophyllase | 0,00057857 | -2,64313056 |
| AT1G34640 | NA | peptidases | 0,0005802 | -1,21119611 |
| AT5G43620 | NA | Pre-mRNA cle | 0,00058487 | -2,34099832 |
| AT4G34000 | ABF3 | abscisic acid r | 0,00058487 | -1,99438406 |
| AT5G45990 | NA | crooked neck | 0,00058495 | -3,45702165 |
| AT5G35090 | na | na | 0,00059044 | -8,60696089 |
| AT2G30945 | na | na | 0,00059102 | -3,95176679 |
| AT3G48950 | NA | Pectin lyase-li | 0,00059104 | -3,8838174 |
| AT5G04275 | na | na | 0,00059182 | 3,0135927 |

| | | | | |
|---|---|---|---|---|
| AT2G47860 | NA | Phototropic-re | 0,00059182 | 1,14444133 |
| AT2G35160 | SGD9 | SU(VAR)3-9 ho | 0,00059396 | -1,10122378 |
| AT1G74020 | SS2 | strictosidine s | 0,00059399 | -1,13849332 |
| AT4G10000 | NA | Thioredoxin fa | 0,0005954 | 1,0698159 |
| AT5G63420 | emb2746 | RNA-metaboli | 0,00059847 | 1,02593277 |
| AT4G03370 | NA | Ubiquitin fam | 0,00059911 | -2,48161333 |
| AT5G05810 | ATL43 | RING/U-box su | 0,00060246 | -1,29262841 |
| AT5G37740 | NA | Calcium-depe | 0,00060876 | -1,38426951 |
| AT5G26570 | ATGWD3 | catalytics;carb | 0,00061021 | 1,00331626 |
| AT4G19830 | NA | FKBP-like pept | 0,00061021 | -1,18748679 |
| AT1G30180.1 | na | na | 0,00061415 | -4,3206882 |
| AT5G63180 | NA | Pectin lyase-li | 0,0006168 | 1,12609887 |
| AT1G07340 | ATSTP2 | sugar transpo | 0,0006168 | -6,73853864 |
| AT5G50200 | ATNRT3.1 | nitrate transm | 0,0006177 | -4,15469496 |
| AT1G61050 | NA | alpha 14-glyco | 0,0006183 | -3,3739301 |
| AT1G55560 | sks14 | SKU5 similar 1 | 0,00061879 | -4,29311268 |
| AT5G50240 | PIMT2 | protein-l-isoas | 0,00062005 | -2,4716744 |
| AT2G30230 | na | na | 0,00062014 | -1,46175113 |
| AT3G05780 | LON3 | lon protease 3 | 0,00062024 | -4,40559113 |
| AT2G45930 | na | na | 0,00062534 | -8,41441274 |
| AT1G19960 | na | na | 0,00062808 | 1,32534818 |
| AT5G06820 | SRF2 | STRUBBELIG-r | 0,00062992 | -2,03511636 |
| AT2G31470 | DOR | F-box and asso | 0,00063074 | -5,45238424 |
| AT1G69520 | NA | S-adenosyl-L-r | 0,00063088 | -1,77385912 |
| AT5G56020 | NA | Got1/Sft2-like | 0,0006356 | -1,09461018 |
| AT5G28442 | na | na | 0,00063842 | -9,4936715 |
| AT5G38200 | NA | Class I glutam | 0,0006399 | -3,469285 |
| AT3G12540 | NA | Protein of unk | 0,00064173 | -4,65805973 |
| AT1G53820 | NA | RING/U-box su | 0,00064257 | -4,54724323 |
| AT1G12600 | NA | UDP-N-acetylg | 0,00064366 | -2,2135661 |
| AT5G52640 | ATHS83 | heat shock pro | 0,0006446 | -1,52220113 |
| AT1G63390 | NA | FAD/NAD(P)-b | 0,00064881 | 2,74424428 |
| AT5G10770 | NA | Eukaryotic asp | 0,00064909 | -1,20211623 |

| | | | | |
|---|---|---|---|---|
| AT3G13210 | NA | crooked neck | 0,00064968 | -1,25667578 |
| AT5G28030 | DES1 | L-cysteine des | 0,00065162 | 1,06610716 |
| AT5G50790 | NA | Nodulin MtN3 | 0,00065707 | -1,20580137 |
| AT3G18720 | NA | F-box family p | 0,00066121 | -3,17978214 |
| AT3G45480 | NA | RING/U-box p | 0,00066489 | -5,25149515 |
| AT1G45145 | ATH5 | thioredoxin H- | 0,00066489 | -1,15808035 |
| AT1G66620 | NA | Protein with R | 0,0006752 | -1,55440261 |
| AT5G65100 | NA | Ethylene inser | 0,00067692 | -8,48681183 |
| AT1G70890 | MLP43 | MLP-like prote | 0,00067961 | 2,41664737 |
| AT4G37050 | AtPLAIVC | PATATIN-like p | 0,00067961 | -5,08954395 |
| AT5G17810 | WOX12 | WUSCHEL rela | 0,00067961 | -1,66608369 |
| AT5G39800 | NA | Mitochondrial | 0,00067961 | -1,4289858 |
| AT4G17580 | NA | Bax inhibitor-1 | 0,00067997 | -5,15440433 |
| AT2G32870 | NA | TRAF-like fam | 0,00068239 | 4,718186 |
| AT4G01450 | NA | nodulin MtN2 | 0,00068263 | -2,55804789 |
| AT4G13840 | NA | HXXXD-type a | 0,00068355 | 1,30019093 |
| AT1G60570 | NA | Galactose oxi | 0,00068511 | -8,46385325 |
| AT5G43170 | AZF3 | zinc-finger pro | 0,00068752 | -3,84705496 |
| AT1G53180 | na | na | 0,00068861 | -2,06956432 |
| AT5G20470 | NA | myosin putati | 0,00069102 | -8,52049565 |
| AT4G01975.1 | na | na | 0,00069419 | 1,45984236 |
| AT5G53048 | na | na | 0,00069419 | -6,19019586 |
| AT3G24850 | NA | Domain of unk | 0,00069419 | -3,38350184 |
| AT4G32500 | AKT5 | K+ transporte | 0,00069419 | -2,5171595 |
| AT1G79520 | NA | Cation efflux f | 0,00069419 | -1,72025242 |
| AT4G21330 | DYT1 | basic helix-loo | 0,00069707 | -2,20545316 |
| AT5G38010 | NA | UDP-Glycosylt | 0,00070193 | 3,09775644 |
| AT5G54190 | PORA | protochloroph | 0,0007057 | -5,40917765 |
| AT2G37170 | PIP2;2 | plasma memb | 0,00070573 | -1,6788426 |
| AT4G18580 | na | na | 0,00070573 | -1,60098584 |
| AT4G14301 | na | na | 0,00070573 | -3,81527981 |
| AT1G73410 | ATMYB54 | myb domain p | 0,00070573 | -3,3871481 |
| AT3G05320 | NA | O-fucosyltrans | 0,00070581 | -2,13623969 |

| | | | | |
|---|---|---|---|---|
| AT4G38000 | DOF4.7 | DNA binding v | 0,00070581 | -1,77826278 |
| AT5G43640 | NA | Ribosomal prc | 0,0007076 | -8,24758244 |
| AT1G15350 | na | na | 0,00070949 | -1,69762249 |
| AT2G34230 | NA | Protein with d | 0,00071343 | -1,93630688 |
| AT2G40050 | NA | Cysteine/Histi | 0,00071928 | 2,47705243 |
| AT4G12930 | na | na | 0,00073224 | -9,34212969 |
| AT5G51250 | NA | Galactose oxic | 0,00073668 | -4,91945077 |
| AT2G33220 | NA | GRIM-19 prot | 0,00073859 | -1,0239626 |
| AT1G09260 | NA | Chaperone Dr | 0,00073863 | -5,97628175 |
| AT5G43730 | NA | Disease resist | 0,00073991 | -3,40032972 |
| AT3G20440 | BE1 | Alpha amylase | 0,000749 | 1,07964183 |
| AT2G43510 | ATTI1 | trypsin inhibit | 0,0007516 | 1,73929325 |
| AT5G61340 | na | na | 0,00075789 | -1,87596534 |
| AT3G23860 | NA | GTP-binding p | 0,0007585 | -4,40669515 |
| AT3G28470 | ATMYB35 | Duplicated ho | 0,0007592 | -1,91092241 |
| AT4G15710 | na | na | 0,00076205 | -8,75089833 |
| AT1G03700 | NA | Uncharacteris | 0,00076205 | -1,97099902 |
| AT5G03440 | na | na | 0,00076205 | -1,20232889 |
| AT3G55590 | NA | Glucose-1-phc | 0,00076719 | -8,09223205 |
| AT4G21120 | AAT1 | amino acid tra | 0,00076824 | -5,4315837 |
| AT1G61300 | NA | LRR and NB-A | 0,00076991 | -1,14819785 |
| AT1G77655 | na | na | 0,00077088 | -4,62663155 |
| AT2G32160 | NA | S-adenosyl-L-r | 0,0007717 | -5,18015953 |
| AT1G74360 | NA | Leucine-rich r | 0,00077343 | -1,38123141 |
| AT1G28500 | NA | Protein of unk | 0,00078058 | -4,05891966 |
| AT1G48660 | NA | Auxin-respons | 0,0007926 | 4,87132351 |
| AT4G35733 | NA | F-box family p | 0,00079487 | -4,64449998 |
| AT5G38280 | PR5K | PR5-like recep | 0,00079643 | 2,7987106 |
| AT3G24280 | SMAP2 | small acidic pr | 0,00079716 | -4,50297326 |
| AT5G26160 | na | na | 0,00080083 | -1,35786153 |
| AT2G35830 | na | na | 0,00080083 | -1,06184922 |
| AT1G73810 | NA | Core-2/I-branc | 0,0008061 | -4,12330405 |
| AT3G61450 | ATSYP73 | syntaxin of pla | 0,00082582 | -1,83400461 |

| | | | | |
|---|---|---|---|---|
| AT4G01440 | NA | nodulin MtN2 | 0,00082733 | 1,0653764 |
| AT5G38396 | NA | F-box/RNI-like | 0,00082733 | -5,78740518 |
| AT4G28090 | sks10 | SKU5 similar : | 0,00083037 | -5,03811614 |
| AT5G09780 | NA | Transcriptiona | 0,00083037 | -1,51067948 |
| AT5G12330 | LRP1 | Lateral root pr | 0,00083164 | 1,2916016 |
| AT1G64640 | AtENODL8 | early nodulin- | 0,00083677 | -1,40096445 |
| AT1G23040 | NA | hydroxyprolin | 0,00083684 | -1,87679952 |
| AT1G20320 | NA | Haloacid deha | 0,00084779 | -3,80267174 |
| AT1G15850 | NA | Transducin/W | 0,00084779 | -3,09876092 |
| AT4G13800 | NA | Protein of unk | 0,00086146 | -3,65100839 |
| AT1G72290 | NA | Kunitz family t | 0,00087145 | -9,25558152 |
| AT1G04290 | NA | Thioesterase s | 0,00087749 | -9,06320101 |
| AT1G36150 | NA | Bifunctional ir | 0,00088617 | -6,78200025 |
| AT2G16790 | NA | P-loop contair | 0,00088617 | -2,76248777 |
| AT5G44569 | na | na | 0,00090278 | -9,26327892 |
| AT2G27310 | NA | F-box family p | 0,00090614 | -2,44335406 |
| AT4G18250 | NA | receptor serin | 0,00090754 | 4,11568149 |
| AT3G20557 | na | na | 0,00091369 | -2,80412359 |
| AT3G17460 | NA | PHD finger far | 0,00091743 | -1,06048742 |
| AT1G60525 | na | na | 0,00093216 | -9,25033601 |
| AT1G56040 | NA | HEAT/U-box d | 0,00093518 | 3,80887015 |
| AT3G28230 | na | na | 0,00093646 | -7,34383432 |
| AT1G22760 | PAB3 | poly(A) bindin | 0,0009365 | -4,0538499 |
| AT5G35480 | na | na | 0,00093713 | 1,92955854 |
| AT1G44608 | na | na | 0,00093765 | -3,45651827 |
| AT2G18540 | NA | RmlC-like cupi | 0,00094032 | 2,10960267 |
| AT2G31720 | NA | Domain of unl | 0,00094165 | -7,92515543 |
| AT1G74010 | NA | Calcium-depe | 0,00094165 | -2,14998861 |
| AT5G49290 | ATRLP56 | receptor like p | 0,00094277 | -5,02883273 |
| AT3G62440 | NA | F-box/RNI-like | 0,0009493 | -8,2743197 |
| AT4G35650 | IDH-III | isocitrate deh | 0,00095133 | -3,04140867 |
| AT1G32928 | na | na | 0,0009519 | -1,04084017 |
| AT1G55928 | NA | Coiled-coil dor | 0,00095789 | -3,52611821 |

| | | | | |
|---|---|---|---|---|
| AT2G27880 | AGO5 | Argonaute far | 0,00096935 | -1,3496231 |
| AT5G17830 | NA | Plasma-memb | 0,00096935 | -1,33164499 |
| AT5G37650 | NA | Family of unkı | 0,0009696 | -4,70519609 |
| AT3G24580 | NA | F-box and ass | 0,0009763 | -4,52416575 |
| AT3G52330 | NA | F-box associat | 0,00098082 | -7,67647005 |
| AT3G58390 | NA | Eukaryotic rel | 0,00098527 | -8,24845756 |
| AT1G74000 | SS3 | strictosidine s | 0,0009871 | 1,64367994 |
| AT3G22750 | NA | Protein kinase | 0,0009871 | -1,91006802 |
| AT2G39210 | NA | Major facilitat | 0,0009871 | -1,41725521 |
| AT5G04220 | ATSYTC | Calcium-depe | 0,00098713 | -1,41850396 |
| AT1G59700 | ATGSTU16 | glutathione S- | 0,00098713 | -1,12833735 |
| AT3G21040.1 | na | na | 0,00100225 | -1,68142274 |
| AT2G28090 | NA | Heavy metal t | 0,00101589 | -2,11901955 |
| AT2G36270 | ABI5 | Basic-leucine | 0,00102234 | -1,5325695 |
| AT3G16580 | NA | F-box and ass | 0,00102495 | -5,48239685 |
| AT1G80440 | NA | Galactose oxi | 0,00103257 | -2,47826288 |
| AT2G01990 | na | na | 0,00103595 | 1,01793908 |
| AT3G26940 | CDG1 | Protein kinase | 0,00103771 | -4,85211251 |
| AT3G13228 | NA | RING/U-box sı | 0,00104417 | -2,74070537 |
| AT1G07620 | ATOBGM | GTP-binding p | 0,00104901 | -4,325484 |
| AT2G23945 | NA | Eukaryotic asp | 0,00106215 | -4,47249838 |
| AT5G28635.1 | na | na | 0,00106524 | -5,84839187 |
| AT3G42850 | NA | Mevalonate/g | 0,00106814 | -3,26857067 |
| AT1G13370 | NA | Histone super | 0,00106993 | -2,17453044 |
| AT4G08115.1 | na | na | 0,00107139 | -9,16112483 |
| AT1G67650 | NA | SRP72 RNA-bi | 0,00107789 | -1,52390858 |
| AT3G21530 | NA | DNAse I-like sı | 0,00107793 | -2,23080845 |
| AT3G43910 | na | na | 0,00107931 | -5,47755388 |
| AT4G00320 | NA | F-box/RNI-like | 0,00108273 | -8,63368049 |
| AT3G15400 | ATA20 | anther 20 | 0,00112396 | -4,34037396 |
| AT2G32050 | NA | Family of unkı | 0,00113837 | -3,49482584 |
| AT5G44420 | LCR77 | plant defensir | 0,00113845 | 7,58777752 |
| AT2G01010 | na | na | 0,00114201 | 1,75551728 |

| | | | | |
|---|---|---|---|---|
| AT3G12580 | ATHSP70 | heat shock pr | 0,00114494 | -1,80382371 |
| AT2G23060 | NA | Acyl-CoA N-ac | 0,00114615 | 1,00800392 |
| AT1G49920 | NA | MuDR family t | 0,00114689 | -1,20871501 |
| AT1G72260 | THI2.1 | thionin 2.1 | 0,00115095 | -1,64592711 |
| AT1G15170 | NA | MATE efflux fa | 0,00115469 | -1,54741413 |
| AT5G65590 | NA | Dof-type zinc | 0,00115653 | 1,09398341 |
| AT5G52547 | na | na | 0,00115653 | -1,57540651 |
| AT1G63070 | NA | pentatricopep | 0,00116337 | 2,49781272 |
| AT3G55090 | NA | ABC-2 type tra | 0,00116337 | -1,93865911 |
| AT2G32905 | NA | Domain of unl | 0,0011647 | -2,48514386 |
| AT1G55740 | AtSIP1 | seed imbibitic | 0,0011647 | -1,65232104 |
| AT3G43710 | NA | Galactose oxi | 0,00117783 | -7,89773635 |
| AT1G66090 | NA | Disease resist | 0,00117812 | -9,11824037 |
| AT5G15660 | NA | F-box and ass | 0,0011836 | -4,96418627 |
| AT4G21130 | EMB2271 | Transducin/W | 0,00118582 | -4,66051246 |
| AT1G76480 | na | na | 0,00120052 | 2,22016618 |
| AT3G12830 | NA | SAUR-like aux | 0,00120611 | -1,28232797 |
| AT2G23330.1 | na | na | 0,00120839 | -1,26474848 |
| AT5G44480 | DUR | NAD(P)-bindir | 0,00120853 | -3,51035913 |
| AT2G40340 | AtERF48 | Integrase-type | 0,0012189 | 1,50793674 |
| AT1G48960 | NA | Adenine nucle | 0,0012189 | -1,28997531 |
| AT4G13570 | HTA4 | histone H2A 4 | 0,00123236 | 8,30290611 |
| AT5G56368 | na | na | 0,00123906 | -3,08656441 |
| AT1G65050 | NA | TRAF-like supe | 0,00124767 | -3,72635916 |
| AT5G16980 | NA | Zinc-binding d | 0,00125385 | -8,40529919 |
| AT1G14700 | ATPAP3 | purple acid ph | 0,00125513 | -1,00630102 |
| AT5G18290 | SIP1;2 | Aquaporin-like | 0,00125538 | -1,94808697 |
| AT2G01340 | At17.1 | NA | 0,0012587 | -5,64700485 |
| AT1G54000 | NA | GDSL-like Lipa | 0,00126081 | 1,09118659 |
| AT1G06100 | NA | Fatty acid des | 0,00126196 | 3,27132889 |
| AT1G67792 | na | na | 0,00126196 | -4,80450825 |
| AT4G39363 | na | na | 0,00126196 | -3,12252798 |
| AT3G60580 | NA | C2H2-like zinc | 0,00127373 | -1,22333437 |

| | | | | |
|---|---|---|---|---|
| AT2G35950 | EDA12 | embryo sac de | 0,0012774 | -2,11820735 |
| AT5G63350 | na | na | 0,00128049 | -1,66226771 |
| AT5G66070 | NA | RING/U-box su | 0,00128054 | -1,71116362 |
| AT1G65110 | NA | Ubiquitin carb | 0,00128772 | 1,56713812 |
| AT4G02830 | na | na | 0,00130227 | -2,94857863 |
| AT1G71280 | NA | DEA(D/H)-box | 0,00130753 | -4,14656002 |
| AT1G72100 | NA | late embryoge | 0,00130862 | 2,97027715 |
| AT5G16920 | NA | Fasciclin-like a | 0,00130862 | -6,07094636 |
| AT4G19620 | na | na | 0,00130914 | -3,05997944 |
| AT4G02670 | AtIDD12 | indeterminate | 0,00130914 | -2,14717608 |
| AT1G72890 | NA | Disease resista | 0,00130914 | -1,38923632 |
| AT4G10040 | CYTC-2 | cytochrome c- | 0,00130914 | -1,36444963 |
| AT5G53815.1 | na | na | 0,00130975 | -9,06740631 |
| AT5G40730 | AGP24 | arabinogalacta | 0,00130975 | -2,29524655 |
| AT3G45010 | scpl48 | serine carboxy | 0,00131753 | -1,89485481 |
| AT1G53023 | NA | Ubiquitin-conj | 0,00131771 | -3,44652011 |
| AT2G29950 | ELF4-L1 | ELF4-like 1 | 0,00131936 | -2,55641842 |
| AT5G44980 | NA | F-box/RNI-like | 0,00132626 | -6,70670041 |
| AT1G68390 | NA | Core-2/I-branc | 0,00133225 | -2,91292532 |
| AT3G09380 | NA | Protein of unk | 0,00133498 | -4,9643482 |
| AT5G01335.1 | na | na | 0,00133731 | -9,04043805 |
| AT4G22160 | na | na | 0,0013469 | -1,21516781 |
| AT5G27230 | NA | Frigida-like pr | 0,00135603 | -1,52929517 |
| AT2G48140 | EDA4 | Bifunctional ir | 0,00136277 | -4,44871531 |
| AT5G08460 | NA | GDSL-like Lipa | 0,00136691 | 4,0087934 |
| AT2G16760 | NA | Calcium-depe | 0,00136691 | -1,45591321 |
| AT1G16940 | NA | F-box/RNI-like | 0,0013769 | -7,8976564 |
| AT5G37415 | AGL105 | AGAMOUS-lik | 0,00138699 | -7,81378651 |
| AT3G04420 | anac048 | NAC domain c | 0,00138699 | -2,96315844 |
| AT5G53510 | ATOPT9 | oligopeptide t | 0,00138765 | -11,4481734 |
| AT1G21529 | na | na | 0,00138765 | -7,98103401 |
| AT2G41470 | na | na | 0,00138765 | -1,43210179 |
| AT5G02150 | Fes1C | Fes1C | 0,00140151 | 1,1523403 |

| | | | | |
|---|---|---|---|---|
| AT2G22030 | NA | Galactose oxi | 0,00140151 | -10,3350794 |
| AT5G52055.1 | na | na | 0,00140765 | -4,47839677 |
| AT4G25330 | na | na | 0,00141578 | 1,98748124 |
| AT1G61700 | NA | RNA polymera | 0,00141578 | -4,01204523 |
| AT3G29020 | AtMYB110 | myb domain p | 0,00141578 | -2,53586511 |
| AT1G23760 | JP630 | BURP domain- | 0,00142236 | -4,28193519 |
| AT3G17320 | NA | F-box and ass | 0,00142522 | -4,42767193 |
| AT2G23690 | na | na | 0,00144746 | -1,22221806 |
| AT5G57090 | AGR | Auxin efflux ca | 0,00145412 | 1,57871514 |
| AT2G35460 | NA | Late embryog | 0,0014638 | -8,03392845 |
| AT3G19595 | NA | Haloacid deha | 0,00146434 | -4,91001714 |
| AT4G21865 | na | na | 0,00147046 | -1,91040949 |
| AT2G36110 | NA | Polynucleotid | 0,00147568 | -8,78616694 |
| AT1G64035 | na | na | 0,00147568 | -4,97311057 |
| AT5G47530 | NA | Auxin-respons | 0,00148 | -5,13202379 |
| AT2G27220 | BLH5 | BEL1-like hom | 0,00148174 | 1,01493748 |
| AT5G66870 | ASL1 | ASYMMETRIC | 0,00148174 | -2,96082387 |
| AT4G20050 | QRT3 | Pectin lyase-li | 0,00149216 | -3,49133938 |
| AT5G17590 | NA | Putative mem | 0,00149772 | -3,39839178 |
| AT4G04460 | NA | Saposin-like a | 0,0015021 | -3,47658986 |
| AT5G44550 | NA | Uncharacteris | 0,00151401 | -2,67213194 |
| AT4G13030 | NA | P-loop contair | 0,00151479 | -4,67692895 |
| AT1G12380 | na | na | 0,00151479 | -1,04367447 |
| AT5G11360 | NA | Interleukin-1 r | 0,0015287 | -2,79251682 |
| AT5G41500 | NA | F-box and ass | 0,00153036 | -8,192008 |
| AT1G49150 | na | na | 0,00153378 | -3,66676343 |
| AT2G25810 | TIP4;1 | tonoplast intri | 0,00154102 | -1,41540124 |
| AT3G20760 | NA | Nse4 compon | 0,00154321 | -5,00026447 |
| AT5G08695 | NA | RNA-binding ( | 0,00154422 | -7,93041604 |
| AT3G49950 | NA | GRAS family tr | 0,00156193 | 2,50254733 |
| AT5G28145.1 | na | na | 0,00156284 | -9,19539918 |
| AT1G20790 | NA | F-box family p | 0,00157005 | -5,91437396 |
| AT5G55570 | na | na | 0,00158654 | 3,36841691 |

| | | | | |
|---|---|---|---|---|
| AT5G05490 | ATREC8 | Rad21/Rec8-li | 0,00159455 | -1,77098445 |
| AT5G47560 | ATSDAT | tonoplast dica | 0,00159838 | -1,61421541 |
| AT5G28262 | na | na | 0,00160445 | -5,02815781 |
| AT5G54206 | na | na | 0,00160713 | 2,5071105 |
| AT1G53080 | NA | Legume lectin | 0,00161247 | 1,49971426 |
| AT5G28430.1 | na | na | 0,00161247 | -9,26642732 |
| AT5G63730 | ARI14 | IBR domain-cc | 0,00161247 | -4,47274393 |
| AT5G05040 | NA | Cystatin/mon | 0,00161303 | -2,26169512 |
| AT4G06534 | na | na | 0,00162555 | -3,96061262 |
| AT3G10290 | NA | Nucleotide-su | 0,0016422 | -1,56618834 |
| AT1G49530 | GGPS6 | geranylgerany | 0,0016422 | -1,55281333 |
| AT4G36120 | NA | Plant protein | 0,0016422 | -1,11853873 |
| AT5G43340 | PHT1;6 | phosphate tra | 0,00164682 | -5,44055491 |
| AT5G02060 | NA | Uncharacteris | 0,00164803 | -2,64124996 |
| AT5G47600 | NA | HSP20-like ch | 0,00164877 | -1,01032657 |
| AT4G20140 | GSO1 | Leucine-rich r | 0,00166242 | -1,34296257 |
| AT4G34138 | UGT73B1 | UDP-glucosyl | 0,00169965 | 1,04212263 |
| AT5G51870 | AGL71 | AGAMOUS-lik | 0,00170039 | 1,592096 |
| AT4G13900 | na | na | 0,00170039 | -3,8272547 |
| AT1G76560 | CP12-3 | CP12 domain- | 0,00170448 | -1,43504584 |
| AT2G36550 | na | na | 0,00172082 | 4,32853177 |
| AT2G39435 | NA | Phosphatidyli | 0,00172238 | -1,00737197 |
| AT3G59480 | NA | pfkB-like carb | 0,00173096 | -3,7423461 |
| AT4G13810 | AtRLP47 | receptor like p | 0,00173863 | -5,70532984 |
| AT1G05330 | na | na | 0,00175959 | -2,81427827 |
| AT1G51402 | na | na | 0,00177 | -9,13280772 |
| AT1G76470 | NA | NAD(P)-bindir | 0,00177957 | -4,94365238 |
| AT1G70110 | NA | Concanavalin | 0,00178009 | -4,52223114 |
| AT3G11930 | NA | Adenine nucle | 0,00178009 | -1,1982008 |
| AT3G19850 | NA | Phototropic-r | 0,0017835 | -1,1008081 |
| AT1G14520 | MIOX1 | myo-inositol c | 0,00180083 | -2,04697491 |
| AT5G37750 | NA | Chaperone Dr | 0,00180515 | -8,64251948 |
| AT5G10600 | CYP81K2 | cytochrome P | 0,00180596 | -1,13960148 |

| | | | | |
|---|---|---|---|---|
| AT3G50570 | NA | hydroxyprolin | 0,0018184 | -3,98765549 |
| AT1G62560 | FMO GS-OX3 | flavin-monoo› | 0,00182665 | 1,33155374 |
| AT4G18300 | NA | Trimeric LpxA- | 0,00183122 | -1,01637652 |
| AT4G28530 | anac074 | NAC domain c | 0,00183161 | -1,77140272 |
| AT1G48820 | NA | Terpenoid cyc | 0,0018388 | -2,89544969 |
| AT3G16650 | NA | Transducin/W | 0,0018388 | -1,25036472 |
| AT5G15380 | DRM1 | domains rearr | 0,00184181 | -1,79761128 |
| AT3G14540 | NA | Terpenoid cyc | 0,00184405 | -3,98891175 |
| AT5G03285 | na | na | 0,00185496 | -2,28004239 |
| AT1G72140 | NA | Major facilitat | 0,00185733 | -3,64947827 |
| AT1G61080 | NA | Hydroxyprolin | 0,00185985 | -8,03393524 |
| AT2G21890 | ATCAD3 | cinnamyl alcol | 0,00185985 | -3,02957767 |
| AT3G55580 | NA | Regulator of c | 0,00186453 | -1,27360383 |
| AT4G15720 | NA | Tetratricopept | 0,00186742 | 1,6681684 |
| AT5G41761 | na | na | 0,00187069 | 3,58403596 |
| AT4G19840 | ATPP2-A1 | phloem prote | 0,00187076 | -1,17360836 |
| AT5G39910 | NA | Pectin lyase-li | 0,00187258 | -8,03589539 |
| AT4G38080 | NA | hydroxyprolin | 0,00187709 | 1,63180408 |
| AT1G80660 | AHA9 | H(+)-ATPase 9 | 0,00187709 | -3,79410909 |
| AT3G06545 | na | na | 0,00188188 | -5,49813487 |
| AT1G11740 | NA | ankyrin repeat | 0,00189143 | -3,17184949 |
| AT2G31050 | NA | Cupredoxin su | 0,00190834 | -8,03455277 |
| AT1G47565.1 | na | na | 0,00193183 | -5,8739265 |
| AT4G19130 | NA | Replication fa | 0,00193267 | -1,05269465 |
| AT1G65790 | ARK1 | receptor kinas | 0,00193853 | 1,99691854 |
| AT5G39410 | NA | Saccharopine | 0,00194711 | 1,02389389 |
| AT5G41835.1 | na | na | 0,00194861 | -8,87208882 |
| AT1G70170 | MMP | matrix metallo | 0,00195116 | -3,27488829 |
| AT4G25100 | ATFSD1 | Fe superoxide | 0,00196612 | 2,29250843 |
| AT4G18630 | NA | Protein of unk | 0,00196612 | 1,50314218 |
| AT5G17540 | NA | HXXXD-type a | 0,00199025 | -1,93192904 |
| AT1G16290 | na | na | 0,00199325 | -1,12404104 |
| AT5G37870 | NA | Protein with R | 0,00199599 | -8,43752237 |

| | | | | |
|---|---|---|---|---|
| AT2G40520 | NA | Nucleotidyltra | 0,00199968 | 1,06054062 |
| AT2G42170 | NA | Actin family p | 0,0020039 | 1,29814875 |
| AT5G60660 | PIP2;4 | plasma memb | 0,0020039 | -1,47594217 |
| AT1G79700 | NA | Integrase-type | 0,00201045 | -1,07734938 |
| AT3G50510 | LBD28 | LOB domain-c | 0,00203031 | -5,05582924 |
| AT3G18900 | na | na | 0,00203981 | -1,02558568 |
| AT1G35210 | na | na | 0,00204904 | 3,34578432 |
| AT1G15772 | na | na | 0,00205957 | -3,03694438 |
| AT5G43090 | APUM13 | pumilio 13 | 0,00205957 | -2,71083164 |
| AT5G46260 | NA | disease resista | 0,00207274 | 2,46356866 |
| AT2G43840 | UGT74F1 | UDP-glycosylt | 0,00207274 | -1,14302549 |
| AT4G33330 | GUX2 | plant glycoger | 0,00207636 | -1,56233703 |
| AT1G09880 | NA | Rhamnogalact | 0,00209026 | -2,91163488 |
| AT3G27710 | ARARI3 | RING/U-box su | 0,00209079 | -3,04943513 |
| AT5G53050 | NA | alpha/beta-Hy | 0,00209425 | -1,42928964 |
| AT1G67365 | na | na | 0,00210574 | -8,78565703 |
| AT1G61460 | NA | S-locus protei | 0,00211473 | 1,24490529 |
| AT4G32105 | NA | Beta-13-N-Ace | 0,00214879 | -2,23686953 |
| AT4G20100 | NA | PQ-loop repea | 0,00215873 | -5,23529585 |
| AT2G29600 | NA | Galactose oxi | 0,00216873 | 1,43949292 |
| AT3G54990 | SMZ | Integrase-type | 0,00217419 | -1,6870161 |
| AT5G66110 | NA | Heavy metal t | 0,00218105 | -3,21195585 |
| AT5G08260 | scpl35 | serine carboxy | 0,00218447 | 1,66716486 |
| AT1G75050 | NA | Pathogenesis- | 0,00219255 | -2,96729342 |
| AT5G56390 | NA | F-box/RNI-like | 0,00219755 | -8,60097179 |
| AT5G24790 | NA | Protein of unk | 0,00219837 | -4,76024554 |
| AT4G15975 | NA | RING/U-box su | 0,00221414 | -8,56539339 |
| AT1G69430 | na | na | 0,00223427 | -2,64161871 |
| AT5G67060 | HEC1 | basic helix-loo | 0,00224302 | -3,06799928 |
| AT4G20390 | NA | Uncharacteris | 0,00224302 | -2,21781073 |
| AT1G56360 | ATPAP6 | purple acid ph | 0,00225602 | -6,57134315 |
| AT2G28560 | ATRAD51B | DNA repair (R | 0,00226197 | 1,30992593 |
| AT3G04530 | ATPPCK2 | phosphoenolp | 0,00228023 | -2,57893806 |

| | | | | |
|---|---|---|---|---|
| AT3G26130 | NA | Cellulase (glyc | 0,00228208 | -2,45581114 |
| AT1G65780 | NA | P-loop contair | 0,00228837 | -2,26693143 |
| AT1G63380 | NA | NAD(P)-bindir | 0,00229168 | 1,72369253 |
| AT5G47510 | NA | Sec14p-like pł | 0,00230181 | -2,59261379 |
| AT2G33750 | ATPUP2 | purine perme | 0,00230791 | -3,42198341 |
| AT1G65200 | NA | Ubiquitin carb | 0,00232105 | 1,07369187 |
| AT5G50665 | na | na | 0,00233461 | -2,13763863 |
| AT5G07880 | ATSNAP29 | synaptosomal | 0,00233461 | -1,69412102 |
| AT1G13910 | NA | Leucine-rich r | 0,00235291 | -1,49114841 |
| AT1G43770 | NA | RING/FYVE/PH | 0,00236028 | -1,2265535 |
| AT4G39753 | NA | Galactose oxi | 0,00237133 | -4,93181396 |
| AT5G15030 | NA | Paired amphip | 0,00237793 | -1,02334112 |
| AT2G42480 | NA | TRAF-like fam | 0,00239899 | -5,42468298 |
| AT4G22280 | NA | F-box/RNI-like | 0,00240356 | 1,00370456 |
| AT3G11160 | na | na | 0,00241539 | -4,01125584 |
| AT3G12775 | NA | ubiquitin-conj | 0,00242111 | -4,4727073 |
| AT5G40220 | AGL43 | AGAMOUS-lik | 0,00242646 | -8,08670931 |
| AT3G32980 | NA | Peroxidase su | 0,00243809 | 3,46139945 |
| AT1G47980 | na | na | 0,00243994 | -5,37839029 |
| AT4G28150 | NA | Protein of unk | 0,00243994 | -1,14541857 |
| AT3G19270 | CYP707A4 | cytochrome P | 0,00245303 | 1,37592757 |
| AT4G36230 | na | na | 0,00246801 | -2,15984769 |
| AT2G29460 | ATGSTU4 | glutathione S- | 0,00249914 | -3,58562639 |
| AT5G33382.1 | na | na | 0,00250359 | -8,66123327 |
| AT3G44550 | FAR5 | fatty acid redu | 0,00251721 | -1,62404723 |
| AT4G34800 | NA | SAUR-like aux | 0,00252561 | -4,0296753 |
| AT4G36950 | MAPKKK21 | mitogen-activ | 0,00253367 | -1,85564264 |
| AT5G28415.1 | na | na | 0,00255525 | -8,5725496 |
| AT3G20660 | 04-oct | organic cation | 0,00255525 | -1,05246442 |
| AT1G53360 | NA | F-box associat | 0,00256527 | -7,80614446 |
| AT4G19000 | ATIWS2 | Transcription | 0,00256772 | -1,53446561 |
| AT4G34170 | NA | Galactose oxi | 0,00256801 | -4,59861672 |
| AT3G49020 | NA | FBD F-box anc | 0,00259641 | -4,48411426 |

| | | | | |
|---|---|---|---|---|
| AT2G16750 | NA | Protein kinase | 0,00259706 | -3,50945682 |
| AT5G27580 | AGL89 | AGAMOUS-lik | 0,00264112 | -8,58135838 |
| AT4G15460 | NA | glycine-rich pr | 0,00264439 | -3,54384891 |
| AT3G62520.1 | na | na | 0,00264558 | -8,6973975 |
| AT2G42410 | ATZFP11 | zinc finger prc | 0,00264558 | -1,57054502 |
| AT4G11750 | NA | Galactose oxic | 0,00265371 | -7,78193699 |
| AT1G73550 | NA | Bifunctional ir | 0,00265371 | -2,18093233 |
| AT5G23470 | NA | Haloacid deha | 0,00265859 | -4,7918401 |
| AT4G02960.1 | na | na | 0,00266013 | -8,61223481 |
| AT1G30710 | NA | FAD-binding B | 0,00266835 | -4,11407357 |
| AT1G49120 | NA | Integrase-type | 0,00266899 | -3,5143749 |
| AT5G59080 | na | na | 0,0026838 | -2,8635615 |
| AT1G65170 | NA | Ubiquitin carb | 0,00269912 | -3,1524051 |
| AT5G28910 | na | na | 0,00270091 | 1,09594205 |
| AT1G44191 | NA | ECA1 gametog | 0,00272342 | -6,68547391 |
| AT5G66830 | NA | F-box family p | 0,002727 | -1,9420501 |
| AT4G35090 | CAT2 | catalase 2 | 0,00274719 | -1,28008525 |
| AT2G31770 | ARI9 | RING/U-box su | 0,00274855 | -6,47091509 |
| AT5G52415 | na | na | 0,00276455 | -8,12779495 |
| AT4G12010 | NA | Disease resista | 0,00279502 | -1,54810594 |
| AT4G18360 | NA | Aldolase-type | 0,00279553 | -1,25696832 |
| AT4G01130 | NA | GDSL-like Lipa | 0,00280218 | 1,58991772 |
| AT3G44220 | NA | Late embryog | 0,00282095 | -1,41751678 |
| AT1G64720 | CP5 | Polyketide cyc | 0,00282722 | -1,02052113 |
| AT2G01560 | NA | Plant protein : | 0,00282822 | -2,55890218 |
| AT4G25010 | NA | Nodulin MtN3 | 0,00284063 | -1,3870425 |
| AT1G59850 | NA | ARM repeat su | 0,00284092 | -2,7359073 |
| AT1G64250.1 | na | na | 0,00284092 | -2,61559137 |
| AT4G14530 | na | na | 0,00286534 | -4,27843426 |
| AT1G34245 | EPF2 | Putative mem | 0,00286907 | 1,09595536 |
| AT5G17330 | GAD | glutamate dec | 0,00287377 | -1,8684122 |
| AT1G78955 | CAMS1 | camelliol C syr | 0,00287716 | -1,53445763 |
| AT4G16892 | na | na | 0,00288213 | 1,24576546 |

| | | | | |
|---|---|---|---|---|
| AT3G09540 | NA | Pectin lyase-li | 0,00288213 | 1,00775366 |
| AT5G42230 | scpl41 | serine carboxy | 0,00288606 | -1,45474762 |
| AT4G02650 | NA | ENTH/ANTH/V | 0,00290326 | -3,48179683 |
| AT5G53810 | NA | O-methyltrans | 0,00294066 | -4,23267831 |
| AT5G18090 | NA | AP2/B3-like tr | 0,00294066 | -2,78814268 |
| AT1G03340 | na | na | 0,00294286 | -1,72839531 |
| AT1G67370 | ASY1 | DNA-binding H | 0,00294627 | -1,03832866 |
| AT1G48150 | NA | MADS-box tra | 0,00294886 | -3,58663635 |
| AT4G18190 | ATPUP6 | purine perme | 0,0029566 | -7,46793561 |
| AT4G36590 | NA | MADS-box tra | 0,0029566 | -2,99725387 |
| AT4G17990 | na | na | 0,00296726 | -8,20466392 |
| AT1G52270 | na | na | 0,00298585 | -8,53345866 |
| AT5G55430 | na | na | 0,00298632 | -3,52740697 |
| AT3G08700 | UBC12 | ubiquitin-conj | 0,0029907 | -3,99343512 |
| AT3G62760 | ATGSTF13 | Glutathione S- | 0,00299377 | -3,98877473 |
| AT5G54660 | NA | HSP20-like ch | 0,00299377 | -1,28175961 |
| AT5G64540 | na | na | 0,00299803 | -1,69898634 |
| AT2G37880 | NA | Protein of unk | 0,00300846 | -1,09377834 |
| AT1G75920 | NA | GDSL-like Lipa | 0,0030117 | -4,40703202 |
| AT4G34650 | SQS2 | squalene synt | 0,00302317 | 1,18191079 |
| AT5G59810 | ATSBT5.4 | Subtilase fami | 0,00302317 | -6,12208839 |
| AT5G02600 | NA | Heavy metal t | 0,00302439 | -1,14582074 |
| AT4G24260 | ATGH9A3 | glycosyl hydrc | 0,00307562 | -1,31107324 |
| AT1G73510 | na | na | 0,00311508 | -4,70230405 |
| AT3G48450 | NA | RPM1-interac | 0,00312364 | -2,52388794 |
| AT2G36080 | NA | AP2/B3-like tr | 0,00312364 | -1,47897654 |
| AT1G04620 | NA | coenzyme F42 | 0,00312403 | 1,32097117 |
| AT1G31740 | BGAL15 | beta-galactosi | 0,00312955 | -8,89398822 |
| AT3G28960 | NA | Transmembra | 0,00314206 | -3,97466774 |
| AT1G09170 | NA | P-loop nucleo | 0,00315014 | 1,64108783 |
| AT4G24040 | ATTRE1 | trehalase 1 | 0,00315617 | -1,4452894 |
| AT3G53460 | CP29 | chloroplast RN | 0,00317312 | 1,12942057 |
| AT4G25070 | na | na | 0,00318869 | -1,5397598 |

| | | | | |
|---|---|---|---|---|
| AT5G48740 | NA | Leucine-rich r | 0,00319319 | 1,54873813 |
| AT1G03720 | NA | Cysteine prote | 0,00319438 | -2,11657664 |
| AT4G22790 | NA | MATE efflux fa | 0,00320718 | -1,3321877 |
| AT4G16990 | RLM3 | disease resista | 0,00320901 | 1,53009667 |
| AT4G04020 | FIB | fibrillin | 0,00322579 | -1,18229352 |
| AT4G06536 | NA | SPla/RYanodir | 0,00323766 | -3,96445985 |
| AT3G19055 | na | na | 0,00328083 | -7,62518352 |
| AT5G61270 | PIF7 | phytochrome- | 0,00328279 | -2,29455605 |
| AT5G46540 | PGP7 | P-glycoprotein | 0,00329207 | -2,36313368 |
| AT2G44700 | NA | Galactose oxic | 0,00329207 | -1,11443209 |
| AT5G52400 | CYP715A1 | cytochrome P | 0,00330957 | -3,57222376 |
| AT3G58160 | ATMYOS3 | P-loop contair | 0,00331756 | 1,03530606 |
| AT1G26590 | NA | C2H2-like zinc | 0,00331836 | -1,33282282 |
| AT5G22260 | MS1 | RING/FYVE/PH | 0,00333309 | -4,01909743 |
| AT2G35760 | NA | Uncharacteris | 0,00333545 | -1,35553416 |
| AT3G23955 | NA | F-box family p | 0,00334311 | -8,30704092 |
| AT1G64690 | BLT | branchless tric | 0,00336482 | -1,11229124 |
| AT1G49700 | NA | Plant protein 1 | 0,00339474 | -2,37531528 |
| AT5G37040 | NA | F-box family p | 0,00339595 | -3,69520438 |
| AT1G62270 | NA | F-box and asso | 0,00339659 | -4,13684232 |
| AT1G64060 | ATRBOH F | respiratory bu | 0,00340319 | -1,13313074 |
| AT4G30845 | na | na | 0,00340905 | 1,15612048 |
| AT4G37860 | NA | SPT2 chromat | 0,00340905 | -2,11713754 |
| AT1G14860 | atnudt18 | nudix hydrolas | 0,00341376 | -4,93338673 |
| AT2G38010 | NA | Neutral/alkalir | 0,00341376 | -1,25323603 |
| AT4G09480.1 | na | na | 0,00341484 | -8,37894731 |
| AT4G20810 | NA | transcription i | 0,0034149 | -5,11019635 |
| AT1G49720 | ABF1 | abscisic acid r | 0,003433 | 1,25408242 |
| AT3G50350 | NA | Protein of unk | 0,00343975 | -1,48872971 |
| AT3G50170 | NA | Plant protein o | 0,00344173 | 1,01025188 |
| AT3G23430 | ATPHO1 | phosphate 1 | 0,00344879 | -1,08882341 |
| AT2G35740 | ATINT3 | nositol transp | 0,0034884 | -2,00613646 |
| AT1G22150 | SULTR1;3 | sulfate transp | 0,003497 | -4,29577692 |

| | | | | |
|---|---|---|---|---|
| AT4G33900 | NA | Galactose oxi( | 0,003497 | -2,70698425 |
| AT5G03545 | AT4 | NA | 0,0035134 | -2,28855439 |
| AT5G28000 | NA | Polyketide cy( | 0,00352208 | 4,74681499 |
| AT4G15953 | NA | Maternally ex | 0,00352803 | -5,23942873 |
| AT3G14970 | NA | RING/U-box su | 0,00353192 | -4,2866365 |
| AT2G41445 | na | na | 0,00355795 | -8,34185854 |
| AT5G66480 | na | na | 0,00358251 | -1,81245871 |
| AT5G47450 | ATTIP2;3 | tonoplast intri | 0,00358572 | 4,64025764 |
| AT5G22170 | na | na | 0,00358689 | -8,12233909 |
| AT3G25080 | NA | Protein of unk | 0,0035873 | -4,05194762 |
| AT2G16720 | ATMYB7 | myb domain p | 0,00361256 | -1,40683611 |
| AT3G01311 | NA | Protein of unk | 0,00367017 | -4,26897373 |
| AT5G38620 | NA | MADS-box tra | 0,00367486 | -7,26820703 |
| AT4G16835 | NA | Tetratricopep | 0,00367999 | -2,08443972 |
| AT1G11980 | RUB3 | ubiquitin-relat | 0,00368742 | -4,39132283 |
| AT5G53635 | NA | F-box/RNI-like | 0,00372381 | -8,56343267 |
| AT1G66540 | NA | Cytochrome P | 0,0037273 | -8,14434051 |
| AT2G03200 | NA | Eukaryotic asp | 0,0037273 | -4,54821075 |
| AT3G08885 | na | na | 0,00373677 | -1,25522703 |
| AT3G47040 | NA | Glycosyl hydro | 0,00374107 | -7,70075508 |
| AT4G19570 | NA | Chaperone Dr | 0,00374107 | -2,077014 |
| AT4G38650 | NA | Glycosyl hydro | 0,00374247 | -1,29338987 |
| AT4G13820 | NA | Leucine-rich re | 0,00379435 | -2,54672259 |
| AT2G07707 | NA | Plant mitocho | 0,00381174 | 1,04262256 |
| AT3G15510 | ANAC056 | NAC domain c | 0,00381754 | -2,97681768 |
| AT5G10380 | ATRING1 | RING/U-box su | 0,00384788 | -1,12769808 |
| AT5G62970 | NA | Protein with R | 0,0038537 | -4,99262163 |
| AT4G16745 | NA | Exostosin fam | 0,00386336 | -2,66595488 |
| AT5G65205 | NA | NAD(P)-bindir | 0,003869 | -1,81749913 |
| AT2G19610 | NA | RING/U-box su | 0,00389012 | -3,66537616 |
| AT1G18830 | NA | Transducin/W | 0,00389844 | -2,36274321 |
| AT1G62810 | NA | Copper amine | 0,00391318 | -1,4083502 |
| AT5G28470 | NA | Major facilitat | 0,00392743 | -5,24676193 |

| | | | | |
|---|---|---|---|---|
| AT1G32140 | NA | F-box family p | 0,0039715 | -4,30207946 |
| AT3G62860 | NA | alpha/beta-Hy | 0,00397392 | -1,03945925 |
| AT1G47271 | NA | Cystathionine | 0,00399792 | 3,62418378 |
| AT2G01280 | MEE65 | Cyclin/Brf1-lik | 0,00402236 | -8,43354496 |
| AT5G17080 | NA | Cysteine prote | 0,00402236 | -8,43354496 |
| AT1G20657 | na | na | 0,00404299 | -4,06138179 |
| AT5G27200 | ACP5 | acyl carrier pr | 0,00405217 | 1,47361736 |
| AT5G59040 | COPT3 | copper transp | 0,00407199 | -2,90124874 |
| AT2G41415 | NA | Maternally ex | 0,00407209 | -6,0561655 |
| AT1G59780 | NA | NB-ARC doma | 0,0041051 | 1,69788111 |
| AT2G40750 | ATWRKY54 | WRKY DNA-bi | 0,00411599 | 1,2421485 |
| AT3G15830 | NA | phosphatidic a | 0,00414741 | -6,46394048 |
| AT2G34010 | na | na | 0,00416028 | -3,70722796 |
| AT5G11580 | NA | Regulator of c | 0,00416315 | -1,08379874 |
| AT3G16340 | ATPDR1 | pleiotropic dru | 0,0041723 | -1,75385978 |
| AT4G16960 | NA | Disease resista | 0,00420114 | -2,51910949 |
| AT4G23580 | NA | Galactose oxic | 0,00420296 | -4,93639101 |
| AT4G10440 | NA | S-adenosyl-L-r | 0,00420296 | -4,40821548 |
| AT5G14690 | na | na | 0,004216 | -2,8310677 |
| AT3G04050 | NA | Pyruvate kinas | 0,00423167 | -2,15325293 |
| AT2G44460 | BGLU28 | beta glucosida | 0,00423961 | -1,64669611 |
| AT2G01580 | na | na | 0,00424148 | -7,581949 |
| AT3G31023.1 | na | na | 0,0042969 | -4,58124053 |
| AT5G09950 | NA | Tetratricopept | 0,00429869 | 1,4366109 |
| AT3G03170 | na | na | 0,00432329 | 1,25725736 |
| AT1G23110 | na | na | 0,00432329 | -3,08789261 |
| AT2G44230 | NA | Plant protein ( | 0,00433524 | -1,2365979 |
| AT2G46130 | ATWRKY43 | WRKY DNA-bi | 0,00434395 | -9,37078454 |
| AT4G16840 | na | na | 0,00435663 | -1,58100195 |
| AT1G30660 | NA | nucleic acid bi | 0,00435663 | -1,15188717 |
| AT1G71330 | ATNAP5 | non-intrinsic A | 0,00437078 | 1,09117464 |
| AT1G20680 | NA | Protein of unk | 0,00438703 | -5,85273977 |
| AT5G48770 | NA | Disease resista | 0,00439317 | -8,22720647 |

| | | | | |
|---|---|---|---|---|
| AT5G45210 | NA | Disease resista | 0,00442266 | -3,52207929 |
| AT3G55150 | ATEXO70H1 | exocyst subur | 0,00446297 | -4,16482965 |
| AT1G50290 | na | na | 0,00446498 | -8,21196038 |
| AT1G65385 | na | na | 0,00447596 | -3,89512424 |
| AT1G65130 | NA | Ubiquitin carb | 0,00449093 | -1,16396915 |
| AT1G20132 | NA | GDSL-like Lipa | 0,00449238 | -5,03484145 |
| AT2G27280 | NA | Coiled-coil dor | 0,00453317 | -3,70647204 |
| AT5G49730 | ATFRO6 | ferric reductic | 0,00453317 | -1,24958685 |
| AT1G80130 | NA | Tetratricopept | 0,00454264 | 1,26863327 |
| AT4G10150 | NA | RING/U-box su | 0,00456383 | -2,25747144 |
| AT3G53150 | UGT73D1 | UDP-glucosyl t | 0,00461555 | -3,8549353 |
| AT5G45116.1 | na | na | 0,00462224 | -8,18095055 |
| AT5G27100 | ATGLR2.1 | glutamate rec | 0,00462374 | -1,32692075 |
| AT5G38035.1 | na | na | 0,00462968 | 1,21937078 |
| AT2G44170 | na | na | 0,00463993 | 1,21365438 |
| AT1G07090 | LSH6 | Protein of unk | 0,00463993 | -1,54195329 |
| AT1G47930 | na | na | 0,00465322 | -8,19178095 |
| AT2G24370 | NA | Protein kinase | 0,00470188 | -2,55725557 |
| AT1G70581 | na | na | 0,0047118 | -3,4493885 |
| AT4G05260 | NA | Ubiquitin-like | 0,00479039 | -3,41122366 |
| AT5G38870.1 | na | na | 0,00482103 | -1,58491751 |
| AT3G06750 | NA | hydroxyprolin | 0,00487949 | -1,10504545 |
| AT1G57730 | NA | RING/U-box su | 0,00491979 | -3,79837963 |
| AT4G16563 | NA | Eukaryotic asp | 0,00492076 | -2,55200584 |
| AT1G30170 | NA | Protein of unk | 0,00492076 | -1,37021775 |
| AT1G67600 | NA | Acid phosphat | 0,00495216 | -1,15114978 |
| AT3G59270 | NA | FBD-like doma | 0,0049804 | 1,66948905 |
| AT1G65180 | NA | Cysteine/Histi | 0,0050146 | 1,31071386 |
| AT4G00780 | NA | TRAF-like fam | 0,00507335 | -1,15022063 |
| AT2G32780 | ATUBP1 | ubiquitin-spec | 0,00508239 | -2,79261387 |
| AT2G13680 | ATGSL02 | callose syntha | 0,00508614 | -1,35847102 |
| AT4G33070 | NA | Thiamine pyrc | 0,00508614 | -1,30969288 |
| AT5G25390 | SHN2 | Integrase-type | 0,00508614 | -1,06543114 |

| | | | | |
|---|---|---|---|---|
| AT5G65080 | AGL68 | K-box region a | 0,00511134 | -3,19203159 |
| AT1G43910 | NA | P-loop contair | 0,00511306 | -1,24789813 |
| AT5G02980 | NA | Galactose oxic | 0,00511549 | -7,85528199 |
| AT5G11242 | na | na | 0,00518718 | -8,12022195 |
| AT5G48850 | ATSDI1 | Tetratricopep | 0,00520191 | -1,39176498 |
| AT1G73325 | NA | Kunitz family t | 0,00520447 | -1,34974159 |
| AT4G01883 | NA | Polyketide cyc | 0,00520693 | -1,43724573 |
| AT4G04080 | ATISU3 | ISCU-like 3 | 0,00526059 | -4,63399458 |
| AT1G71880 | ATSUC1 | sucrose-proto | 0,0052733 | 1,16915265 |
| AT3G05820 | At-A/N-InvH | invertase H | 0,00529115 | -1,55011781 |
| AT4G11530 | CRK34 | cysteine-rich I | 0,00529227 | 1,22882644 |
| AT2G40116 | NA | Phosphoinosit | 0,00529227 | -1,80408003 |
| AT2G39790 | NA | Mitochondrial | 0,00533034 | -4,85698344 |
| AT3G19040 | HAF2 | histone acetyl | 0,00534768 | -1,55357024 |
| AT5G26582.1 | na | na | 0,00538493 | 1,63759843 |
| AT1G06140 | NA | Pentatricopep | 0,00538844 | 1,14095705 |
| AT2G42005 | NA | Transmembra | 0,00539517 | -1,05474949 |
| AT3G57380 | NA | Glycosyltransf | 0,00542148 | -1,95638812 |
| AT5G10260 | AtRABH1e | RAB GTPase h | 0,00543028 | -1,13966772 |
| AT1G17060 | CYP72C1 | cytochrome p | 0,00547127 | -2,42032652 |
| AT1G23420 | INO | Plant-specific | 0,00547496 | -8,61186279 |
| AT4G16560 | NA | HSP20-like cha | 0,00552859 | -2,50563257 |
| AT4G08100.1 | na | na | 0,00555585 | -8,05453158 |
| AT1G56650 | ATMYB75 | production of | 0,00559134 | 2,12035506 |
| AT3G62670 | ARR20 | response regu | 0,00560369 | -3,27595079 |
| AT1G17710 | NA | Pyridoxal phos | 0,00562966 | -4,48402523 |
| AT3G45280 | ATSYP72 | syntaxin of pla | 0,00564419 | -2,90257651 |
| AT1G53260 | na | na | 0,00564419 | -2,00940894 |
| AT5G53110 | NA | RING/U-box su | 0,00567173 | -4,09953261 |
| AT1G53370 | NA | F-box and asso | 0,0056721 | -3,49242003 |
| AT2G43450 | na | na | 0,00568195 | -2,46083613 |
| AT3G57620 | NA | glyoxal oxidas | 0,0057322 | -6,79643983 |
| AT3G59880 | na | na | 0,0057322 | -3,09646466 |

| | | | | |
|---|---|---|---|---|
| AT5G26630 | NA | MADS-box tra | 0,0057322 | -2,01638345 |
| AT1G63150 | NA | Tetratricopep | 0,00573526 | -1,2398032 |
| AT4G01250 | AtWRKY22 | WRKY family t | 0,00574705 | -1,15841291 |
| AT3G60100 | CSY5 | citrate syntha | 0,00576334 | -8,1701663 |
| AT4G14746 | na | na | 0,00576334 | -1,05597564 |
| AT1G24625 | ZFP7 | zinc finger prc | 0,00577574 | 2,2662625 |
| AT5G41650 | NA | Lactoylglutath | 0,00577574 | -2,71204295 |
| AT3G14210 | ESM1 | epithiospecifi | 0,00583468 | 1,60228077 |
| AT5G24530 | DMR6 | 2-oxoglutarat | 0,00583468 | 1,46886727 |
| AT1G80480 | PTAC17 | plastid transcr | 0,00584858 | 1,06295557 |
| AT4G01430 | NA | nodulin MtN2 | 0,00586826 | -2,33191216 |
| AT5G25290 | NA | F-box family p | 0,00587651 | -5,41013124 |
| AT1G13140 | CYP86C3 | cytochrome P | 0,00588785 | -3,76043301 |
| AT2G46990 | IAA20 | indole-3-aceti | 0,00592013 | 1,50972906 |
| AT4G16890 | BAL | disease resista | 0,00594218 | -2,03038758 |
| AT5G18840 | NA | Major facilitat | 0,0059834 | 1,09894126 |
| AT1G14100 | FUT8 | fucosyltransfe | 0,00600574 | -4,29761223 |
| AT5G13880 | na | na | 0,00600889 | 1,07628726 |
| AT1G10070 | ATBCAT-2 | branched-cha | 0,00602487 | -1,10341844 |
| AT2G15770 | NA | Cupredoxin su | 0,00603227 | -4,73881449 |
| AT3G07070 | NA | Protein kinase | 0,0060625 | -4,15813987 |
| AT1G06920 | ATOFP4 | ovate family p | 0,00607173 | -2,38055695 |
| AT4G40010 | SNRK2-7 | SNF1-related p | 0,00610289 | -1,73868965 |
| AT1G50880 | NA | F-box and ass | 0,00613053 | -4,12388386 |
| AT3G15357 | na | na | 0,0061399 | -2,30813429 |
| AT3G28210 | PMZ | zinc finger (AN | 0,00630767 | -2,63330584 |
| AT5G18600 | NA | Thioredoxin su | 0,00631949 | -2,43273425 |
| AT1G49620 | ICK5 | Cyclin-depenc | 0,0064434 | -1,05219291 |
| AT5G05880 | NA | UDP-Glycosylt | 0,00653259 | -2,56087653 |
| AT5G23160 | na | na | 0,00653891 | -5,15626656 |
| AT4G25780 | NA | CAP (Cysteine | 0,00657674 | -2,46634333 |
| AT2G45403 | na | na | 0,00659461 | -2,89025559 |
| AT1G02520 | PGP11 | P-glycoprotein | 0,00659757 | -4,02762436 |

| | | | | |
|---|---|---|---|---|
| AT1G59860 | NA | HSP20-like cha | 0,00661604 | 1,52244821 |
| AT1G35690.1 | na | na | 0,00665565 | 1,758929 |
| AT5G28460 | NA | Pentatricopep | 0,00668813 | -7,92513881 |
| AT5G05870 | UGT76C1 | UDP-glucosyl t | 0,00678915 | 1,14724689 |
| AT1G76500 | AHL29 | Predicted AT-l | 0,00680072 | -2,7056443 |
| AT5G35110 | na | na | 0,00683483 | -4,3717975 |
| AT3G46520 | ACT12 | actin-12 | 0,00684045 | -10,0371259 |
| AT5G41860 | na | na | 0,00687726 | -1,82979535 |
| AT2G43960 | NA | SWAP (Suppre | 0,00691106 | -4,26335945 |
| AT4G14690 | ELIP2 | Chlorophyll A- | 0,00693175 | -1,56727027 |
| AT1G73260 | ATKTI1 | kunitz trypsin | 0,00697637 | 3,4815127 |
| AT3G22240 | na | na | 0,00697949 | -4,06839427 |
| AT3G22886 | na | na | 0,00703312 | -3,24407833 |
| AT2G46880 | ATPAP14 | purple acid ph | 0,00705779 | -7,25651479 |
| AT5G09500 | NA | Ribosomal pro | 0,00708241 | -5,59835733 |
| AT4G30830 | NA | Protein of unk | 0,00710937 | -1,60748399 |
| AT1G76430 | PHT1;9 | phosphate tra | 0,00712431 | -3,19918083 |
| AT1G24590 | DRN-LIKE | DORNROSCHE | 0,0071644 | 1,81276077 |
| AT5G57240 | ORP4C | OSBP(oxyster | 0,00721728 | -4,88227727 |
| AT4G13260 | YUC2 | Flavin-binding | 0,00736661 | 1,16270451 |
| AT2G04100 | NA | MATE efflux fa | 0,00737634 | -1,79206642 |
| AT5G14980 | NA | alpha/beta-Hy | 0,0073971 | -7,42253295 |
| AT5G61940 | NA | Ubiquitin carb | 0,0074148 | -1,1832809 |
| AT5G41200 | AGL75 | AGAMOUS-lik | 0,00745243 | -2,96479538 |
| AT5G51530 | NA | Ubiquitin carb | 0,00745518 | -2,19442227 |
| AT2G18550 | ATHB21 | homeobox pro | 0,00747148 | 2,33157082 |
| AT5G39471 | NA | Cysteine/Histi | 0,00748597 | -4,18309403 |
| AT2G42720 | NA | FBD F-box Skp | 0,00748755 | -1,40307823 |
| AT4G27420 | NA | ABC-2 type tra | 0,00751788 | -7,04228172 |
| AT3G61390 | NA | RING/U-box su | 0,00752728 | -3,69693479 |
| AT3G53590 | NA | Leucine-rich re | 0,00752728 | -2,77050296 |
| AT2G21260 | NA | NAD(P)-linked | 0,00758311 | -1,11027258 |
| AT2G01200 | IAA32 | indole-3-aceti | 0,0075916 | 2,30942274 |

| | | | | |
|---|---|---|---|---|
| AT5G59930 | NA | Cysteine/Histi | 0,00765063 | -7,94654032 |
| AT5G57190 | PSD2 | phosphatidyls | 0,00773701 | -2,93908129 |
| AT5G46270 | NA | Disease resista | 0,00774017 | 1,28137547 |
| AT1G20150 | NA | Subtilisin-like | 0,00774732 | -8,36365804 |
| AT1G47655 | NA | Dof-type zinc | 0,00774732 | -1,14159466 |
| AT3G22080 | NA | TRAF-like fam | 0,00777616 | -4,86929398 |
| AT3G49690 | ATMYB84 | myb domain p | 0,00789327 | -1,0698337 |
| AT2G03000 | NA | RING/U-box su | 0,00790851 | -3,54744539 |
| AT4G14815 | NA | Bifunctional ir | 0,00791392 | -7,40441073 |
| AT3G49055 | na | na | 0,00794133 | -1,71184081 |
| AT1G28700 | NA | Nucleotide-dip | 0,00796612 | -6,88745435 |
| AT4G19645 | NA | TRAM LAG1 ar | 0,00798881 | -3,3291667 |
| AT4G22660 | NA | F-box family p | 0,0079948 | -8,2108018 |
| AT1G17950 | ATMYB52 | myb domain p | 0,00804309 | -2,85966186 |
| AT4G28840 | na | na | 0,00810226 | -3,07345235 |
| AT4G27460 | NA | Cystathionine | 0,00811307 | -1,13190359 |
| AT1G68862 | na | na | 0,00811762 | -1,19780432 |
| AT3G46370 | NA | Leucine-rich re | 0,00813253 | -7,7669795 |
| AT5G38750 | NA | asparaginyl-tR | 0,00819506 | -2,68795321 |
| AT3G25882 | NIMIN-2 | NIM1-interact | 0,00820468 | 1,06419865 |
| AT1G54540 | NA | Late embryog | 0,00820468 | -4,56630413 |
| AT2G03590 | ATUPS1 | ureide permea | 0,0082053 | -1,22937748 |
| AT4G39180 | ATSEC14 | Sec14p-like ph | 0,00821622 | -1,35646902 |
| AT5G20810 | NA | SAUR-like aux | 0,00823465 | -1,26538222 |
| AT5G28650 | ATWRKY74 | WRKY DNA-bi | 0,00824263 | -1,5605053 |
| AT5G60140 | NA | AP2/B3-like tr | 0,00826872 | -4,18019658 |
| AT3G28270 | NA | Protein of unk | 0,00826872 | -1,83937626 |
| AT1G01180 | NA | S-adenosyl-L-r | 0,00826872 | -1,12069221 |
| AT5G17360 | na | na | 0,00827817 | -1,08181362 |
| AT1G80180 | na | na | 0,0082886 | -1,28009202 |
| AT2G31035 | na | na | 0,00829622 | -1,1708086 |
| AT5G16340 | NA | AMP-depende | 0,00833964 | -1,82347786 |
| AT5G26900 | NA | Transducin far | 0,0085174 | -3,68368483 |

| | | | | |
|---|---|---|---|---|
| AT1G19640 | JMT | jasmonic acid | 0,0086009 | -1,99187281 |
| AT3G59700 | ATHLECRK | lectin-recepto | 0,00863977 | -1,56502427 |
| AT4G18090 | na | na | 0,00865491 | -2,41731893 |
| AT1G02813 | NA | Protein of unk | 0,0086609 | -6,55038273 |
| AT3G52720 | ACA1 | alpha carboni | 0,00867291 | -1,90411698 |
| AT3G43600 | AAO2 | aldehyde oxid | 0,00868225 | 1,02194097 |
| AT1G09080 | BIP3 | Heat shock pr | 0,00868238 | -4,9448997 |
| AT2G43930 | NA | Protein kinase | 0,00877201 | -1,38786745 |
| AT2G33100 | ATCSLD1 | cellulose syntl | 0,00879347 | -4,60029617 |
| AT4G36640 | NA | Sec14p-like pl | 0,00882136 | -1,52513627 |
| AT1G72110 | NA | O-acyltransfer | 0,00884655 | -1,87874217 |
| AT1G74290 | NA | alpha/beta-Hy | 0,00888774 | 2,2957082 |
| AT3G20340 | na | na | 0,00889964 | -1,88617168 |
| AT1G50830 | NA | Aminotransfer | 0,00891562 | -2,60180338 |
| AT2G23130 | AGP17 | arabinogalact | 0,00899017 | -1,24762987 |
| AT5G16360 | NA | NC domain-co | 0,00900816 | -2,09716759 |
| AT5G49070 | KCS21 | 3-ketoacyl-Co | 0,00911659 | -3,54374098 |
| AT4G21600 | ENDO5 | endonuclease | 0,00918821 | -1,5029893 |
| AT1G49490 | NA | Leucine-rich r | 0,00924326 | -1,01528662 |
| AT4G13560 | UNE15 | Late embryog | 0,00927271 | 3,11367853 |
| AT4G38030 | NA | Rhamnogalact | 0,00931937 | -3,42442914 |
| AT2G17500 | NA | Auxin efflux ca | 0,00934222 | -1,10442695 |
| AT2G26520 | na | na | 0,00939008 | -1,4936447 |
| AT5G47220 | ATERF-2 | ethylene resp | 0,00940743 | -2,64735352 |
| AT1G35310 | MLP168 | MLP-like prot | 0,00953309 | -3,55839905 |
| AT1G48060 | NA | F-box and ass | 0,00955225 | -1,66996869 |
| AT3G61410 | na | na | 0,00956739 | -1,37956 |
| AT2G44745 | NA | WRKY family t | 0,00967239 | 1,05821228 |
| AT4G28397 | na | na | 0,0096737 | -8,28461962 |
| AT2G35736 | na | na | 0,00970462 | -1,20551117 |
| AT2G21650 | ATRL2 | Homeodomai | 0,00972247 | -2,36890573 |
| AT4G22410 | NA | Ubiquitin C-te | 0,00974331 | 1,76139344 |
| AT2G43610 | NA | Chitinase fami | 0,00974331 | 1,44612871 |

| | | | | |
|---|---|---|---|---|
| AT2G15830 | na | na | 0,00974331 | 1,18043076 |
| AT1G69630 | NA | F-box/RNI-like | 0,00974331 | -2,56630616 |
| AT5G41690 | NA | RNA-binding ( | 0,00974331 | -1,40594151 |
| AT5G02070 | NA | Protein kinase | 0,00974331 | -1,30104702 |
| AT2G42460 | NA | TRAF-like fam | 0,00978967 | -3,30439236 |
| AT1G69990 | NA | Leucine-rich re | 0,00981266 | -2,66968777 |
| AT5G46780 | NA | VQ motif-cont | 0,00983113 | 1,00136449 |
| AT5G36260 | NA | Eukaryotic asp | 0,00984717 | 1,42899042 |
| AT2G22750 | NA | basic helix-loc | 0,00984735 | -4,5954968 |
| AT1G15770 | na | na | 0,00984735 | -2,72005822 |
| AT2G18150 | NA | Peroxidase su | 0,00984735 | -1,61374795 |
| AT1G64700 | na | na | 0,00984735 | -1,58681989 |
| AT3G27440 | UKL5 | uridine kinase | 0,00987424 | -2,17090551 |
| AT1G33700 | NA | Beta-glucosida | 0,00987424 | -1,4324506 |
| AT1G10385 | NA | Vps51/Vps67 | 0,00989047 | -2,236843 |
| AT1G50040 | NA | Protein of unk | 0,00993154 | -1,02694793 |
| AT1G33740 | na | na | 0,0100142 | -8,27431411 |
| AT5G62320 | ATMYB99 | myb domain p | 0,01002433 | -5,57068266 |
| AT1G18400 | BEE1 | BR enhanced e | 0,01003879 | -2,22787302 |
| AT1G43800 | NA | Plant stearoyl- | 0,01012964 | 1,63333701 |
| AT4G19690 | ATIRT1 | iron-regulated | 0,01015908 | -8,87169978 |
| AT5G17740 | NA | P-loop contair | 0,01015908 | -8,08849589 |
| AT5G16200 | NA | 50S ribosomal | 0,01018367 | -1,04173385 |
| AT1G72900 | NA | Toll-Interleuki | 0,01025084 | -1,67233052 |
| AT2G37260 | ATWRKY44 | WRKY family t | 0,01025301 | 1,02492303 |
| AT3G62040 | NA | Haloacid deha | 0,01029544 | -2,29855891 |
| AT1G08165 | na | na | 0,01030027 | -1,59395208 |
| AT4G28395 | A7 | Bifunctional ir | 0,01041277 | -6,64335648 |
| AT5G02420 | na | na | 0,01042829 | -1,84403571 |
| AT3G20450 | NA | B-cell recepto | 0,01043253 | -4,51405643 |
| AT2G16905 | na | na | 0,01053914 | -6,46932484 |
| AT1G55035 | na | na | 0,01056932 | -3,80790064 |
| AT3G55240 | NA | Plant protein 1 | 0,01062711 | 1,21438876 |

| | | | | |
|---|---|---|---|---|
| AT3G46970 | ATPHS2 | alpha-glucan p | 0,01065641 | 1,15214331 |
| AT4G26830 | NA | O-Glycosyl hy | 0,01070608 | -2,92799529 |
| AT5G24900 | CYP714A2 | cytochrome P | 0,01076795 | 2,02298473 |
| AT2G03020 | NA | Heat shock pr | 0,01078673 | -1,11047976 |
| AT4G08710.1 | na | na | 0,01087347 | 7,76008186 |
| AT3G16460 | NA | Mannose-binc | 0,01089757 | -2,59113184 |
| AT2G03740 | NA | late embryoge | 0,01092132 | -6,84973306 |
| AT5G48690 | na | na | 0,01096285 | -1,62651226 |
| AT1G58270 | ZW9 | TRAF-like fam | 0,01104856 | -3,71292611 |
| AT4G06746 | DEAR5 | related to AP2 | 0,01107154 | -2,54825118 |
| AT4G04745 | na | na | 0,01110727 | -4,9847287 |
| AT1G45191 | BGLU1 | Glycosyl hydrc | 0,01120081 | -1,21487742 |
| AT3G09922 | ATIPS1 | induced by ph | 0,01122867 | -3,71212802 |
| AT1G02030 | NA | C2H2-like zinc | 0,01123911 | -1,05313937 |
| AT1G79270 | ECT8 | evolutionarily | 0,01128992 | -1,19154481 |
| AT1G23560 | NA | Domain of unl | 0,01131703 | -7,27915416 |
| AT3G45060 | ATNRT2.6 | high affinity n | 0,01132122 | -2,94487695 |
| AT3G28830 | NA | Protein of unk | 0,01137682 | -4,53101524 |
| AT1G32992 | na | na | 0,01138791 | -2,90026262 |
| AT4G03364 | na | na | 0,01138791 | -2,06067479 |
| AT2G18600 | NA | Ubiquitin-conj | 0,01143842 | -1,37869447 |
| AT1G04090 | NA | Plant protein | 0,01145403 | -1,15654976 |
| AT2G28440 | NA | proline-rich fa | 0,01145901 | -1,31867858 |
| AT4G18335 | na | na | 0,01149723 | -2,30840216 |
| AT1G52660 | NA | P-loop contair | 0,01164829 | -2,4691854 |
| AT4G11460 | CRK30 | cysteine-rich I | 0,0116839 | 2,27556026 |
| AT1G13600 | AtbZIP58 | basic leucine-z | 0,01169866 | 1,11019666 |
| AT1G19380 | NA | Protein of unk | 0,01176978 | -1,85743848 |
| AT2G22510 | NA | hydroxyprolin | 0,01181241 | -4,08292812 |
| AT2G32300 | UCC1 | uclacyanin 1 | 0,01198895 | -2,74113862 |
| AT4G25300 | NA | 2-oxoglutarat | 0,011992 | -1,36309644 |
| AT5G55090 | MAPKKK15 | mitogen-activ | 0,0120065 | -1,53196028 |
| AT1G73340 | NA | Cytochrome P | 0,01203565 | -3,07309438 |

| | | | | |
|---|---|---|---|---|
| AT4G36610 | NA | alpha/beta-Hy | 0,01209764 | -1,14069652 |
| AT5G54010 | NA | UDP-Glycosylt | 0,01211838 | -6,53933038 |
| AT1G51800 | NA | Leucine-rich r | 0,01213146 | -3,57759692 |
| AT4G18510 | CLE2 | CLAVATA3/ES | 0,01213554 | -3,58966489 |
| AT2G31100 | NA | alpha/beta-Hy | 0,01222084 | -2,38104744 |
| AT4G20070 | AAH | allantoate am | 0,01224216 | -1,39988222 |
| AT4G01230 | NA | Reticulon fam | 0,01239455 | -2,05370911 |
| AT4G01170 | na | na | 0,01241133 | -2,76417233 |
| AT3G10340 | PAL4 | phenylalanine | 0,01244758 | -1,90511481 |
| AT2G02890 | NA | F-box family p | 0,01246672 | -7,26233532 |
| AT5G16370 | AAE5 | acyl activating | 0,01248301 | -1,03231362 |
| AT3G59230 | NA | RNI-like super | 0,01254621 | -3,65720323 |
| AT5G22490 | NA | O-acyltransfer | 0,01257574 | 3,19017519 |
| AT4G24890 | ATPAP24 | purple acid ph | 0,01259584 | -4,94479731 |
| AT2G28680 | NA | RmlC-like cupi | 0,01261812 | -7,39373882 |
| AT1G32660 | NA | F-box and ass | 0,0126415 | -4,10435754 |
| AT5G02020 | na | na | 0,01274023 | -1,29443867 |
| AT3G62280 | NA | GDSL-like Lipa | 0,01278184 | -1,52941968 |
| AT4G24230 | ACBP3 | acyl-CoA-bind | 0,01278184 | -1,02367774 |
| AT5G57500 | NA | Galactosyltrar | 0,01285225 | -1,97252894 |
| AT2G47360 | na | na | 0,01295437 | -1,72404803 |
| AT5G11100 | ATSYTD | Calcium-depe | 0,01295437 | -1,11625686 |
| AT3G52160 | KCS15 | 3-ketoacyl-Co | 0,01303597 | -6,28946588 |
| AT4G37260 | ATMYB73 | myb domain p | 0,01309724 | -1,05117759 |
| AT4G09432 | na | na | 0,01310183 | -1,45348615 |
| AT4G35680 | NA | Arabidopsis pr | 0,01310608 | -1,61341229 |
| AT5G52620 | NA | F-box associat | 0,01314164 | -8,16078173 |
| AT1G53410 | na | na | 0,01319314 | -2,31630823 |
| AT2G37435 | NA | Cystatin/mon | 0,01328759 | -2,77880274 |
| AT5G59845 | NA | Gibberellin-re | 0,01334239 | -3,9179379 |
| AT4G26770 | NA | Phosphatidate | 0,01335259 | -1,60525841 |
| AT1G52890 | ANAC019 | NAC domain c | 0,01336893 | -1,02424389 |
| AT1G21940 | na | na | 0,01343777 | -2,24080894 |

| | | | | |
|---|---|---|---|---|
| AT3G06970 | NA | RNA-binding ( | 0,01353633 | -7,88356941 |
| AT1G16530 | ASL9 | ASYMMETRIC | 0,01363009 | 1,74949009 |
| AT3G22910 | NA | ATPase E1-E2 | 0,01363375 | -1,56769896 |
| AT5G56325 | NA | FBD-like doma | 0,01368905 | -6,65601344 |
| AT1G18280 | NA | Bifunctional ir | 0,01372898 | -1,55536587 |
| AT1G27080 | NRT1.6 | nitrate transp | 0,01374282 | -1,47717304 |
| AT5G55490 | ATGEX1 | gamete expre | 0,01379884 | -1,43861351 |
| AT5G62720 | NA | Integral meml | 0,01380704 | 1,46665687 |
| AT1G77210 | AtSTP14 | sugar transpo | 0,01381203 | -4,16802847 |
| AT2G29410 | ATMTPB1 | metal toleran | 0,01386469 | -2,84265972 |
| AT5G24380 | ATYSL2 | YELLOW STRIF | 0,0138906 | -1,69226417 |
| AT3G15740 | NA | RING/U-box su | 0,01389775 | -2,433278 |
| AT1G52650 | NA | F-box/RNI-like | 0,0139436 | 1,19702439 |
| AT4G00040 | NA | Chalcone and | 0,0139725 | -1,18513236 |
| AT4G18290 | KAT2 | potassium cha | 0,01397602 | -1,41871044 |
| AT4G10940 | NA | RING/U-box p | 0,01401 | 1,68601364 |
| AT4G01540 | ANAC068 | NAC with tran | 0,01403212 | -2,34617546 |
| AT3G17760 | GAD5 | glutamate dec | 0,01416736 | -2,57852109 |
| AT5G44020 | NA | HAD superfan | 0,01425724 | -1,0808049 |
| AT1G09720 | NA | Kinase interac | 0,01427024 | -2,80433877 |
| AT4G08670 | NA | Bifunctional ir | 0,01429626 | -5,25013489 |
| AT5G39240 | na | na | 0,01432154 | 1,19137381 |
| AT1G13110 | CYP71B7 | cytochrome P | 0,01436092 | 1,05469161 |
| AT1G63855 | NA | Putative meth | 0,01453609 | -1,08765062 |
| AT3G48180 | na | na | 0,01457165 | -1,81735907 |
| AT5G15960 | KIN1 | stress-respons | 0,0145838 | -1,5494545 |
| AT5G61100 | na | na | 0,01460631 | -1,29410117 |
| AT1G59510 | CF9 | Carbohydrate- | 0,01476622 | -1,06805284 |
| AT1G02820 | NA | Late embryog | 0,01488604 | -2,16603905 |
| AT2G46840 | ATDUF4 | DOMAIN OF U | 0,01489016 | 1,74300972 |
| AT1G65520 | ATECI1 | delta(3) delta( | 0,01490743 | -1,06362454 |
| AT4G18220 | NA | Drug/metabol | 0,01504213 | 1,2712924 |
| AT1G31290 | AGO3 | ARGONAUTE 3 | 0,01511138 | -2,25222757 |

| | | | | |
|---|---|---|---|---|
| AT1G22600 | NA | Late embryog | 0,01525116 | 1,73689025 |
| AT4G16045 | NA | TRAF-like supe | 0,01527885 | -4,54452804 |
| AT1G59590 | ZCF37 | ZCF37 | 0,01529893 | -2,55169121 |
| AT5G56550 | ATOXS3 | oxidative stres | 0,01529893 | -1,75972325 |
| AT5G22270 | na | na | 0,01541695 | -2,22831109 |
| AT1G71691 | NA | GDSL-like Lipa | 0,01543474 | 1,37476787 |
| AT5G50130 | NA | NAD(P)-bindir | 0,01565759 | -1,22587642 |
| AT5G54470 | NA | B-box type zin | 0,01575472 | -1,2660037 |
| AT1G75040 | PR-5 | pathogenesis- | 0,01585419 | 1,51691182 |
| AT2G29740 | UGT71C2 | UDP-glucosyl | 0,01592143 | -2,36217321 |
| AT1G23020 | ATFRO3 | ferric reductic | 0,015932 | -1,06023917 |
| AT5G03495 | NA | RNA-binding ( | 0,01596128 | -1,68892887 |
| AT2G32930 | ZFN2 | zinc finger nu | 0,01607356 | -1,02576307 |
| AT2G42530 | COR15B | cold regulatec | 0,01612369 | 2,47357054 |
| AT1G25275 | na | na | 0,01617947 | 1,00134124 |
| AT1G69850 | ATNRT1:2 | nitrate transp | 0,01617947 | -1,07244766 |
| AT2G28420 | NA | Lactoylglutath | 0,01623244 | 1,21742518 |
| AT4G13330 | NA | S-adenosyl-L-r | 0,01629185 | -1,33595543 |
| AT4G29250 | NA | HXXXD-type a | 0,0162974 | -8,87008509 |
| AT5G57820 | NA | zinc ion bindir | 0,01631652 | -2,1454865 |
| AT2G28160 | ATBHLH029 | FER-like regul | 0,01631965 | 1,53361169 |
| AT1G50590 | NA | RmlC-like cupi | 0,01632229 | -1,27368274 |
| AT5G27420 | ATL31 | carbon/nitrog | 0,01635724 | -1,74619623 |
| AT3G26390 | na | na | 0,01638648 | -4,05239905 |
| AT2G28610 | PRS | Homeodomai | 0,01652451 | 1,01735198 |
| AT1G04360 | NA | RING/U-box su | 0,01657622 | -1,60292023 |
| AT3G25770 | AOC2 | allene oxide c | 0,01658178 | 1,09509622 |
| AT3G55970 | ATJRG21 | jasmonate-reg | 0,01662328 | -2,16194122 |
| AT3G17290.1 | na | na | 0,01670433 | -1,29026675 |
| AT3G12190 | na | na | 0,01678335 | -3,3631735 |
| AT1G69570 | NA | Dof-type zinc | 0,01684336 | -1,78669292 |
| AT1G69820 | GGT3 | gamma-glutar | 0,01688727 | 1,85205111 |
| AT2G16360 | NA | Ribosomal prc | 0,01693656 | -1,66904269 |

| | | | | |
|---|---|---|---|---|
| AT3G03760 | LBD20 | LOB domain-c | 0,01697307 | -10,3601205 |
| AT4G00893 | na | na | 0,01697895 | -2,03472549 |
| AT4G36350 | ATPAP25 | purple acid ph | 0,01701183 | -6,40825022 |
| AT1G70270 | na | na | 0,01716181 | -1,37470869 |
| AT2G19580 | TET2 | tetraspanin2 | 0,01737221 | -1,301647 |
| AT1G09860 | ATPUP16 | purine perme | 0,01740556 | -1,48970151 |
| AT3G26820 | NA | Esterase/lipas | 0,01747504 | -6,06802456 |
| AT1G18120 | na | na | 0,01760972 | -8,36568121 |
| AT5G21130 | NA | Late embryog | 0,01771749 | -3,33674421 |
| AT4G22690 | CYP706A1 | cytochrome P | 0,01773153 | 1,25571061 |
| AT5G10180 | AST68 | slufate transp | 0,01774825 | 1,42877501 |
| AT5G45810 | CIPK19 | CBL-interactin | 0,01780455 | -2,85763063 |
| AT4G37370 | CYP81D8 | cytochrome P | 0,01780557 | -2,64216878 |
| AT1G10300 | NA | Nucleolar GTP | 0,01783874 | -1,28598938 |
| AT5G51580 | na | na | 0,01793764 | -1,84127598 |
| AT1G35170 | NA | TRAM LAG1 a | 0,01798081 | 1,40869661 |
| AT3G19320 | NA | Leucine-rich r | 0,01799864 | -2,33363314 |
| AT3G14700 | NA | SART-1 family | 0,01803006 | -1,69584797 |
| AT5G06930 | na | na | 0,01806536 | -1,30176731 |
| AT2G46950 | CYP709B2 | cytochrome P | 0,01812344 | -1,56133412 |
| AT2G03460 | NA | Galactose oxi | 0,01813812 | -2,15975951 |
| AT1G03870 | FLA9 | FASCICLIN-like | 0,01828128 | -1,87848435 |
| AT1G01460 | ATPIPK11 | Phosphatidyli | 0,01845593 | -3,50411396 |
| AT1G78230 | NA | Outer arm dy | 0,01865004 | -2,1455363 |
| AT4G18210 | ATPUP10 | purine perme | 0,01874017 | 1,29448989 |
| AT1G71015 | na | na | 0,01875015 | -1,65606818 |
| AT5G60270 | NA | Concanavalin | 0,01879382 | -1,67611558 |
| AT3G60890 | ZPR2 | protein bindin | 0,01888275 | -1,4462056 |
| AT3G04110 | ATGLR1.1 | glutamate rec | 0,01895426 | 1,20863489 |
| AT1G05530 | UGT2 | UDP-glucosyl | 0,0190688 | -2,08124716 |
| AT1G61740 | NA | Sulfite export | 0,01921804 | -1,07566799 |
| AT4G12730 | FLA2 | FASCICLIN-like | 0,01938612 | -1,03815642 |
| AT4G39890 | AtRABH1c | RAB GTPase h | 0,01942178 | -2,02930223 |

| | | | | |
|---|---|---|---|---|
| AT5G67300 | ATMYB44 | myb domain p | 0,01952375 | -1,82986274 |
| AT3G26742 | na | na | 0,01954695 | 2,35184426 |
| AT3G51660 | NA | Tautomerase/ | 0,01954695 | 1,27215321 |
| AT1G03445 | BSU1 | Serine/threon | 0,01954695 | -2,44712298 |
| AT3G47340 | ASN1 | glutamine-dep | 0,01954695 | -1,42010821 |
| AT1G01760 | NA | adenosine dea | 0,01968952 | 1,01868264 |
| AT5G02390 | NA | Protein of unk | 0,01971648 | -2,86039739 |
| AT4G39700 | NA | Heavy metal t | 0,01976116 | -1,61210519 |
| AT4G21480 | STP12 | sugar transpo | 0,01976116 | -1,17164813 |
| AT2G41040 | NA | S-adenosyl-L-r | 0,01978069 | -2,98872381 |
| AT1G17147 | NA | VQ motif-cont | 0,01978502 | 1,76906059 |
| AT3G56710 | SIB1 | sigma factor b | 0,01983834 | 1,35696769 |
| AT5G51440 | NA | HSP20-like cha | 0,01988736 | -1,45616402 |
| AT4G29690 | NA | Alkaline-phosp | 0,01990661 | -1,33302178 |
| AT1G50310 | ATSTP9 | sugar transpo | 0,02009813 | 1,50567662 |
| AT1G72280 | AERO1 | endoplasmic r | 0,02014346 | -2,13898223 |
| AT5G48670 | AGL80 | AGAMOUS-lik | 0,02030311 | -1,66668319 |
| AT4G17030 | AT-EXPR | expansin-like | 0,02037055 | -1,78361002 |
| AT1G09500 | NA | NAD(P)-bindir | 0,02038198 | 1,7991996 |
| AT4G18080 | na | na | 0,02050149 | -1,86886948 |
| AT5G59100 | NA | Subtilisin-like | 0,02060095 | -6,67633555 |
| AT4G13340 | NA | Leucine-rich r | 0,02066626 | -1,17779914 |
| AT1G45545 | NA | Plant protein | 0,02067009 | -2,69139998 |
| AT2G01020 | na | na | 0,02076637 | 1,02075042 |
| AT1G53560 | NA | Ribosomal pro | 0,0207727 | 1,27437731 |
| AT5G13150 | ATEXO70C1 | exocyst subur | 0,02077886 | -2,05005622 |
| AT2G42885 | NA | Defensin-like | 0,0208199 | -1,2142291 |
| AT5G16570 | GLN1;4 | glutamine syn | 0,02083177 | -1,41241899 |
| AT3G24420 | NA | alpha/beta-Hy | 0,02090452 | -1,46374669 |
| AT5G16700 | NA | Glycosyl hydro | 0,02100934 | -5,29267469 |
| AT1G74540 | CYP98A8 | cytochrome P | 0,02114149 | -4,12566116 |
| AT1G74630 | NA | Tetratricopep | 0,02118 | 1,00889191 |
| AT4G03220 | NA | Protein with R | 0,02130455 | -2,28132128 |

| | | | | |
|---|---|---|---|---|
| AT4G14130 | XTH15 | xyloglucan en | 0,02131507 | -1,4387417 |
| AT4G33120 | NA | S-adenosyl-L-r | 0,02136791 | 1,60955866 |
| AT5G17340 | NA | Putative mem | 0,02142932 | -8,50941236 |
| AT3G23130 | FLO10 | C2H2 and C2H | 0,02147774 | -1,6073776 |
| AT2G05160 | NA | CCCH-type zin | 0,02155031 | -1,15517775 |
| AT5G55470 | ATNHX3 | Na+/H+ (sodiu | 0,02160048 | -1,00670175 |
| AT1G21326 | NA | VQ motif-cont | 0,02180431 | -2,68246008 |
| AT4G17483 | NA | alpha/beta-Hy | 0,02186487 | -2,1399069 |
| AT2G31980 | AtCYS2 | PHYTOCYSTAT | 0,02187029 | -1,50619549 |
| AT4G37140 | ATMES20 | alpha/beta-Hy | 0,0218746 | 1,0669316 |
| AT5G66390 | NA | Peroxidase su | 0,02193456 | -1,11678506 |
| AT5G03360 | NA | DC1 domain-c | 0,02196368 | -1,96777857 |
| AT1G61110 | anac025 | NAC domain c | 0,02203604 | -6,90387533 |
| AT2G17880 | NA | Chaperone Dr | 0,02226197 | -1,10161549 |
| AT3G60970 | ATMRP15 | multidrug resi | 0,02229393 | -2,37162337 |
| AT5G38680 | NA | Galactose oxi | 0,02230716 | -2,28832545 |
| AT2G35210 | AGD10 | root and polle | 0,02236655 | -1,72191319 |
| AT5G58310 | ATMES18 | methyl estera | 0,02243767 | -3,63726472 |
| AT5G62510 | NA | F-box family p | 0,02250047 | -2,859823 |
| AT1G18520 | TET11 | tetraspanin11 | 0,02253741 | -8,23532767 |
| AT4G30230 | na | na | 0,02256 | 1,38575892 |
| AT4G25420 | AT2301 | 2-oxoglutarat | 0,02288635 | 1,0572318 |
| AT5G49780 | NA | Leucine-rich r | 0,02293945 | 1,4677121 |
| AT5G66690 | UGT72E2 | UDP-Glycosylt | 0,02300443 | -3,5332683 |
| AT2G29500 | NA | HSP20-like ch | 0,02306709 | -1,04326646 |
| AT2G21237 | na | na | 0,02311175 | -3,11961987 |
| AT4G17800 | NA | Predicted AT-l | 0,02317397 | 1,00684686 |
| AT1G61550 | NA | S-locus lectin | 0,02323842 | 2,65048283 |
| AT4G27440 | PORB | protochloroph | 0,02333648 | 1,71113652 |
| AT3G05152 | na | na | 0,02337746 | -1,77335361 |
| AT1G62975 | NA | basic helix-loc | 0,02356472 | 1,48843974 |
| AT1G24140 | NA | Matrixin famil | 0,02373447 | -2,29915951 |
| AT4G19810 | NA | Glycosyl hydr | 0,02397226 | -2,76853072 |

| | | | | |
|---|---|---|---|---|
| AT5G10280 | ATMYB64 | myb domain p | 0,02417825 | -1,28825411 |
| AT1G48400 | NA | F-box/RNI-like | 0,02432274 | -1,82129069 |
| AT1G49832 | na | na | 0,02436034 | 1,25472406 |
| AT1G45100 | NA | RNA-binding ( | 0,02436442 | -6,44781271 |
| AT2G27500 | NA | Glycosyl hydro | 0,02438821 | -1,3825679 |
| AT4G11720 | GCS1 | hapless 2 | 0,0245014 | -1,45234793 |
| AT1G03935 | na | na | 0,02458085 | -1,37832421 |
| AT1G15460 | ATBOR4 | HCO3- transpo | 0,02474239 | -7,18974691 |
| AT5G48400 | ATGLR1.2 | Glutamate rec | 0,02506801 | -2,1193638 |
| AT4G04760 | NA | Major facilitat | 0,02518596 | -4,00967967 |
| AT5G58660 | NA | 2-oxoglutarate | 0,02530452 | -1,8296281 |
| AT4G13611 | na | na | 0,02538903 | -2,61170242 |
| AT2G18700 | ATTPS11 | trehalose pho | 0,025416 | -1,18657697 |
| AT2G32800 | AP4.3A | protein kinase | 0,02549378 | -1,31236507 |
| AT3G44900 | ATCHX4 | cation/H+ exc | 0,02560435 | 1,0588013 |
| AT2G21130 | NA | Cyclophilin-lik | 0,02607157 | 1,60442003 |
| AT1G75166 | na | na | 0,0261807 | -1,12435562 |
| AT4G09012 | NA | Mitochondrial | 0,02620355 | -1,43696762 |
| AT2G38152 | NA | alpha 14-glyco | 0,02627418 | -1,73399886 |
| AT3G27250 | na | na | 0,02631334 | -1,72670441 |
| AT3G50740 | UGT72E1 | UDP-glucosyl | 0,02656621 | 1,23339985 |
| AT5G66020 | ATSAC1B | Phosphoinosit | 0,02662229 | -3,79688215 |
| AT1G62370 | NA | RING/U-box su | 0,02681368 | -1,03533765 |
| AT1G69970 | CLE26 | CLAVATA3/ES | 0,02687559 | -1,20596293 |
| AT1G62510 | NA | Bifunctional in | 0,02697821 | -3,8348316 |
| AT2G37670 | NA | Transducin/W | 0,02697821 | -1,29874384 |
| AT1G06320 | na | na | 0,02698338 | -1,21405765 |
| AT2G39920 | NA | HAD superfam | 0,02708112 | 2,11324671 |
| AT3G22620 | NA | Bifunctional in | 0,02714247 | -1,46848752 |
| AT1G13970 | NA | Protein of unk | 0,02714247 | -1,40195455 |
| AT5G11990 | NA | proline-rich fa | 0,02770859 | -1,29798131 |
| AT4G18550 | NA | alpha/beta-Hy | 0,02778627 | -3,67777307 |
| AT5G59820 | RHL41 | C2H2-type zin | 0,0278334 | -1,99826096 |

| | | | | |
|---|---|---|---|---|
| AT1G35140 | EXL7 | Phosphate-res | 0,02802964 | -4,47694617 |
| AT1G53880 | NA | Eukaryotic tra | 0,0280548 | -1,77661992 |
| AT1G53900 | NA | Eukaryotic tra | 0,0280548 | -1,77661992 |
| AT5G20960 | AAO1 | aldehyde oxid | 0,02809432 | 1,36093206 |
| AT4G29740 | ATCKX4 | cytokinin oxid | 0,02826736 | 1,46467901 |
| AT5G13380 | NA | Auxin-respons | 0,02827219 | -7,86637003 |
| AT3G02310 | AGL4 | K-box region a | 0,02828433 | -4,01733526 |
| AT2G32150 | NA | Haloacid deha | 0,02831379 | -1,01314315 |
| AT2G35700 | ATERF38 | ERF family pro | 0,02849476 | 1,06410553 |
| AT1G14250 | NA | GDA1/CD39 n | 0,02857984 | 2,7504438 |
| AT1G08860 | BON3 | Calcium-depe | 0,02861303 | -1,43578613 |
| AT4G39364 | na | na | 0,0286815 | 1,14785525 |
| AT1G72125 | NA | Major facilitat | 0,02876417 | 1,10597937 |
| AT1G29050 | TBL38 | TRICHOME BII | 0,02897699 | -1,11178624 |
| AT5G41315 | GL3 | basic helix-loo | 0,02909681 | 1,5884661 |
| AT4G15100 | scpl30 | serine carboxy | 0,02913456 | -4,92633773 |
| AT3G13600 | NA | calmodulin-bi | 0,02913456 | -1,32579934 |
| AT2G31620 | NA | Receptor-like | 0,02915588 | -3,02013137 |
| AT2G19070 | SHT | spermidine hy | 0,02916882 | -10,5856346 |
| AT2G20700 | LLG2 | LORELEI-LIKE- | 0,02919067 | -1,53048627 |
| AT5G41750 | NA | Disease resista | 0,02953665 | -1,02238824 |
| AT4G01950 | ATGPAT3 | glycerol-3-pho | 0,02954031 | 1,87973946 |
| AT5G59120 | ATSBT4.13 | subtilase 4.13 | 0,0295923 | -3,02859349 |
| AT3G02480 | NA | Late embryog | 0,02959667 | 1,19131945 |
| AT5G65870 | ATPSK5 | phytosulfokin | 0,02967323 | -1,13388571 |
| AT5G06740 | NA | Concanavalin | 0,02990966 | -1,89537393 |
| AT5G54060 | UF3GT | UDP-glucose:f | 0,03013469 | 1,23501885 |
| AT5G09730 | ATBX3 | beta-xylosidas | 0,03014231 | -1,26066617 |
| AT3G26010 | NA | Galactose oxic | 0,03020505 | -1,93364226 |
| AT2G14475 | na | na | 0,03025547 | -1,72225991 |
| AT2G35290 | na | na | 0,03033593 | -1,16760808 |
| AT4G33565 | NA | RING/U-box su | 0,03066458 | -1,12820372 |
| AT2G24850 | TAT | tyrosine amin | 0,03070796 | 1,68530654 |

| | | | | |
|---|---|---|---|---|
| AT2G16260 | na | na | 0,03097065 | -1,8529974 |
| AT3G21800 | UGT71B8 | UDP-glucosyl | 0,03101786 | -2,78875996 |
| AT2G01890 | ATPAP8 | purple acid ph | 0,03113032 | -1,1832937 |
| AT1G54280 | NA | ATPase E1-E2 | 0,03117631 | -1,28974282 |
| AT3G62740 | BGLU7 | beta glucosida | 0,03135754 | 1,13004575 |
| AT2G22960 | NA | alpha/beta-Hy | 0,03138709 | -1,17215843 |
| AT3G19880 | NA | F-box and ass | 0,03146263 | -6,7133289 |
| AT3G58860 | NA | F-box/RNI-like | 0,03150748 | -2,10518225 |
| AT5G60500 | NA | Undecaprenyl | 0,03162907 | -10,6325839 |
| AT4G26540 | NA | Leucine-rich r | 0,03177232 | 1,18913356 |
| AT5G40780 | LHT1 | lysine histidin | 0,03183829 | -1,3015477 |
| AT5G58630 | na | na | 0,03190456 | 1,0541823 |
| AT1G66810 | NA | Zinc finger C-x | 0,03190456 | -1,16574414 |
| AT1G67990 | ATTSM1 | S-adenosyl-L-r | 0,0319056 | -9,14773944 |
| AT2G40000 | ATHSPRO2 | ortholog of su | 0,0319056 | -1,25557109 |
| AT1G23540 | AtPERK12 | Protein kinase | 0,03192224 | -2,10978517 |
| AT4G10260 | NA | pfkB-like carb | 0,03241584 | 1,3304106 |
| AT3G50390 | NA | Transducin/W | 0,03273091 | 1,38841382 |
| AT5G07600 | NA | Oleosin family | 0,03347974 | -6,94813967 |
| AT3G29639 | na | na | 0,03352283 | 1,72551312 |
| AT5G13990 | ATEXO70C2 | exocyst subur | 0,03358911 | -1,77613368 |
| AT1G24320 | NA | Six-hairpin gly | 0,03374258 | -1,08896034 |
| AT3G09790 | UBQ8 | ubiquitin 8 | 0,03382915 | -1,51029019 |
| AT2G23970 | NA | Class I glutam | 0,03385468 | -2,60846314 |
| AT3G06145 | na | na | 0,03401918 | -1,00944681 |
| AT5G25750 | na | na | 0,03416875 | 2,79191227 |
| AT4G20040 | NA | Pectin lyase-li | 0,03428984 | -1,09307928 |
| AT3G55646 | na | na | 0,03440879 | -3,07401839 |
| AT5G16960 | NA | Zinc-binding d | 0,03453696 | -9,77795749 |
| AT3G51478 | na | na | 0,03475679 | 1,08228148 |
| AT5G04400 | anac077 | NAC domain c | 0,03487351 | -1,88233214 |
| AT3G51590 | LTP12 | lipid transfer p | 0,03498369 | -10,1607923 |
| AT2G29860 | NA | Galactose oxi | 0,03498369 | -6,23656726 |

| | | | | |
|---|---|---|---|---|
| AT1G05000 | NA | Phosphotyros | 0,0350344 | -1,44012821 |
| AT1G65620 | AS2 | Lateral organ | 0,0350344 | -1,33539907 |
| AT1G23060 | na | na | 0,03504672 | -3,63676925 |
| AT4G28040 | NA | nodulin MtN2 | 0,03506774 | -3,81789767 |
| AT4G20770 | NA | Pentatricopep | 0,03513857 | 1,16825197 |
| AT5G25310 | NA | Exostosin fam | 0,03514815 | -1,85486259 |
| AT3G47130 | NA | F-box associat | 0,03516584 | -2,76030139 |
| AT1G55330 | AGP21 | arabinogalact | 0,03539021 | -1,03189197 |
| AT3G51400 | NA | Arabidopsis pr | 0,03541174 | 3,04611025 |
| AT3G48740 | NA | Nodulin MtN3 | 0,03561752 | -1,0858237 |
| AT4G04450 | AtWRKY42 | WRKY family t | 0,03575377 | 1,04629793 |
| AT5G43440 | NA | 2-oxoglutarate | 0,03594181 | -2,49514895 |
| AT2G37280 | ATPDR5 | pleiotropic dr | 0,03611456 | -1,2143686 |
| AT3G04070 | anac047 | NAC domain c | 0,03652074 | -2,51317516 |
| AT3G55890 | NA | Yippee family | 0,03653602 | -2,03790958 |
| AT4G21690 | ATGA3OX3 | gibberellin 3-c | 0,03658943 | 1,67447061 |
| AT1G43765 | na | na | 0,03658943 | 1,04516054 |
| AT1G60360 | NA | RING/U-box su | 0,03665589 | -1,29556693 |
| AT2G07777 | NA | ATP synthase | 0,03677632 | -1,2086562 |
| AT3G62150 | PGP21 | P-glycoprotein | 0,03713911 | 1,26378448 |
| AT1G69900 | NA | Actin cross-lin | 0,03726625 | 1,39621673 |
| AT3G61730 | RMF | reduced male | 0,03734277 | -1,182849 |
| AT4G00695 | na | na | 0,0375367 | -1,15393231 |
| AT4G38400 | ATEXLA2 | expansin-like | 0,03755744 | -1,18427651 |
| AT4G26670 | NA | Mitochondrial | 0,03764927 | 1,20200376 |
| AT4G15070 | NA | Cysteine/Histi | 0,03778338 | 1,06354461 |
| AT2G41180 | NA | VQ motif-cont | 0,03804237 | 1,08463058 |
| AT1G79170 | na | na | 0,03804237 | -1,43786497 |
| AT2G23800 | GGPS2 | geranylgerany | 0,03822703 | -5,5720859 |
| AT5G60510 | NA | Undecaprenyl | 0,03823755 | -12,2010826 |
| AT1G35190 | NA | 2-oxoglutarate | 0,03830517 | -1,00365874 |
| AT4G01680 | AtMYB55 | myb domain p | 0,03836077 | -1,25195743 |
| AT3G19680 | NA | Protein of unk | 0,03856774 | -1,63576677 |

| | | | | |
|---|---|---|---|---|
| AT5G40630 | NA | Ubiquitin-like | 0,03869574 | 1,28633801 |
| AT4G15093 | NA | catalytic LigB s | 0,0391015 | -1,02670908 |
| AT1G20120 | NA | GDSL-like Lipa | 0,03919301 | -5,33533377 |
| AT1G16730 | UP6 | unknown prot | 0,03920864 | -1,13480274 |
| AT5G42830 | NA | HXXXD-type a | 0,03931718 | -1,08217665 |
| AT2G18130 | ATPAP11 | purple acid ph | 0,03947104 | -10,2460901 |
| AT3G10600 | CAT7 | cationic amino | 0,03984168 | -1,75901785 |
| AT1G65490 | na | na | 0,03985441 | -1,5061968 |
| AT5G49700 | NA | Predicted AT-l | 0,03990258 | 1,1771115 |
| AT3G25880 | NA | NAD(P)-bindir | 0,04028641 | 1,47027656 |
| AT1G65240 | NA | Eukaryotic asp | 0,04037386 | -1,1948976 |
| AT5G28320 | na | na | 0,04038781 | -1,18964358 |
| AT1G21510 | na | na | 0,040582 | 1,05992938 |
| AT5G58390 | NA | Peroxidase su | 0,04085389 | -2,08067626 |
| AT5G19190 | na | na | 0,04097452 | -1,75727951 |
| AT3G48390 | NA | MA3 domain- | 0,04104455 | -2,39551809 |
| AT1G66230 | AtMYB20 | myb domain p | 0,04130994 | -1,39132428 |
| AT3G11680 | NA | Aluminium ac | 0,04134917 | -1,32307763 |
| AT5G39020 | NA | Malectin/rece | 0,04143471 | -1,5507957 |
| AT2G43440 | NA | F-box and ass | 0,04146651 | -1,31769131 |
| AT2G34370 | NA | Pentatricopep | 0,04153404 | -2,47463936 |
| AT1G21620 | APUM20 | pumilio 20 | 0,04153404 | -1,94954403 |
| AT2G05380 | GRP3S | glycine-rich pr | 0,04166133 | -1,0271543 |
| AT4G13320 | na | na | 0,04192389 | -1,41829265 |
| AT2G21220 | NA | SAUR-like aux | 0,04211897 | -2,65996917 |
| AT1G54300 | na | na | 0,04261262 | -1,56762516 |
| AT4G33970 | NA | Leucine-rich r | 0,04276984 | -1,80265231 |
| AT1G72240 | na | na | 0,04317472 | -1,42681679 |
| AT3G63160 | na | na | 0,0434333 | 1,11319737 |
| AT1G62490 | NA | Mitochondrial | 0,04358934 | 1,48525509 |
| AT3G28330 | NA | F-box family p | 0,04429248 | -2,91760109 |
| AT4G25430 | na | na | 0,04442678 | 1,13253102 |
| AT4G36040 | NA | Chaperone Dr | 0,04442678 | -1,11713498 |

| | | | | |
|---|---|---|---|---|
| AT3G21660 | NA | UBX domain-c | 0,04449272 | -1,45849192 |
| AT5G07500 | PEI1 | Zinc finger C-x | 0,04468675 | -1,30208644 |
| AT1G75490 | NA | Integrase-type | 0,04470253 | -1,71540232 |
| AT1G28370 | ATERF11 | ERF domain pr | 0,04486504 | 2,3237176 |
| AT1G22440 | NA | Zinc-binding a | 0,0451219 | -2,04123098 |
| AT5G60330 | na | na | 0,04525158 | -1,7030443 |
| AT1G10220 | na | na | 0,04525158 | -1,62622246 |
| AT5G25450 | NA | Cytochrome b | 0,04610981 | -1,55915286 |
| AT3G15240 | NA | Serine/threon | 0,04619535 | -2,34205154 |
| AT5G58460 | ATCHX25 | cation/H+ exc | 0,04651184 | -1,68578031 |
| AT1G27110 | NA | Tetratricopept | 0,04725865 | -1,16644733 |
| AT3G13850 | LBD22 | LOB domain-c | 0,04798785 | -1,57055797 |
| AT5G49770 | NA | Leucine-rich re | 0,04810498 | -1,6687542 |
| AT5G15940 | NA | NAD(P)-bindir | 0,04826907 | -1,01008997 |
| AT4G29610 | NA | Cytidine/deox | 0,04827007 | 1,44539835 |
| AT1G30940 | na | na | 0,04881942 | -2,90477439 |
| AT1G69490 | ANAC029 | NAC-like activ | 0,04913153 | -1,62839791 |
