## supplemental figures for "SEPALLATA MADS transcription factors act as key regulators in fertilization efficiency, ovule outer integument growth and mucilage secretory cell differentiation in Arabidopsis"

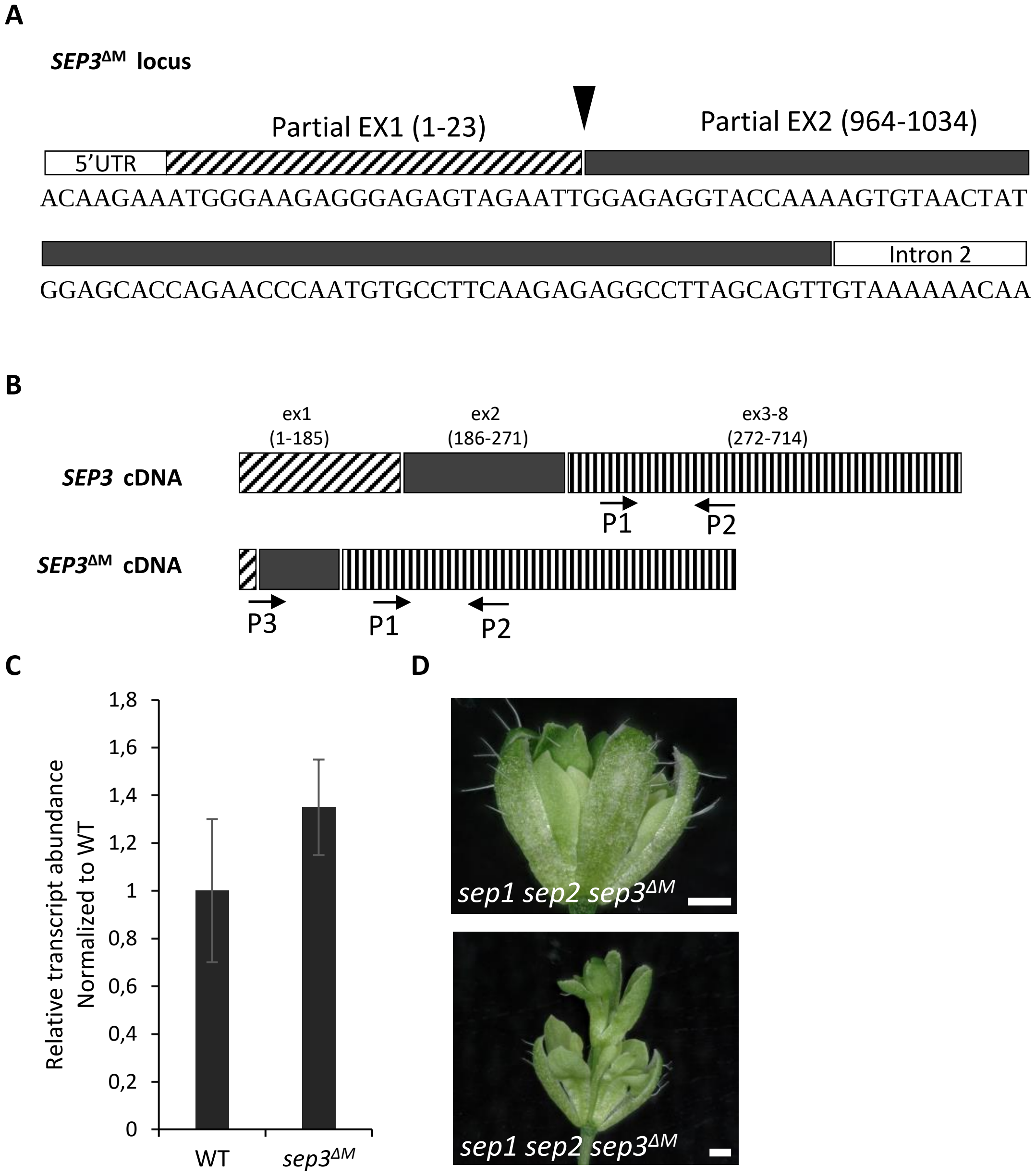


**Supplementary Figure S1. Validation of *SEP3*$^{\Delta M}$ expression in *sep3*$^{\Delta M}$**

**(A)** Sequence surrounding the Cas9 deletion at the *SEP3*$^{\Delta M}$ locus. Arrow indicates the site of deletion. The hatched and filled rectangles define the sequences left from the first (bp 1 to 23) and second exon (ex; bp 202 to 271) after deletion, respectively. Deletion did not lead to frame shift. Numbering refers to WT genomic sequence starting at 1 with ATG. **(B)** Coding DNA derived from *SEP3 and SEP3*$^{\Delta M}$ loci with position of primers P1, P2 and P3 used for qPCR analysis in C. Numbering refers to WT *SEP3* cDNA sequence starting at 1 from the ATG. **(C)** Relative quantification of *SEP3*$^{\Delta M}$ expression in the mutant compared to *SEP3* in the WT. qPCR performed with P2, P3 combination amplifies specifically *SEP3*$^{\Delta M}$ cDNA, while combination of P1 and P3 amplify *SEP3* cDNA in WT and *SEP3*$^{\Delta M}$ cDNA in the mutant and allow to calculate the relative abundance of *SEP3*$^{\Delta M}$ in the mutant compared to *SEP3* in the WT. *SEP3* expression in WT expression was set to one. Data correspond to the mean of 2 independent biological replicates ± SD. **(D)** Introduced by cross into the *sep1 sep2* mutant, *sep3*$^{\Delta M}$ lead to loss of organ identity and loss of flower determinacy highlighting the loss of *SEP3* function. Scale indicates 500µm.

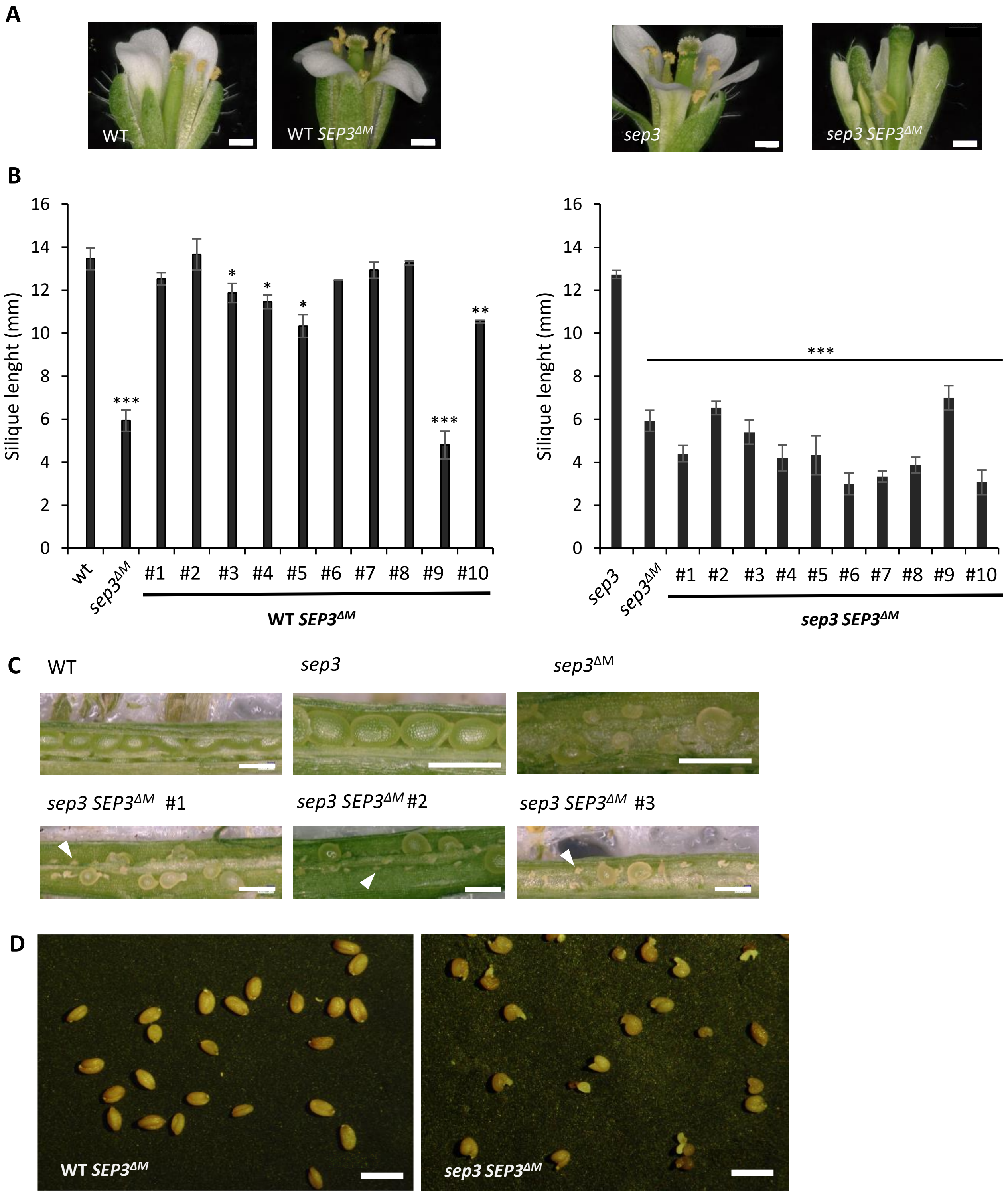


**Supplementary Figure S2: Floral organs, silique development and seed morphology in T1 WT and *sep3* plants, expressing *SEP3*$^{\Delta M}$ as transgene.**

**(A)** Representative flower development of WT and *sep3* expressing *SEP3*$^{\Delta M}$ (WT *SEP3*$^{\Delta M}$ , *sep3 SEP3*$^{\Delta M}$ ). **(B)** Silique length measured on 10 WT and *sep3* T1 plant expressing *SEP3*$^{\Delta M}$. Data represent the mean ± SEM. **(C)** Pictures of opened silique in WT, *sep3*, *sep3*$^{\Delta M}$ and 3 independent T1 *sep3* plant expressing *SEP3*$^{\Delta M}$ . The large number of unfertilized ovules (*) observed in *sep3* expressing *SEP3*$^{\Delta M}$ (*sep3 SEP3*$^{\Delta M}$) indicate that expression of *SEP3*$^{\Delta M}$ reduced fertilization as observed in *sep3*$^{\Delta M}$ . **(D)** Representative seed morphology in T1WT and *sep3* plants expressing *SEP3*$^{\Delta M}$ as transgene (*sep3 SEP3*$^{\Delta M}$). Abnormal seed shape is observed in *sep3*$^{\Delta M}$ and *sep3 SEP3*$^{\Delta M}$. Asterisks indicate significant differences from WT (left graph) or *sep3* (right graph) (* $P < 0.05$, **$P < 0.01$, ***$P < 0.001$). Scale:500µm.

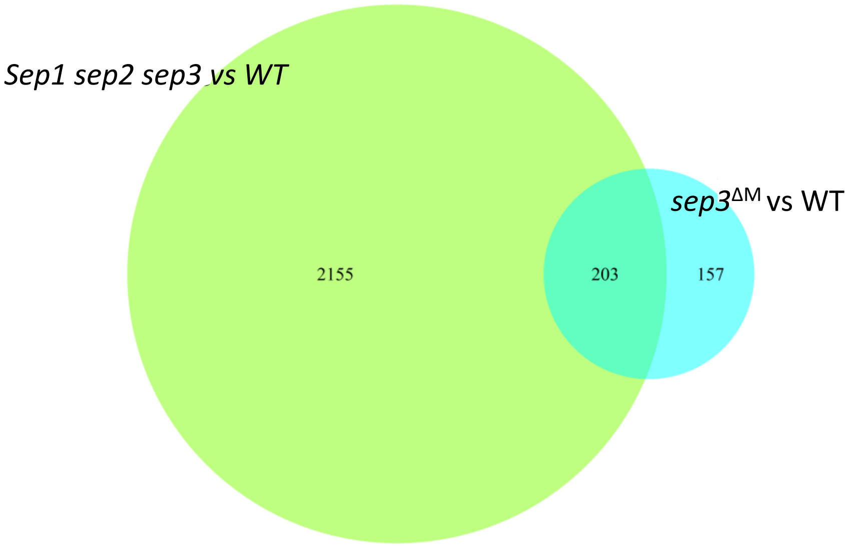


**Supplementary Figure S3. Genes deregulated in *sep3*$^{\Delta M}$ inflorescence containing stage 10/11 flower buds.**

Venn diagrams for deregulated genes in *sep1 sep2 sep3* and *sep3*$^{\Delta M}$ mutants versus WT. RNA-seq were performed in WT Col-0 and *sep3*$^{\Delta M}$ grown in parallel (This work) and compared to previously performed RNA-seq in WT Col-0 and *sep1 sep2 sep3* (Lai et al., NAR 2020).

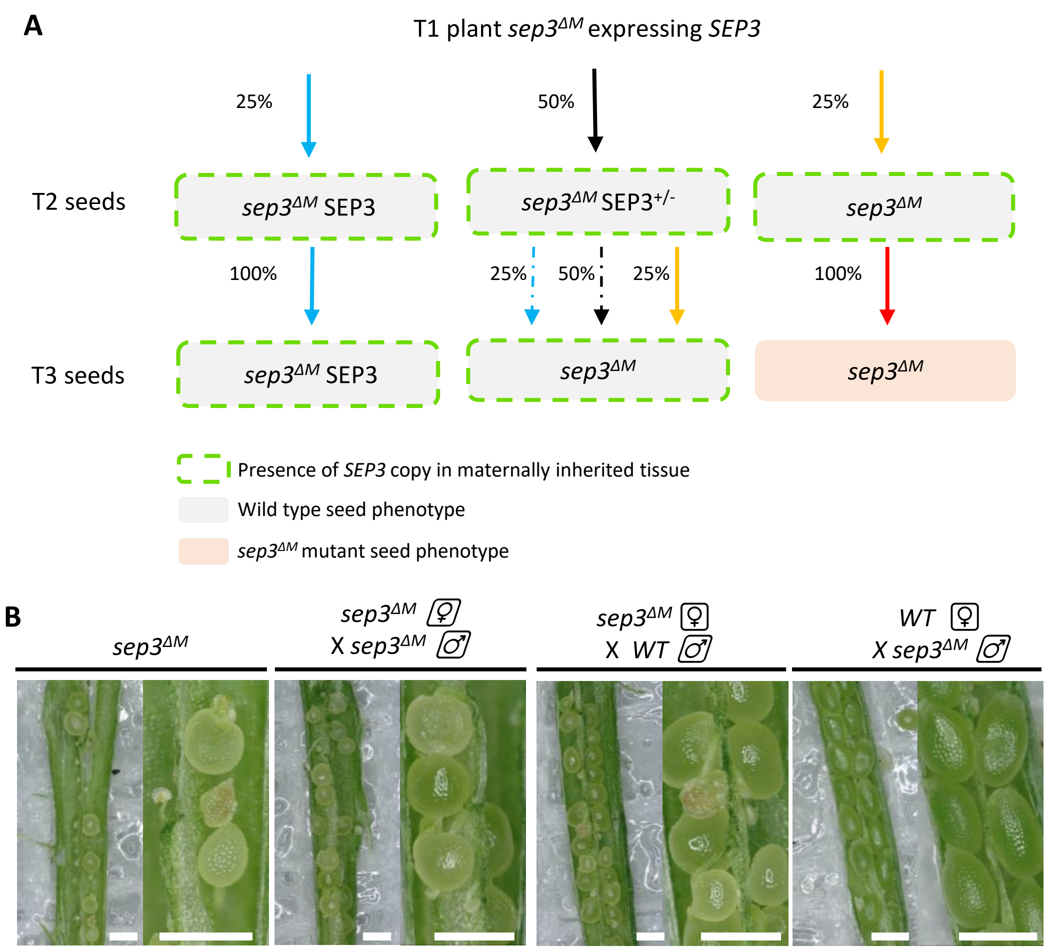


**Supplementary Figure S4. Seed phenotype in *sep3$^{ΔM}$* shows sporophytic origin.**
**(A)** Schematic representation of complementation analysis of *sep3$^{ΔM}$* with *SEP3*, showing that seed phenotype has a sporophytic origin. Seeds containing *SEP3* as transgene in the maternally inherited tissue show WT phenotype even if the gametophyte is homozygous for the *sep3$^{ΔM}$* mutation. The different color of arrow indicate different set of genotypes (*Blue*: homozygous *sep3$^{ΔM}$* expressing two copies of *SEP3* in the gametophyte; *Black*: homozygous *sep3$^{ΔM}$* expressing one copy of *SEP3* in the gametophyte*; Yellow*: homozygous *sep3$^{ΔM}$* with no copy of *SEP3* in the gametophyte, but with SEP3 in the integument; *Red*: homozygous *sep3$^{ΔM}$* with no copy of *SEP3* in the gametophyte, or in the integument. Genotypes are not represented when the lines are dashed. **(B)** *Left panel*: Opened silique of *sep3$^{ΔM}$* in condition of self-pollination highlighting reduced ovule fertilization, seed abortion and malformed seed. *Medium panels*: Opened silique of *sep3$^{ΔM}$* after hand pollination *with sep3$^{ΔM}$* or WT pollen. Both crosses triggers seeds phenotype highlighting its sporophytic origin. Hand pollination enhances number of fertilized ovule and developing seeds in the mutant but still several ovules are unfertilized. *Right panel*: indicates that mutant pollen does not trigger abnormal seed shape after fertilization of WT ovules. Scale bar is 500 µm.

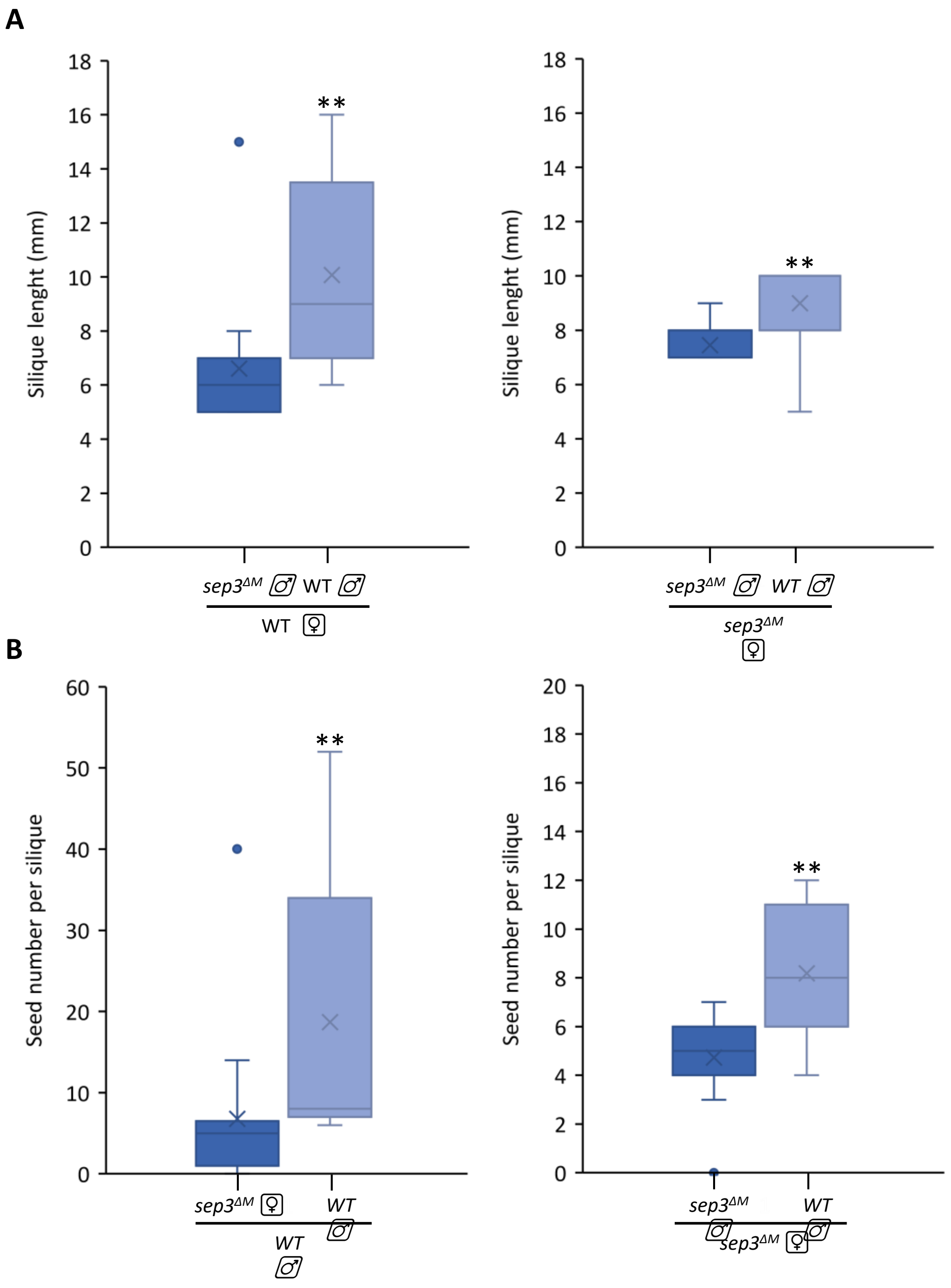


**Supplementary Figure S5. Comparative pollen fitness in WT and *sep3*$^{\Delta M}$ mutants.**
Silique elongation **(A)** and seed number per silique **(B)** are reduced when mutant pollen was used to pollinate WT carpel (left panels), highlighting a reduced pollen fitness in *sep3*$^{\Delta M}$ compared to WT pollen. Similar results were observed when *sep3*$^{\Delta M}$ was backcrossed with WT and mutant pollen (right panels). Thirteen crosses were performed in each direction. Asterisks indicate significant differences from mutant pollen (* $P < 0.05$, **$P < 0.01$, ***$P < 0.001$).

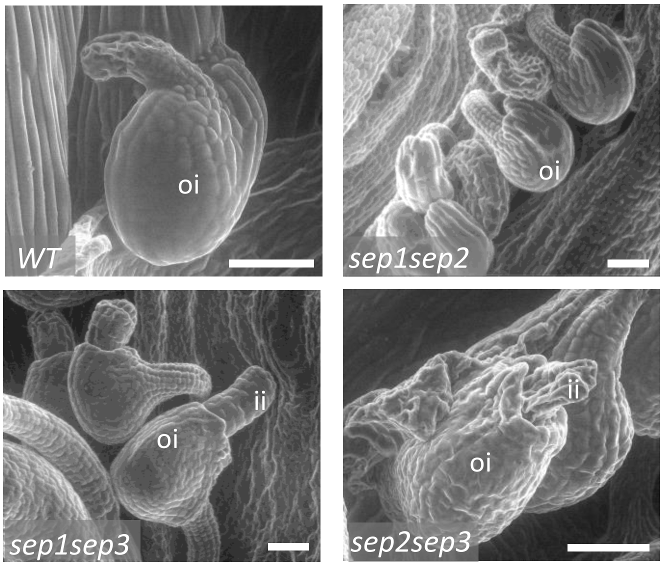


**Supplementary Figure S6**. ***SEP* titration in *Arabidopsis thaliana* ovule development.**
Electron microscopy of mature ovules in double *sep* mutant highlight that *SEP3* is required for correct growth of the outer integument. In WT and *sep1 sep2* ovules, outer integument covers the inner integument while inner integument is visible in *sep1 sep3* and *sep2 sep3*. *ii* : inner integument, *oi*: outer integument. Scale bar is 50 µm .

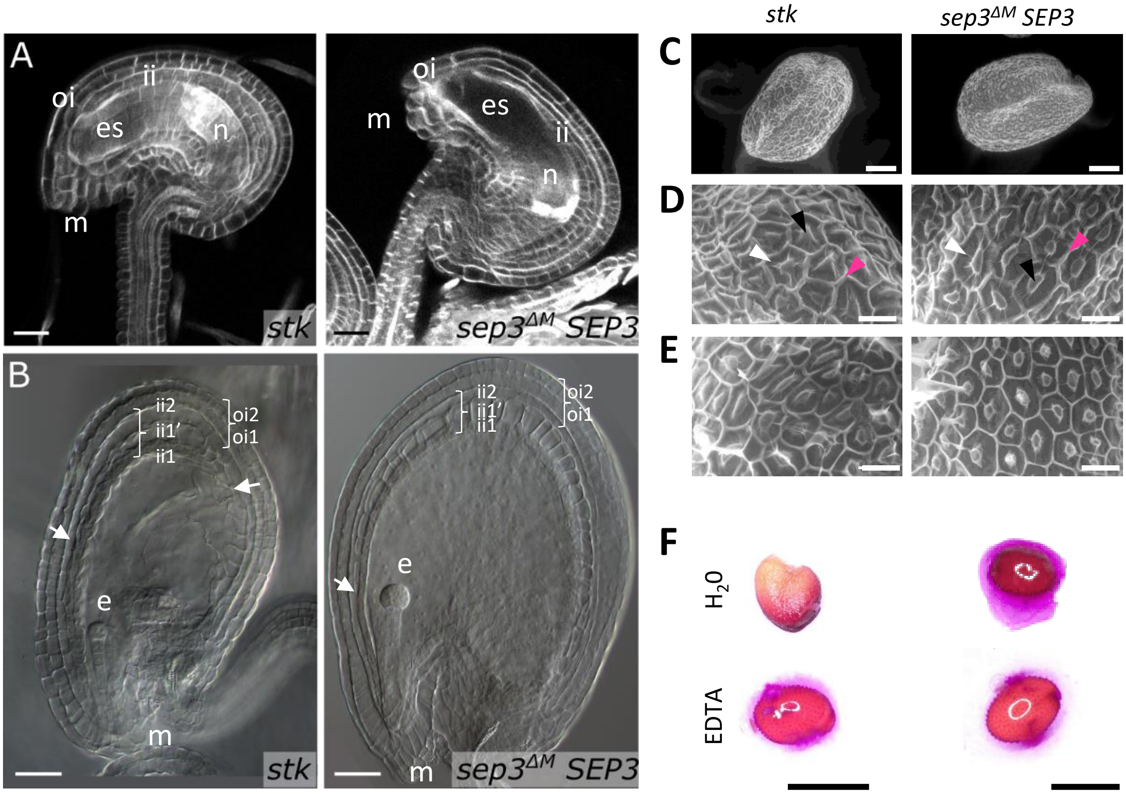


**Supplementary Figure S7**: **Integument growth, seed coat and mucilage secretory cells differentiation in *stk* and s*ep3$^{\Delta M}$* expressing *SEP3*.**

**(A)** Confocal analysis of *stk* (n=6) and **s***ep3$^{\Delta M}$* ovules expressing *SEP3* (n=5), extracted from carpel at anthesis, show ovules similar as WT in both genotypes. **(B)** Seed clearing 3 days after hand pollination of *stk* (n=6) and **s***ep3$^{\Delta M}$* ovules expressing *SEP3* (n=9). The 5 layers of the seed coat are well differentiated in both genotypes. Scanning electron microscopy of *stk* and complemented s*ep3$^{\Delta M}$* performed on whole dried seeds **(C)**, dried cell surface **(D)** and after ON water imbibition **(E)** highlight the lack of mucilage release in *stk* mutants. A minimum of three seeds were observed. **(F)** Mucilage staining with ruthenium red after overnight imbibition and EDTA treatment in the various seeds genotypes. The presence of the red allow after EDTA treatment confirms the production of mucilage in *stk*. *ii*: inner integument, *oi*: outer integument, es: embryo sac, *n*: nucellus, *m*: micropyle, *e*: embryo. Black arrow indicate columella, pink arrow, radial cell wall and white arrow, mucilage. Scales: 20 µm in A and 40 µm in B, 100 µm in C, 300µm in D and E, and 500 µm in F.

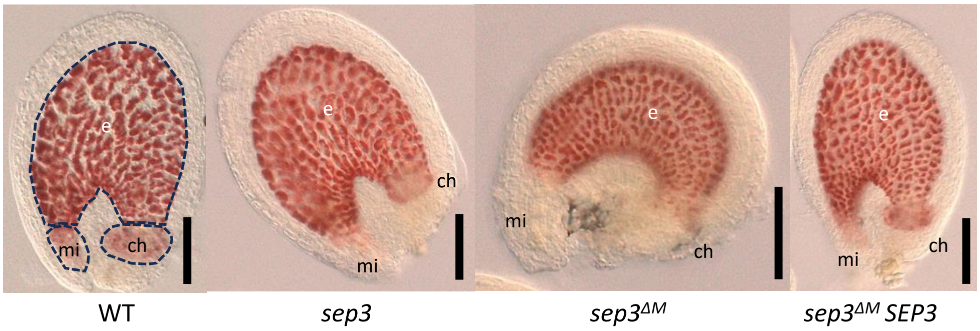


**Supplementary Figure S8. PAs accumulation in *Arabidopsis* seed 3 DAP.** Whole-mount vanillin staining confirmed the presence of PAs in the WT endothelium, the micropyle and chalazal regions. PAs accumulated in the endothelium of *sep3* and s*ep3$^{\Delta M}$* but was not visible in the chalazal and micropylar regions. WT accumulation was restored back to WT in s*ep3$^{\Delta M}$* *expressing SEP3* (*sep3$^{\Delta M}$ SEP3*). Dash lanes highlighted the endothelium, the micropyle and the chalazal regions. *e*: endothelium, *mi*: micropyle, *ch*: chalazal, *DAP*: day after pollination. Scale bar: 100 µm.

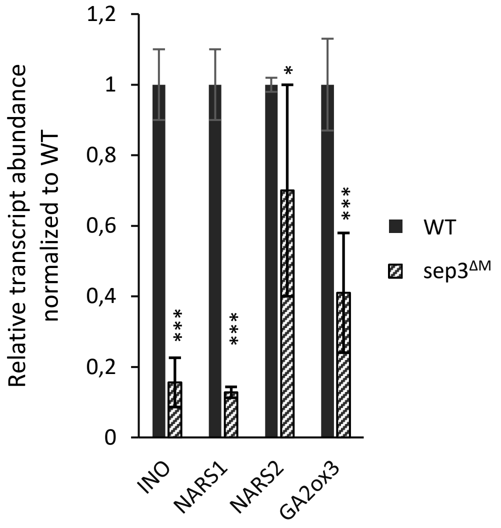


**Supplementary Figure S9. Relative quantification of *INO*, *NARS1*, *NARS2* and GA2ox3 expression in *sep3$^{\Delta M}$* mutant compared to WT.**
qPCR were performed on cDNA synthetized from RNA used in the RNA-seq experiments. The WT expression was set up to 1. Expression was normalized to *ACTIN2* and *EF-1α*. Asterisks indicate significant differences from WT (* $P < 0.05$, **$P < 0.01$, ***$P < 0.001$).

### Figure S10

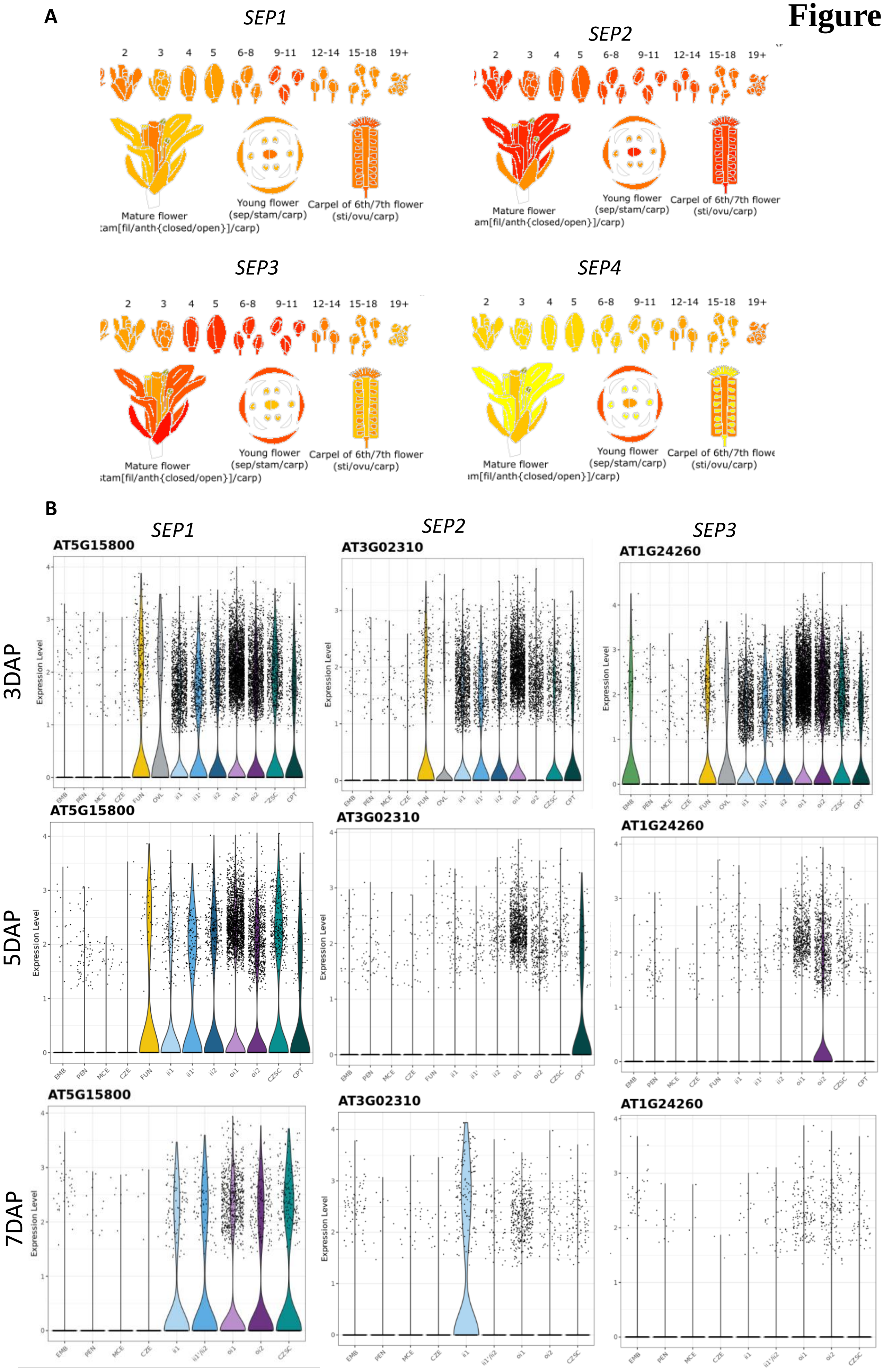
A
SEP1
SEP2
SEP3
SEP4
2 3 4 5 6-8 9-11 12-14 15-18 19+
Mature flower
Young flower (sep/stam/carp)
Carpel of 6th/7th flower (sti/ovu/carp)
B
SEP1
SEP2
SEP3
3DAP
5DAP
7DAP
AT5G15800
AT3G02310
AT1G24260
Expression Level
EMB PEN MCE CZE FUN OVL ii1 ii1' ii2 oi1 oi2 CZSC CPT
EMB PEN MCE CZE ii1 ii1'/ii2 oi1 oi2 CZSC

**Supplementary Figure S10.** ***SEP*** **expression patterns**

**(A)** Expression pattern for each member of the *SEP* family is shown for floral organs based on TAIR database (Huala et al. 2001). From yellow to red indicates increasing expression. **(B)** Expression pattern of *SEP1*, *SEP2* and *SEP3* family in ovules and during seed development after 3, 5 and 7 DAP using the snRNA Atlas data base (Martin et al. 2026). Emb: embryo, PEN: peripheral endosperm, MCE: micropylar endosperm, CZE: chalazal endosperm, FUN: funiculus, OVL: ovule, CZSC: chalazal seed coat, CPT: chalazal proliferating tissus.
